# Discovery of ICOS-targeted small molecules using affinity selection mass spectrometry screening

**DOI:** 10.1101/2024.08.04.606538

**Authors:** Longfei Zhang, Laura Calvo-Barreiro, Victor de Sousa Batista, Katarzyna Świderek, Moustafa T. Gabr

## Abstract

Inducible T cell co-stimulator (ICOS) is a positive immune checkpoint receptor expressed on the surface of activated T cells, which could promote cell function after being stimulated with ICOS ligand (ICOS-L). Although clinical benefits have been reported in the ICOS modulation-based treatment for cancer and autoimmune disease, current modulators are restricted in biologics, whereas ICOS-targeted small molecules are lacking. To fill this gap, we performed an affinity selection mass spectrometry (ASMS) screening for ICOS binding using a library of 15,600 molecules. To the best of our knowledge, this is the first study that utilizes ASMS screening to discover small molecules targeting immune checkpoints. Compound **9** with a promising ICOS/ICOS-L inhibitory profile (IC_50_ = 29.38 ± 3.41 µM) was selected as the template for the modification. Following preliminary structure-activity relationship (SAR) study and molecular dynamic (MD) simulation revealed the critical role of the ortho-hydroxy group on compound **9** in the ICOS binding, as it could stabilize the interaction via the hydrogen bond formation with residuals on the glycan, and the depletion could lead to an activity lost. This work validates a promising inhibitor for the ICOS/ICOS-L interaction, and we anticipate future modifications could provide more potent modulators for this interaction.

## Introduction

Immune checkpoints play an important role in modulating the immune system, which could promote immune cell differentiation, suppress overactive T effector cells, and maintain immunologic homeostasis. However, the dysfunction in immune checkpoints could also lead to various diseases, such as cancer and autoimmune diseases. Based on the different functions, immune checkpoint receptors could be classified into positive and negative receptors, expressed on the surface of lymphocytes and antigen-presenting cells (APCs), and stimulate or inhibit immune response upon binding with corresponding ligands via certain pathways, respectively. Inducible T cell co-stimulator (ICOS), a member of the CD28 superfamily, is a positive immune checkpoint receptor and expresses on activated T cells, which could regulate effector and memory T cell responses and maintenance^[1]^ after binding with ICOS ligand (ICOS-L) on APCs.^[2]^ Interestingly, ICOS-L could also be found on the surface of cancer cells^[3]^, which could recruit and activate the regulatory T (Treg) cells, and thus hamper the anti-tumor immune response via direct cell-cell interaction or cytokines secretion. Due to the dual outcome of the ICOS/ICOS-L pathway on effector and Treg cells, targeting this immune checkpoint could be crucial for cancer immunotherapy and autoimmune diseases. Clinical data in metastatic melanoma patients pointed out that those patients who showed an increase in CD4^+^ICOS^hi^ T cells after anti-cytotoxic T-lymphocyte-associated protein 4 (CTLA-4) treatment had better overall survival^[4]^. Subsequent preclinical studies in the B16/BL6 melanoma model proved that ipilimumab, an anti-CTLA-4 treatment, induced ICOS^+^FOXP3^-^ T cells, and these, in turn, increased effector T/Treg cell ratio and promoted anti-tumor responses^[5]^. Finally, the administration of antibodies targeting ICOS/ICOS-L interaction together with the inhibition of other negative immune checkpoints, such as programmed cell death protein 1 (PD-1) and CTLA-4, improved anti-tumor immune responses^[6]^.

In recent years, various biological reagents with high binding abilities towards ICOS have been reported and demonstrated excellent performance in preclinical and clinical trials^[7]^. **MEDI-570** is a human antagonistic fucosylated immunoglobulin G1 kappa monoclonal antibody against ICOS with picomole-level binding affinity^[7-8]^. In 2023, Chavez et. al reported the phase I clinical trial results of **MEDI-570** in patients with relapsed/refractory peripheral T-cell lymphoma or angioimmunoblastic T-cell lymphoma (AITL)^[9]^. **MEDI-570** demonstrated encouraging clinical activity in mainly AITL patients with a 44% response rate (including partial response), and the side effects are tolerated^[9]^. Following immunophenotypic analysis revealed a dosage-dependent reduction in CD4^+^ICOS^+^ T lymphocytes in patients after treatment. **JTX-2011** was reported in 2020 by Hanson et. al, which is an agonistic monoclonal antibody towards ICOS for receptor stimulation^[10]^. **JTX-2011** showed promising treatment efficiency on mouse tumor models alone or combined with anti-PD-1 or anti-CTLA-4^[10]^. Currently, **JTX-2011** is still undergoing a Phase I/II clinical trial for evaluating the efficiency alone or in combination with nivolumab, ipilimumab, or perbrolizumab in patients with advanced and/or refractory solid tumors (NCT02904226). Compared with biologicals, small molecules have been demonstrated to have many advantages, such as oral bioavailability, greater tumor penetration, and cell-penetrating capabilities,^[11]^ and therefore receive intense focus in medicine development. Owing to the facility in structure optimization, the long-term immune-related damaging effects and toxicity in the patients could be easily reduced by pharmacokinetics properties optimization. Besides, because of the relatively high stability at room temperature, molecular medicine also demonstrated advantages over biologicals in transport and storage, therefore receiving focus from pharmaceutical industries.

However, as the development work is still in the early stage, rare small molecule inhibitors for ICOS/ICOS-L interaction could be developed,^[12]^ and none is approved for clinical application. Affinity selection mass spectrometry (ASMS) is a mass spectrometry (MS) based high-throughput screening (HTS) technology for hits identification of biological targets (such as protein and DNA), which could identify active compounds from a large number of candidates within a short time.^[13]^ In the ASMS screening, after co-incubating with individual or mixed candidates, the target protein could be separated from free compounds via size-exclusive chromatography (SEC), and liquid chromatography-mass spectrometry will be used to characterize the binding molecules. Compared with other screening methods used for ICOS hit screening, such as pharmacophore-based virtual screening^[12b]^, random screening of chemical libraries^[14]^, and screening of focused libraries of immunomodulators^[15]^, ASMS demonstrated distinct advantages in both screening accuracy and candidate compatibility. Based on the advantage of SEC, the incubation could be performed without any requirement of protein tag or immobilization, which could keep the target protein in its natural condition, thus improving the reliability of identified hits. Besides, as the readout in ASMS is molecular masses of tested molecules, fluorescence compounds could also be selected as candidates, which is challenging in fluorescence assay-based screening, as the signal disturbing might cause false results.

Aiming to identify novel scaffolds as inhibitors of ICOS/ICOS-L interaction, we performed ASMS screening on 15,600 compounds from the Maybridge HitCreator and HitFinder libraries. One compound (compound **9**) with promising ICOS/ICOS-L inhibitory profile was validated and selected as the template for the preliminary structure-activity relationship (SAR) study. After the functional group modification of position R_1_, R_2_, and R_3_ of compound **9**, we validated the key structural features essential for ICOS/ICOS-L inhibition. Furthermore, computational studies were performed to identify a potential binding site of compound **9** in ICOS and to shed light on its key geometrical features.

## Results and Discussion

### ASMS Screening

We first selected a subset of 15,600 compounds comprised within the Maybridge HitCreator and HitFinder libraries. Each one of these commercial libraries is originally composed of roughly 14,000 compounds from the entire 550,000 Maybridge Screening Collection. Their design is based on two main criteria: i) keeping a high and representative diversity of the entire collection, which provides a higher hit probability than larger but less diverse collections; and ii) reliability, only drug-like compounds that meet Lipinski’s Rule of Five^[16]^ and additional filters, such as PSA (polar surface area) ≤ 140Å^2^, are included.

For the ASMS screening, briefly, after co-incubating with a mixture of tested compounds, the complexes of ICOS protein and bound compounds can be purified from the free molecules via SEC. Afterwards, the bound compounds are dissociated from the complex and separated using high-performance liquid chromatography (HPLC) and identified by time-of-flight mass spectrometry (**Figure 1**). As the first step to setting up the amount of protein that was going to be used in the ASMS screening, a desired hit rate of ∼1% was chosen. Different concentrations of ICOS protein concentrations (1 to 0.1 µM), were incubated with a mixture of 1,600 compounds (10 µM) in a physiologically relevant buffer showing a ∼30% to ∼1% hit rate, respectively. Finally, 45 batches for a total of 15,600 compounds were performed and 415 hits were obtained (**Supporting Information, SI, Table S1**).

**Figure 1.**
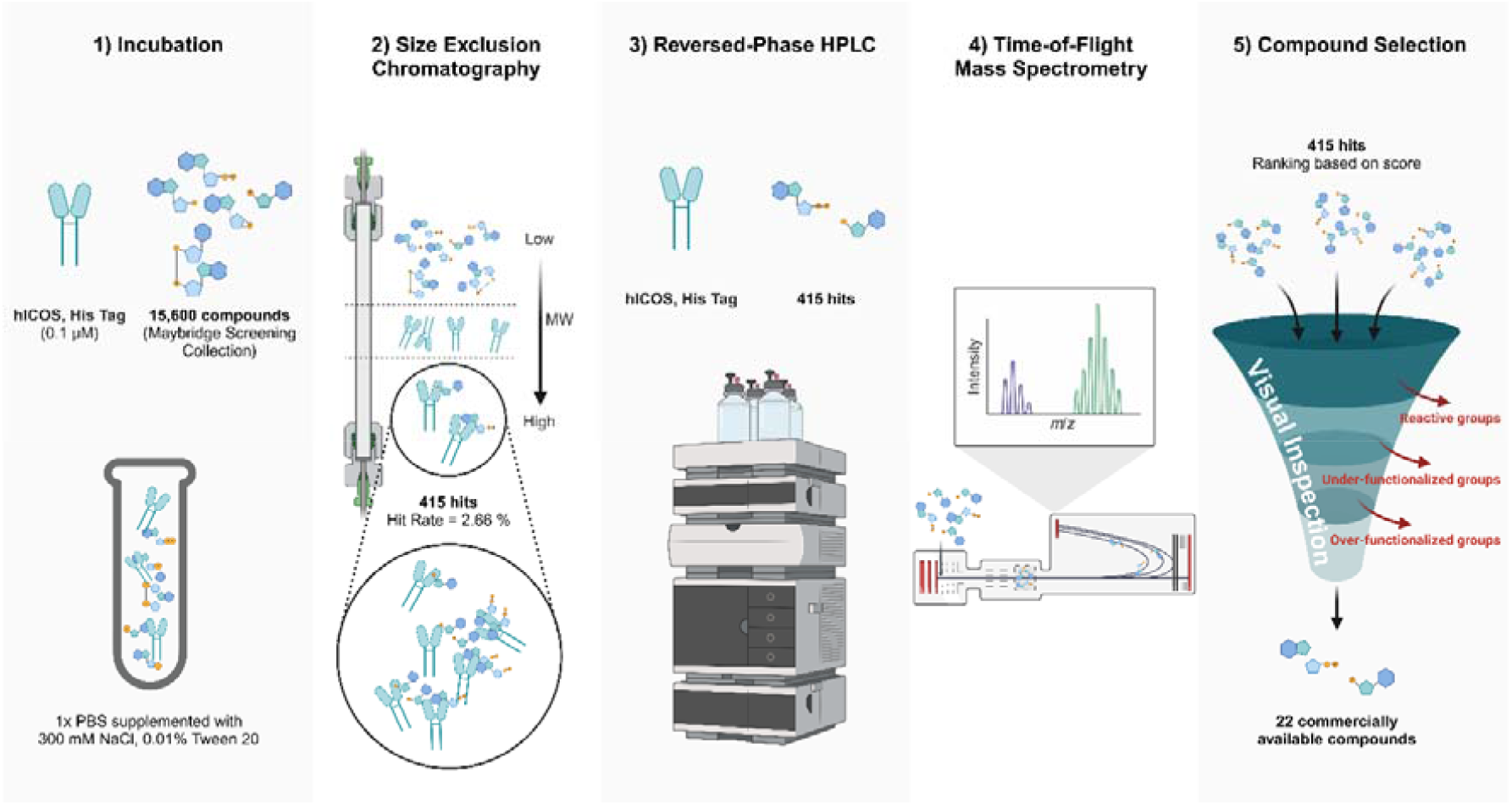
Schematic representation of ASMS screening workflow for small molecule candidates identification.

We performed a visual inspection to select ∼30 compounds (out of the identified potential 415 hits) based on the following criteria: i) removal of compounds that possess reactive groups, ii) removal of compounds that are under-functionalized relative to their size (i.e. do not incorporate sufficient functional groups to engage in specific interaction with ICOS), and iii) removal of compounds that are over-functionalized (i.e. possess numerous functional groups, which leaves no room for optimization). Thus, the aim of this step was to select drug-like compounds that represent good starting points for further optimization. Finally, a total of 22 commercially available compounds were acquired for subsequent hit validation using biorthogonal assays (**Supporting Information, Table S1**). Five out of 22 compounds were excluded due to aggregation or technical issues (scan anomalies, quenching or autofluorescence) observed at the screening platform step: compounds **18** to **22**. Dianthus was selected as the screening platform and surface plasmon resonance (SPR) as the validation platform. The development and optimization of the affinity screening platform, Dianthus, was performed as previously described by our group^[12b]^.

### Biophysical Screening

All 17 compounds were included in a single-dose screening (100 µM or 500 µM) upon incubation with ICOS protein. Compounds that showed a higher than three times the standard deviation of the negative control (empirical rule) were selected for further binding affinity assays (**Figure 2A**). Subsequent binding affinity assays with selected eight small molecules showed that compounds **3, 9, 10** and **11** bound to ICOS protein with variable equilibrium constants, K_D_ values, in the micromolar range: 400.25 ± 177.17 µM, 143.67 ± 79.65 µM, 147.02 ± 140.83 µM, and 162.80 ± 110.85 µM, respectively (**Figure 2B - E**). The remaining candidates did not show a dose response effect (*data not shown*). To confirm these latter results, we measured the binding affinity of the **3**/ICOS, **9**/ICOS, **10**/ICOS, and **11**/ICOS complexes by SPR and confirmed that compounds **3** and **9** bound to target protein ICOS with a K_D_ value equal to 163.67 ± 52.73 µM and 185 ± 126.05 µM, respectively (**Figure 2F - G**). Finally, we tested if compounds **3** and **9** could function as inhibitors of the ICOS/ICOSL interaction. For this, we utilized our previously reported Time-Resolved Förster’s Resonance Energy Transfer (TR-FRET) assay^[12b]^. No dose-dependent response within the range between negative and positive control was observed when ascendent concentrations of compound **3** were incubated with ICOS and ICOS-L proteins in the TR-FRET assay, indicating that no inhibition of ICOS/ICOS-L binding was performed by these compounds (*data not shown*). However, compound **9** showed ICOS/ICOS-L inhibitory capabilities and presented an IC_50_ of 29.38 ± 3.41 µM (**Figure 2H**).

**Figure 2.**
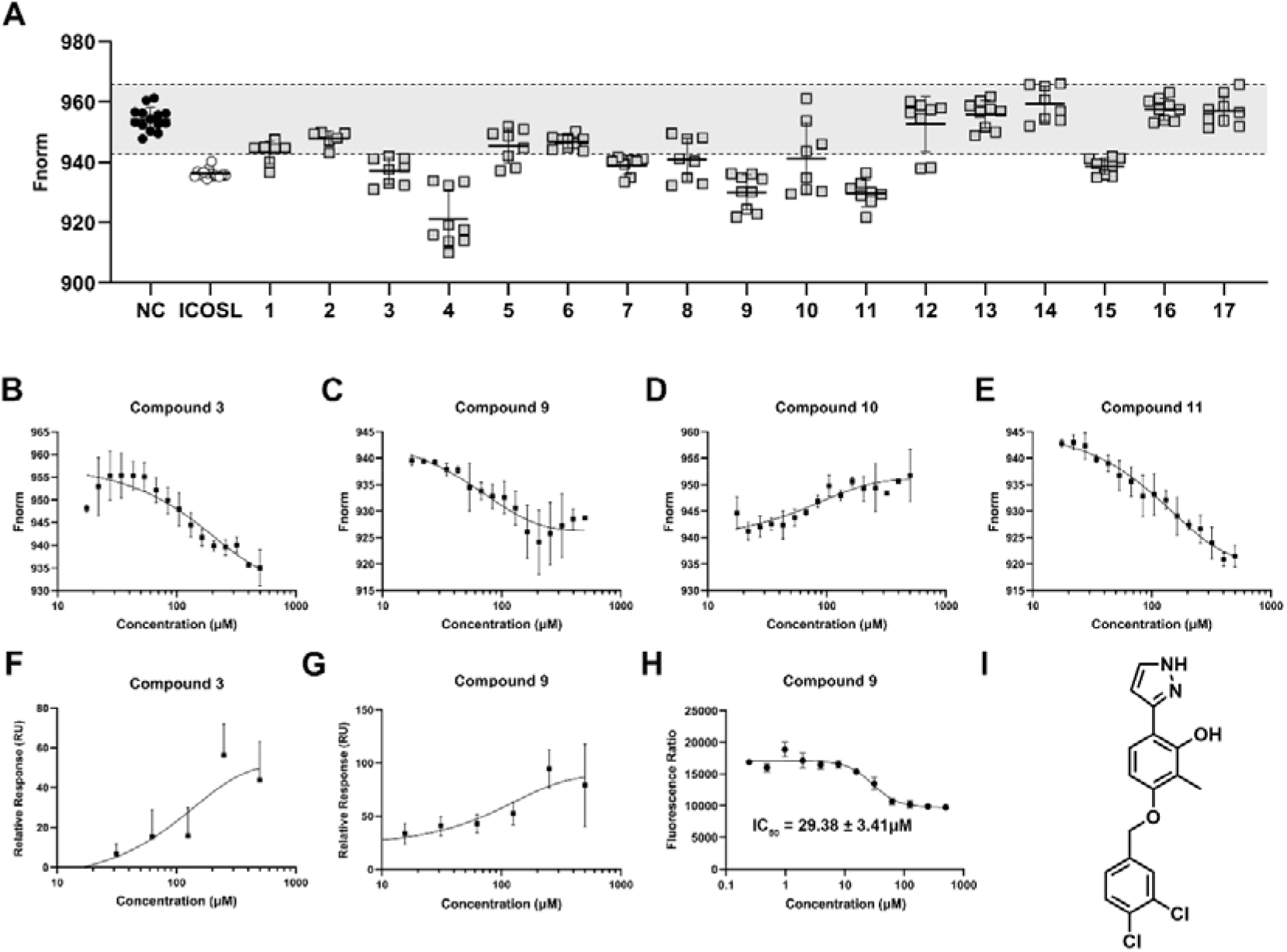
Biophysical screening (using Dianthus) for the 17 candidates. (**A**) Single-dosage screening towards ICOS (10 nM) using Dianthus. Black dots and white dots indicate the negative (ICOS solely) and positive control (ICOS with ICOS-L), respectively. Grey area indicates the F_norm_ difference lower than three times the standard deviation of the negative control. (**B** - **E**) Dosage-dependent curve of compound **3** (**B**), **9** (**C**), **10** (**D**), and **11** (**E**) binding to ICOS using MST. Buffer condition of MST: 10 mM HEPES, 150 mM NaCl, 0.005% Tween 20, 5% DMSO, pH 7.4. (**F** - **G**) SPR binding curve of compound **3** (**F**) and **9** (**G**) towards ICOS. Buffer condition of SPR: 1x PBS-P+ 5% DMSO. (**H**) Inhibition curve of compound **9** with ICOS/ICOS-L (10 nM / 10 nM) using FRET assay. Buffer for RRET assay: PBS with 5% DMSO. (**I**) Structure of compound **9**. F_norm_ and fluorescence ratio were measured in triplicate with results given as the mean ± SD.

### Preliminary structure-activity Relationship (SAR) Study

Inspired by the promising ICOS/ICOS-L inhibitory profile of compound **9**, six close structural analogues with different substituent groups were designed, synthesized, and evaluated for exploring the structure-activity relationship with ICOS protein (**Table 1**). The synthetic routes of compound **D1** - **D6** are shown in **Scheme 1**. For synthesizing derivatives with benzene or pyridine, 2-methylbenzene-1,3-diol was brominated by **OXONE®** at room temperature within the presence of ammonium bromide to provide compound **I1**. 4- (bromomethyl)-1,2-dichlorobenzene was conjugated to **I1** or commercially available 4-bromo-2-methylphenol (**I2**) in alkaline condition to prepare bromine compound **I3** and **I4** with a moderate yield (68.5%). After protecting the hydroxy group with the chloromethyl methyl ether (MOMCl), a Pd(PPh_3_)_4_ catalyzed Suzuki cross-coupling reaction was performed between compound **I3** or **I5** with the phenylboronic acid or pyridin-4-ylboronic acid to provide intermediate **I6** and final product **D2** and **D3** with moderate yields (23.4% - 63.7%). After the deprotection of methoxymethyl in trifluoroacetic/CH_2_Cl_2_ (TFA/DCM, 1/1, v/v) system, the final product **D5** could be prepared (yield: 21.3%). Compared with compounds with six-membered rings, the synthesis of derivatives with the 1H-pyrazol-3-yl group is more complex. Commercially purchased compounds 4-hydroxy-3-methylbenzaldehyde or 2,4-dihydroxy-3-methylbenzaldehyde were conjugated with substituted benzyl bromide to give aldehyde compounds **I7** and **I8** with a high yield (90.2% - 100%). After protecting the hydroxy group with the MOMCl (yield: 78.1%), the Wittig reaction was employed to transform the formyl group into acrylaldehyde (yield: 45.1%). For facilitating the following cyclization reaction, the tosylate group was introduced as the leaving group via the azine condensation between the 4-methylbenzenesulfonohydrazide and corresponding substituent cinnamaldehyde to provide the azine intermediates **I12** and **I13**. The cyclization and deprotection were used in sequence to provide the final product with the 1H-pyrazol-3-yl group (yield: 57.0% - 58.2%). For compound **D6**, after carboxylating the 2-methylbenzene-1,3-diol in CO_2_ atmosphere, 4-(bromomethyl)-1,2-dichlorobenzene was reacted with the coarse product to give the final product with a high yield (two-step yield: 66.3%). The structures of all intermediates were verified by ^1^H-NMR and MS, and the structures of all final products were characterized by ^1^H-NMR, ^13^C-NMR, and MS.

**Table 1.**
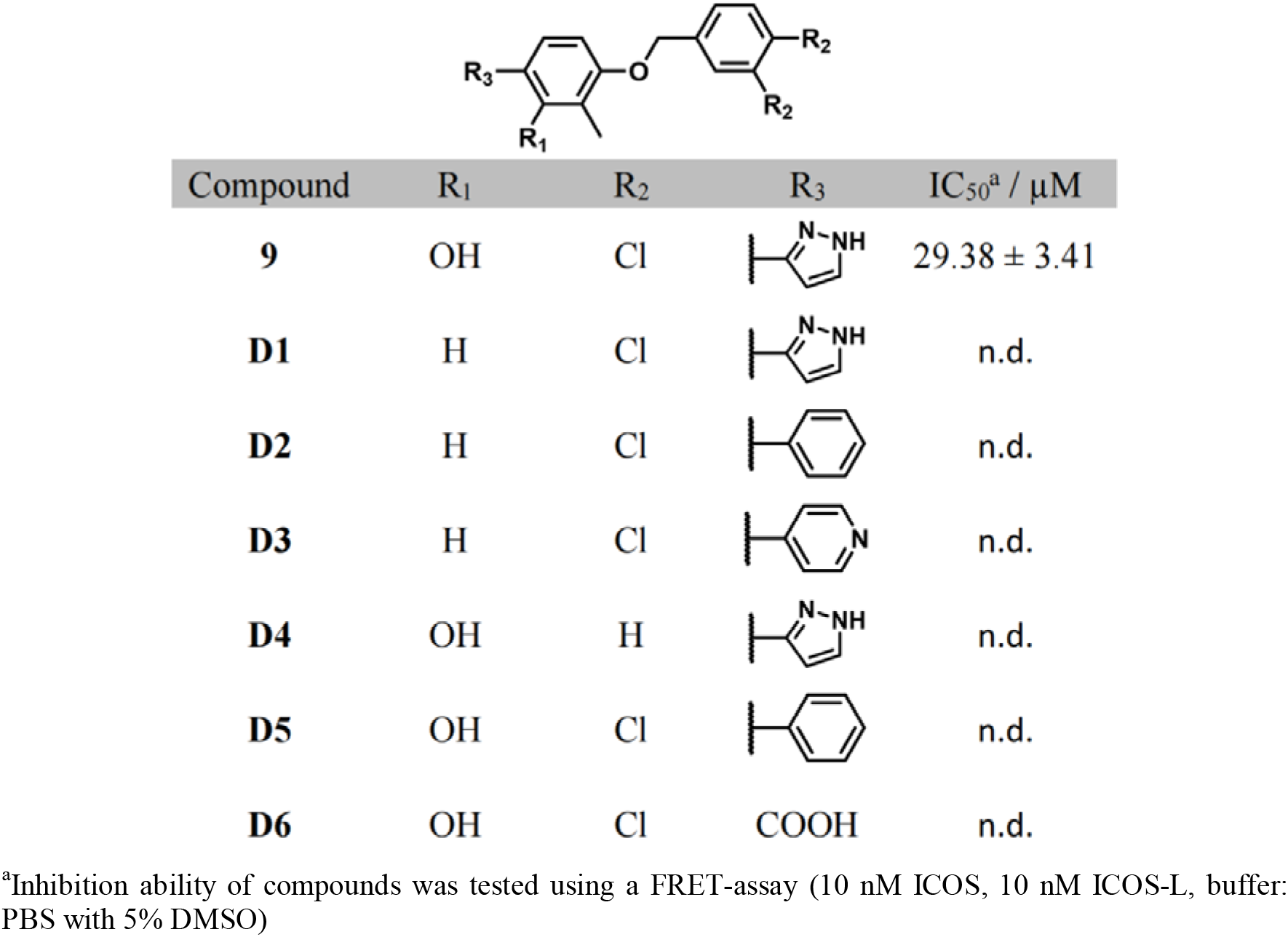
Structures and biological properties compound **9** derivatives.

|  |  |  |  |  |
| --- | --- | --- | --- | --- |
| Compound | R<sub>1</sub> | R<sub>2</sub> | R<sub>3</sub> | IC<sub>50</sub><sup>a</sup> / μM |
| **9** | OH | Cl | | 29.38 ± 3.41 |
| **D1** | H | Cl |  | n.d. |
| **D2** | H | Cl |  | n.d. |
| **D3** | H | Cl |  | n.d. |
| **D4** | OH | H |  | n.d. |
| **D5** | OH | Cl |  | n.d. |
| **D6** | OH | Cl | COOH | n.d. |

<sup>a</sup>Inhibition ability of compounds was tested using a FRET-assay (10 nM ICOS, 10 nM ICOS-L, buffer: PBS with 5% DMSO)

**Scheme 1.**
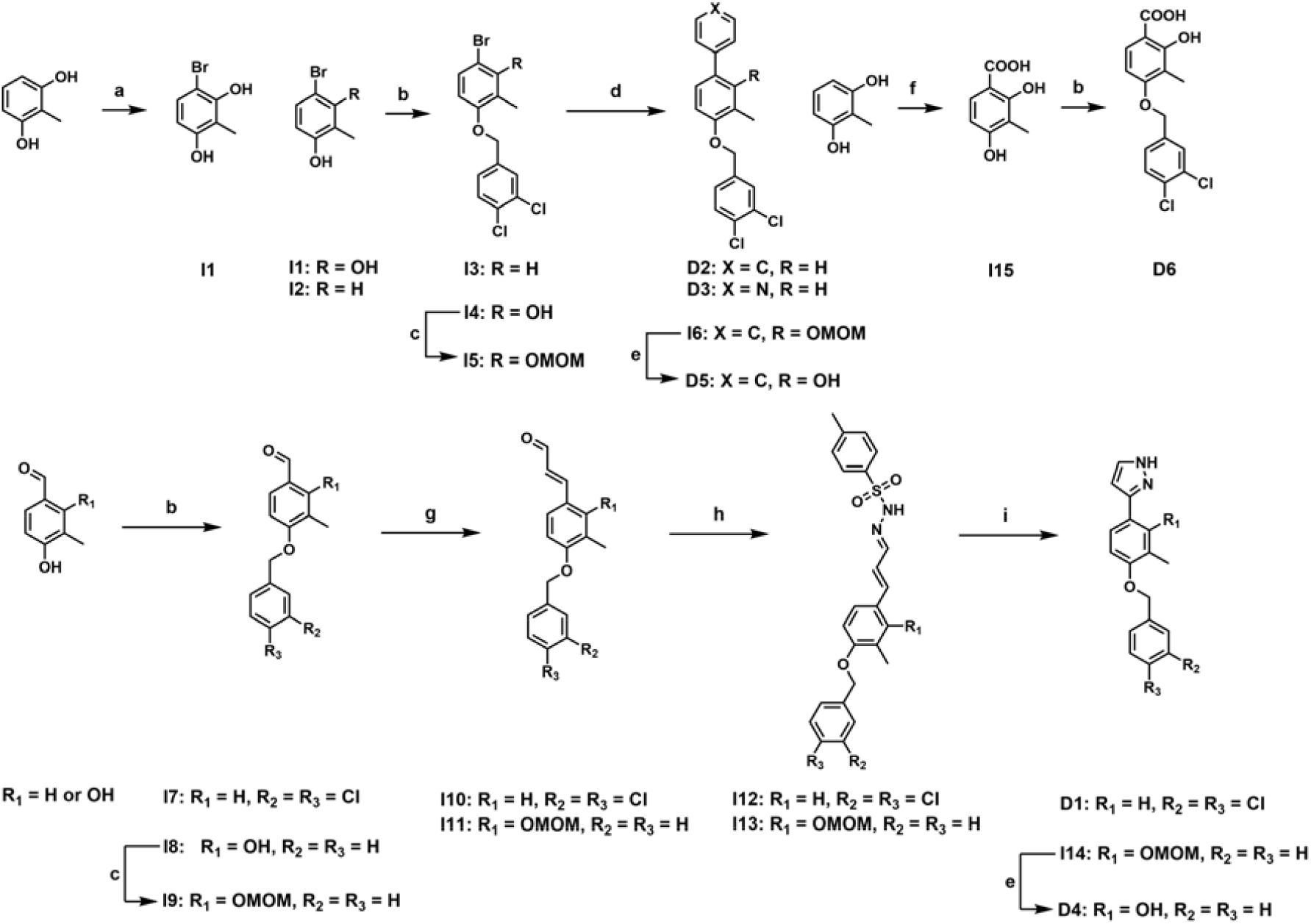
Synthetic route of compound **D1** - **D6**. (a) Oxone^®^, NH_4_Br, MeOH, r.t., 0.5 h; (b) 4- (bromomethyl)-1,2-dichlorobenzene, K_2_CO_3_, KI, DMF, 90 °C, 3 h, 68.5%; (c) MOMCl, DIPEA, DCM, 50 °C, 3 h, three-step yield: 6.1%; (d) phenylboronic acid for **D2** and **I6** or pyridin-4-ylboronic acid for **D3**, Ph(PPh_3_)_4_, K_2_CO_3_, toluene, EtOH, H_2_O, 90 °C, overnight, 24.3% - 63.7%; (e) TFA, DCM, r.t., 1 h; (f) CO_2_, DBU, ACN, r.t., 2 d; (g) (1) (1,3-Dioxolan-2-ylmethyl)triphenylphosphonium bromide, NaH, 18-crown-6, THF, r.t., overnight; (2) HCl (1 M aqueous solution), THF, r.t., 0.5 h, 45.1%; (h) 4-methylbenzenesulfonohydrazide, MeOH, 60 °C, 3 h, 58.2% - 71.3%; (i) CsF, TEBAC, THF, 70 °C, overnight, 57.0% - 58.2%.

For studying the impact of the functional group substitution on the ICOS/ICOS-L inhibitory profiles, all compounds were tested using the ICOS/ICOS-L TR-FRET assay. As shown in **Table 1**, for compounds without the 2-OH group at R_1_ (**D1** - **D3**), dramatic decreases in the inhibition abilities of ICOS/ICOS-L interaction could be observed. In particular, for **D1**, which shares the same structure as compound **9** except for the depletion of the ortho-hydroxy group, no dosage-dependent change could be observed with the increasing concentration of compound (**Figure S1**, starting from 500 µM, 1.3-fold dilution), indicating the critical role of 2-OH in the binding-site recognition. Besides, the dichloro group on the benzyl ring also plays an important role in the ICOS binding of compound **9**, as **D4** demonstrated an inhibition activity elimination after the hydrogenation of ortho-dichlorobenzene (**Table 1**). As an aromatic ring with two adjacent nitrogen atoms, theoretically, 1*H*-pyrazole could stabilize the molecule/protein interaction via π - π stacking or forming hydrogen bonds with amino acid residual from the surrounding environment^[17]^. For studying the exact effect of the 1*H*-pyrazole in the binding with ICOS, new derivatives with sole π - π stacking (**Table 1, D5**, benzene) or hydrogen bond forming (**Table 1, D6**, carboxylic acid) ability were designed and tested. Interestingly, even though the dramatic deterioration in the inhibition of ICOS/ICOS-L interaction was detected for both derivatives, **D6** showed relatively better inhibition ability than **D5**, as the dosage-dependent effect could be detected starting from 295.9 µM of **D6**, while only random spots were found in the other compound, indicating that 1*H*-pyrazole stabilizes the ICOS binding via hydrogen binding instead of π - π stacking.

### Docking studies and molecular dynamics (MD) simulations

Comparative analysis of interactions between **9** (active compound) and **D1** (inactive compound) bound in ICOS was done based on the results obtained from two docking programs, i.e. GOLD (v.2021.02)^[18]^ and Autodock Vina (v.1.2.5),^[19]^ and those generated afterwards during 100 ns of unrestricted NPT molecular dynamics (MD) simulations, done with NAMD (ver. 2.14)^[20]^ software and AMBER^[21]^ force field. All MD simulations, as well as system setup, were performed as described in the Experimental Procedures (**SI**). The MD simulations were based on consensus docking poses for **9, which** were obtained by applying the procedure described in detail in **SI**. Six potential binding sites were identified, α, β, γ, δ, ε and φ, as shown in **Table S2** (**SI**) and highlighted in **Figure S2**. Results of MD simulations allowed us to conclude that only one variant, with **9** bound in the φ site of ICOS could be considered as the most promising binding place. This choice was justified by a small root mean square deviation (r.m.s.d.) of 4.4 Å at the end of the simulations, computed for the heavy atoms of the central ring of compound **9**. This value indicates a modest shift of compound **9** from its original docking position during MD simulations. Additionally, a long-lasting hydrogen bond contact (present in 69.6% of the snapshot population) was established between Gln54 of ICOS and the ortho-hydroxyl of compound **9**. The other identified binding sites were excluded based on significantly larger r.m.s.d. values or lack of H-bond contact, as described in the **SI** and shown in **Table S2 (SI)**.

To further confirm that the φ site is the most probable **9** binding region, two more replicas of 100 ns unrestricted NPT MD simulations were performed. Both confirmed the originally explored behaviour of **9**, as shown in **Table S3** and illustrated in **Figure 3A**, where the time evolution of orientation of **9** in the binding site and the r.m.s.d values are illustrated. Subsequently, three additional MD simulations for **D1** were done. The initial structure of the ICOS-**D1** complex was prepared based on a simulated ICOS-**9** variant, where the ortho-hydroxyl group of **9** was substituted by a hydrogen atom, transforming it directly to **D1**. Significantly different behaviour of **D1** during MD simulations in the binding site was observed as shown in **Figure 3B**. As can be seen, during simulations the **D1** compound was drifting away from the binding site in all cases, as reflected by a significant increase in the r.m.s.d. value in comparison to the **9** compound. This result agrees with experimental findings and confirms that the hydroxyl group is key for the recognition step. Therefore, φ pocket can be considered the most probable place to bind **9**.

**Figure 3.**
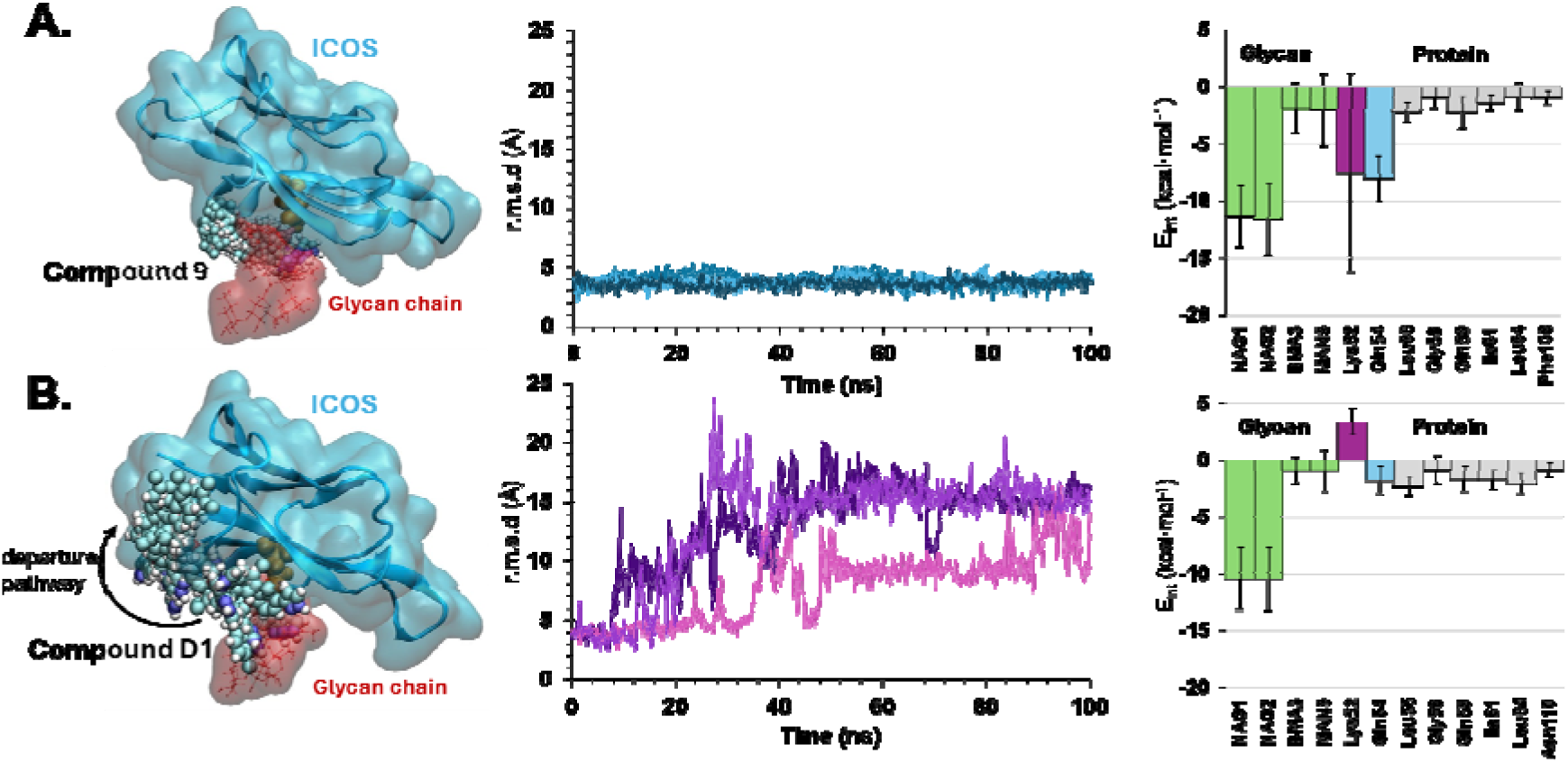
Results of the 100 ns of unrestricted MD simulations for ICOS-compound **9 (A)** and ICOS-**D1(B)** complexes. The time evolution of r.m.s.d computed for the heavy atoms of the central ring of compound **9** and **D1** and values of non-bonding interaction energies (E_int_) were estimated from 30,000 structures generated in three replicas for **9** while in the **D1** case, only those frames where r.m.s.d. was found below 5.5 Å were considered.

Geometrical insights into the ICOS-compound **9** complex allowed the conclusion that **9** does not bind to any particular cavity, instead, it is placed between the surface of the protein and the Asn110-NAG-NAG-BMA-MAN2 glycan chain, as shown in **Figure 4A**. The binding site can be divided into two regions: one part with strong hydrophobic and one with strong hydrophilic character. The o-cresol and pyrazole rings of **9** are located within the latter, where the key H-bond is formed. The remaining dichlorinated ring is placed in the hydrophobic area. To obtain more insight into why **9**, in contrast to **D1**, inhibits ICOS protein, the average values of non-bonding interaction energy (E_int_) were computed per residue for the protein, glycans and both studied compounds, as shown in **Figure 3 and Tables S4** and **S5** (**SI**).

**Figure 4.**
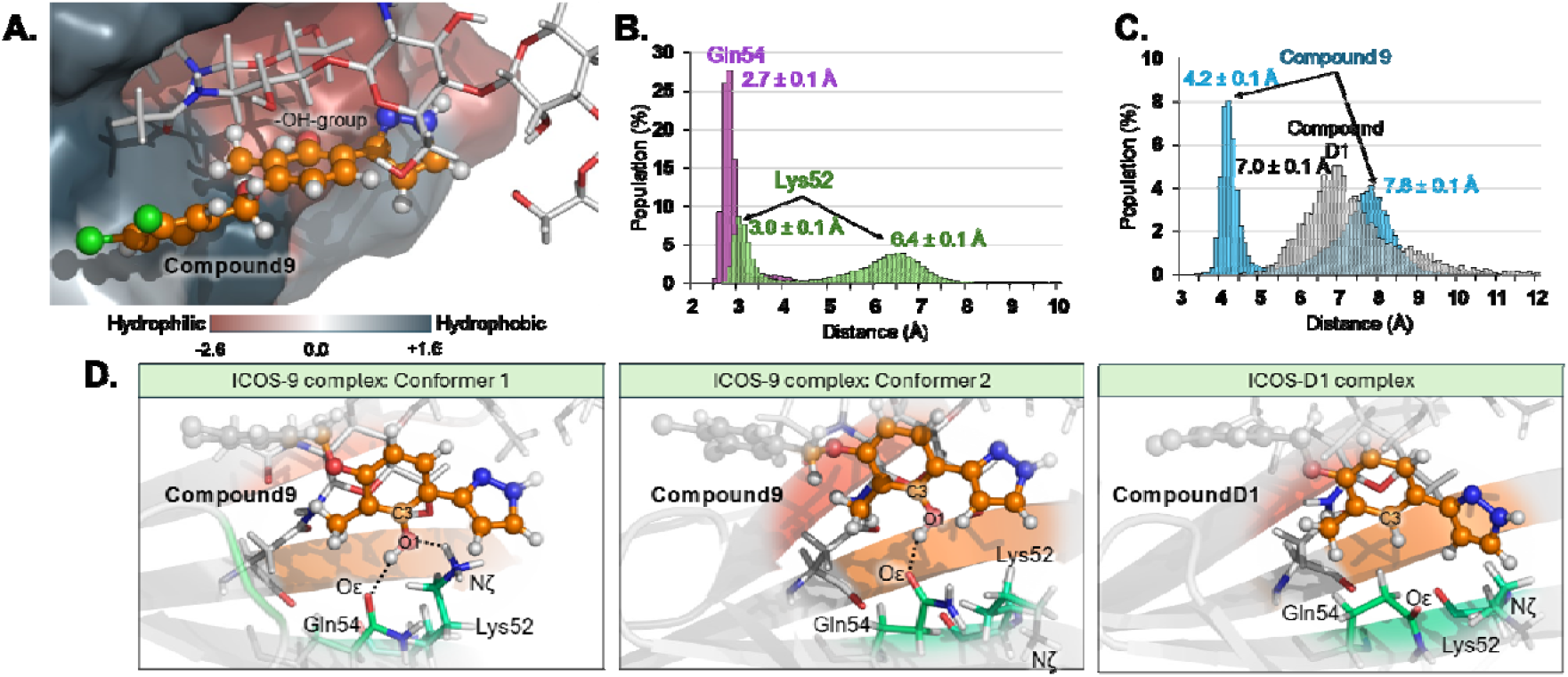
**(A)** Hydrophobicity map generated on the φ binding site of ICOS using the Eisenberg hydrophobicity scale^[22]^. **(B)** Population analysis of the distances between the O1 atom of **9** and Gln54:Oε (magenta) and Lys52:Nζ (green) and **(C)** between Lys52:Nζ and C3 of 9 (Blue) and **D1** compound (grey). All 30,000 structures generated in three replicas were included in the analysis for **9**. In the case of the **D1** only frames with r.m.s.d. below 5.5 Å were considered. **(D)** The structure of the φ binding site of ICOS with the relative position of the Lys52 and Gln54 regarding positions of **9** and **D1**.

The main difference observed for **9** and **D1** is the strength of interaction energy established with Gln54 and the inversion of interaction character with Lys52 residue. While the difference in values of E_int_ with Gln54 (−8.1 ± 2.0 and −1.8 ± 1.2 kcal/mol for **9** and **D1**, respectively) was expected and is rather obvious due to the existence of a strong H-bond formed between **9** and this residue (the contact that is not present in the case of **D1**), the role of Lys52 is not so evident. Nonetheless, its presence seems to be important because depending on the used compound it forms periodically attractive (in the case of **9**, −7.5 ± 8.7 kcal/mol) or repulsive (in the case of **D1**, 3.4 ± 1.1 kcal/mol) interactions. Furthermore, it is highly likely, that repulsive forces between D1 and Lys52 could initially cause its departure from the binding site. Therefore, additional analysis of the behaviour of Lys52 during MD simulations was done. As shown in **Figures 4B and 4C** and **S4** (**SI**), when **9** is bound to ICOS, Lys52 oscillates between two accessible conformations. In the first (most populated), a close distance to the ortho-hydroxyl group of **9** is established providing additional short-lived hydrogen bond contact (around 20 ns), while in the second configuration, a significantly distal position of Lys is observed. Nevertheless, as shown in **Figure S4** (**SI**) it seems that when ICOS is complexed with **9**, Lys52 can freely switch between one and another conformation, which only periodically provides a complementary stabilization effect. Two alternative conformers of lysine are shown in **Figure 4D**. In the case of **D1**, in contrast, only one conformation of Lys52 is available, as shown in **Figure 4D**. As demonstrated by E_int_ calculations this orientation of Lys52 results in the destabilisation of **D1**-Lys52 mutual interactions.

## Conclusion

An ASMS screening was performed on 15,600 compounds from the Maybridge HitCreator and HitFinder libraries for scouting the novel molecular inhibitor hits of ICOS/ICOS-L interaction. Followed by the biological properties verification, compound **9** with an IC_50_ value of 29.38 ± 3.41 µM for ICOS/ICOS-L inhibition was identified. In the following SAR study, it was noticed that the ortho-hydroxy group of **9** might play a critical role in the protein binding process, and the hydrogenation could lead to a dramatic decrease in the inhibition activity. Similar outcomes were obtained in the dichloro group (R2) or 1*H*-pyrazole ring (R3) substitution. Following MD simulation study revealed that the hydroxy group could stabilize the ICOS binding via the hydrogen bond formation with the Gln54:Oε or Lys52:Nζ on the glycan, which could be the reason for the activity deterioration in **D1**, as the -OH group was hydrogenated on the derivative. Our work validates a new scaffold as an inhibitor for the ICOS/ICOS-L interaction and paves the way for the development of more potent ICOS/ICOS-L modulators. Importantly, our work introduces ASMS screening as a promising tool to identify small molecule inhibitors of immune checkpoints.

## Conflict of Interest

The authors declare no conflict of interest.

## Data Availability Statement

The data that support the findings of this study are available in the Supporting Information.

## Acknowledgement

We gratefully acknowledge financial support from the the NIH (Award ID: R01DK137299). We would like to thank Dr. Somaya Abdel-Rahman for assistance with the ICOS/ICOS-L TR-FRET assay. This work was supported by the Spanish Ministerio de Ciencia, Innovación y Universidades (ref. PID2021-123332OB-C21), the Generalitat Valenciana (PROMETEO with ref. CIPROM/2021/079). K.Ś. thanks to Ministerio de Ciencia e Innovación and Fondo Social Europeo for a Ramon y Cajal contract (Ref. RYC2020-030596-I). The authors wish to thank the staff of the Servei d’Informàtica of the Universitat Jaume I, as well as the local computational resources founded by Generalitat Valenciana - European Regional Development Fund (REF: IDIFEDER/2021/02).

## Table of Contents

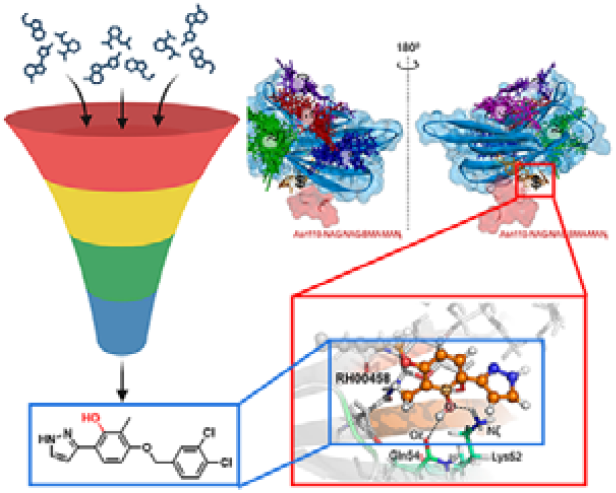

## Notes

### Competing Interest Statement

The authors have declared no competing interest.

