## Supporting Information for "Discovery of ICOS-targeted small molecules using affinity selection mass spectrometry screening"

### Experimental Procedures

#### Affinity selection mass spectrometry (ASMS) screening

Assay setup and conditions: ASMS was done using the Automated Ligand Identification System (ALIS) on an Agilent 2D-HPLC system interfaced to an Agilent Time-of-Flight MS. The assay conditions were 0.1 μM ICOS using PBS (300 mM NaCl, 0.01% Tween20). Source of ICOS for ASMS: human ICOS Protein, His Tag (Cat # ICS-H52H6, Acro Biosystems, Newark, DE, USA)

ASMS Method: The ALIS instrumentation consists of an Agilent 1260 HPLC pump for the size-exclusion chromatography coupled to an Agilent 1290 UHPLC pump for the reversed phase (RP) chromatography with a high-pressure switching valve interfaced to an Agilent 6230B Time-of-Flight Mass Spectrometer. All MS acquisitions were done in positive ion acquisition mode. Data analysis was performed with a combination of Agilent Mass Hunter and proprietary custom software.

- SEC conditions: Buffer A: 700 mM Ammonium Acetate Buffer B: 70% Acetonitrile Column: 50 x 2.1 mm Polyhydroxyethyl A, 3 μm 200 Å porosity (PolyLC)
- RP conditions: Buffer A: Water + 0.1% Formic Acid Buffer B: 90% Acetonitrile + 0.1% Formic Acid Column: 50 x 2.1 mm Kinetex 2.6 um C18 100 Å (Phenomenex)

45 pools of compounds for a total of 15,600 compounds (from the Maybridge HitCreator and HitFinder libraries) at 10 μM were prepared in DMSO. Multiple copies of single use assay plates were prepared by dispensing 250 nL of each pool into a 384-well plate. The test compounds in DMSO were diluted to 10 mL in the assay buffer (PBS, 300 mM NaCl, 0.01% Tween20). One copy of the 45 pools was run in a conventional HPLC-MS experiment to verify that the compounds could be detected. Data was analyzed using proprietary ASMS software. A total of 415 hits were identified.

#### General Information

Compounds **1** - **22** and **32** were obtained from commercial vendors in lyophilized format (**Table S1**). These molecules were dissolved in DMSO (D8418, Sigma Aldrich, St. Louis, MO, USA) at 10 mM and stored at -30°C until use. All reagents for synthesis were purchased commercially and used without purification. Reaction were monitored by thin-layer chromatography (TLC) on TLC plates (AAdvance Instruments, Gel 60 F254 type, 60 Å, USA) and liquid chromatograph mass spectrometer (LC-MS, Acquity UPLC H-Class system equipped with a PDA eλ detector and SQ detector 2, Waters, USA). HPLC analysis was performed on an Kinetex 1.7u C18 reverse phase column (Phenomenex, 100A, 50 mm × 2.1 mm, USA) and eluted at a flow rate of 1.0 mL/min. Mobile phase A was water (with 0.1% formic acid), and mobile phase B was acetonitrile (with 0.1% formic acid). After reaction, the corase production were loaded on RediSep^®^ Silver column (12 g, 24 g, Teledyne ISCO, Lincoln, NE, USA) and purified by a Combi-Flash Rf^+^ Lumen system (Teledyne ISCO, Lincoln, NE, USA). All intermediates were verified by ^1^H NMR and MS spectra, and ^1^H and ^13^C NMR and MS spectra were used to characterize all final products. ^1^H and ^13^C NMR spectra were collected on a Bruker Avance III 500HD (500 or 125 MHz, Billerica, MA, USA) in CDCl_3_ (DLM-7-0312, Cambridge Iscotope Laboratories, Inc., Tewksbury, MA, USA) or DMSO-*d*_6_ (296147, Sigma Aldrich, St. Louis, MO, USA) at room temperature. Chemical shifts were reported in parts per million (ppm) compared with tetramethylsilane (δ 0.00, s). Coupling constants were reported in hertz (Hz). Multiplicities were reported as s (singlet), d (doublet), t (triplet), and m (multiplet). MS spectra were collected on the SQ detector 2 mass spectrometer (Waters, USA). MST-based binding affinity measurement were performed on a NT.23 Pico Dianthus (NanoTemper Technologies, Munich, Germany). SPR-based binding test were performed on a Biacore^TM^ 8K (Cytiva, Marlborough, MA, USA). TR-FRET measurements were acquired on a Tecan Infinite M1000 Pro equipment (Tecan, Männedorf Switzerland).

#### Dianthus: Affinity Screening Platform

The Monolith His-Tag Labeling Kit RED-tris-NTA 2^nd^ Generation (Cat # MO-L018, NanoTemper Technologies, München, Germany) was used for the labeling of the human ICOS Protein, His Tag (Cat # ICS-H52H6, Acro Biosystems, Newark, DE, USA) following manufacturers’ instructions. Briefly, after determining the affinity of the His-labeling dye for the His-tagged human ICOS protein, 100 nM dye solution was mixed with 200 nM human ICOS protein in assay buffer and incubated for 30 minutes at room temperature in the dark before every experiment.

Compounds were diluted in assay buffer to the corresponding final concentration in the presence of 5% DMSO. Human ICOS-L, Fc Tag (Cat # B72-H5254, Acro Biosystems) in assay buffer supplemented with 5% DMSO was used as positive controls while 5% DMSO in assay buffer was used as a negative control.

Labeled protein was further diluted to 20 nM in assay buffer, incubated 1:1 with the corresponding compound or control for 2 hours at room temperature in the dark, and centrifuged for 30 seconds at 1000 *xg* before loading into Dianthus NT.23 Pico (NanoTemper Technologies). The assay buffer was 10 mM HEPES, 150 mM NaCl, 0.005% Tween 20, pH 7.4 and the reaction volume was 20 µl for all experiments. The equipment was set to 25°C as set temperature and samples were measured for 1 second without heating (before infrared laser activation) and during 5 seconds with the infrared laser turned on. Results are displayed as fluorescence values: i) relative fluorescence, fluorescence obtained during the length of the experiment (TRIC) and normalized to the fluorescence obtained at the set temperature; ii) normalized fluorescence (Fnorm, %): ratio between fluorescence values after and before laser activation.

In the single-dose experiments, three technical replicates were performed per compound, and two technical replicates were performed for the positive and negative controls at regular intervals in the 384-well plate. In the binding affinity experiments, a 16-point serial dilution was prepared for every selected compound. Three independent experiments were performed in the single-dose and binding affinity experiments to determine potential binding molecules and affinity constants, respectively. Data was acquired and analyzed using DI.Control and DI.Screening Analysis software (NanoTemper Technologies), respectively.

#### Surface Plasmon Resonance

The binding analysis was performed on a Biacore^TM^ 8K instrument (Cytiva, Marlborough, MA, USA) at 25°C using 1x PBS-P+ buffer (Cat # 28995084, Cytiva, Marlborough, MA, USA) supplemented with 5% DMSO as the running buffer. ICOS was captured on a sensor chip NTA at a flow rate of 10 μl/min for 60 seconds yielding an immobilization level of approximately 2,250 RU using the NTA reagent kit (Cytiva, Marlborough, MA, USA). The solutions provided are a nickel solution (0.5 mM NiCl_2_) for creating the chelating surface on the sensor chip NTA and an EDTA (350mM) solution for regeneration of the surface. For binding analysis, serial dilutions of the analyte (500 µM to 7.81 µM, 2-fold dilution, 7 channels) were prepared in running buffer and injected over the chip at a flow rate of 30 μl/min for 120 seconds (association phase). The dissociation phase was set at 30 minutes. Data was obtained using the Biacore 8K Control Software (Cytiva, Marlborough, MA, USA) and analyzed by non-linear curve fitting using a steady-state affinity analysis applying the software Biacore^TM^ Insight Evaluation Software (Cytiva, Marlborough, MA, USA).

**Time-Resolved Förster’s Resonance Energy Transfer (TR-FRET) Assay**

In a 96-well low volume white assay plate, gradient concentration (2-fole dilution from 500 µM) of tested molecules were mixed with Fc-tagged human ICOS (Cat # 10344-H02H, Sino Biological, Beijing, China), His-tagged human ICOS-L (Cat #11559-H08H, Sino Biological, Beijing, China), XL665 labeled anti-human Ab (Cat # 61HFCXLF, Revvity, Waltham, MA, USA), and Tb cryptate labeled anti-His mAb (Cat # 61HI2TLF, Revvity, Waltham, MA, USA). The final concentration of two proteins are 10 nM. After incubating at room temperature for 1 hour with gentle shaking, the measurement were performed on the Tecan Infinite M1000 Pro equipment (λ_ex_: 340, λ_em_: 620 nm). The signals were calculated as a ratio: Ratio = Fluorescence_665nm_ / Fluorescence_620nm_ × 10^4^, and were measured in triplicate with results given as the mean ± SD. Buffer condition: PBS with 5% DMSO.

#### System Setup and Blind Docking

#### The ICOS protein model was built based on atom cartesian coordinates provided in the Protein Data Bank (https://www.rcsb.org/) for the ICOS-ligand complex as available under PDB ID: 6x4g^[1]^. For this study, the ICOS-L was removed from the structure, and missing residues in the chain of the ICOS protein, i.e., amino acid residues located between positions 22-30 and 129-130 were added as described in detail previously^[2]^. The generated improved ICOS structure was directly used in blind docking studies of compound 9 which were performed using six different scoring functions (SFs) from two software packages: GOLD (v.2021.02)^[3]^ and Autodock Vina (v.1.2.5)^[4]^. The used SFs were ASP^[5]^, CHEMPLP^[6]^, ChemScore^[7]^ and GoldScore^[8]^ from GOLD, and Vina^[4a]^ and Vinardo^[9]^ from Autodock Vina. The protein was prepared for docking using the built-in tools provided by each program: in GOLD, 937 hydrogen atoms were added and the ligand and protein atom types were assigned automatically. The formal charge of atoms was in this case deduced by counting the bond order of the atom and comparing the result with the atom's normal valency. On the contrary, in the Autodock Vina approach, partial atomic charges of amino acids were assigned explicitly to each atom based on the Kollman charges^[10]^, and those for the ligand and atypical protein structures such as glycans were based on the Gasteiger model.^[11]^ The coordinates for the blind docking were set to x = 37.528, y = -53.372, z = -6.577, with a radius of 30 Å in all three x, y and z axes, which was enough to encompass the whole protein. A maximum of 100 binding poses of the chosen compound were calculated by each SF. During docking the protein atoms were fixed, while the ligand was treated as fully flexible. The structural outputs were clustered using the complete linkage algorithm with a cut-off of 4 Å. In the case of the GOLD software, resulting positions were clustered with the built-in tools and in Autodock Vina and Vinardo with the ClusDOCK tool from the PacDOCK web server^[12]^. The overall central position of these clusters indicated the existence of six potential binding regions of Compound 9 (see Figure S2). Based on obtained positions the coordinates for each region were defined by visual inspection taking into account the approximation of the center of mass of each cluster and were used for targeted docking purposes. The coordinates for each of the putative sites are given in Table S6.

**Site-Specific Docking**

In order to identify probable binding conformations, targeted docking simulations were performed for the six identified sites using the same SFs, the same parameters and treatment for protein and ligand as described above. In these docking studies, a 10 Å search radius was used. In GOLD, the diverse solution option was turned on to increase the coverage of the conformational space. In the Autodock Vina SFs, this was unnecessary, since they enforce diversity by design^[4a]^. The complete set of docking results for each site and SF was clustered individually using the complete linkage algorithm, with a 2 Å cut-off, to obtain distinct sets of poses for each site. Then, the representative structures of the clusters obtained for the six SFs and each site were reclustered together using the cluster average algorithm, with a cut-off of 2 Å. This allowed obtaining consensus poses which increased the probability of obtaining more realistic binding conformation^[13]^. For GOLD, only clusters indicated by three or more SFs were selected while for Vina, only poses indicated by both SFs and having negative binding energy were accepted. Moreover, the cases where the SFs of both programs agreed were considered. This resulted in 5 poses for α, 12 for β, 8 for γ, 25 for δ, 24 for ε and 21 for φ sites. A summary of the consensus docking approach^[13]^ is given in **Figure S3**.

**Molecular Dynamics Simulations**

To reduce the amount of consensus docking poses, only those variants that potentially formed a hydrogen bond between the -OH group of **9** and polar groups of the protein were selected for further probing through Molecular Dynamics (MD) simulations. This choice was supported by a hypothesis that the presence of the -OH group is vital to **9** activity. This assumption was made based on experimental findings, which proved that compound **D1**, depleted of this group, was inactive as shown in **Table 1**. Based on this metric, eight poses were selected from three different sites, i.e. one in β (β1), five in δ (δ1, δ2, δ3, δ4 and δ5) and two in φ (φ1, and φ2). Crystal structures of ICOS with the docked poses were used as the initial model for MD simulations at the molecular mechanics (MM) level. AMBER^[14]^ force field (FF) parameters for standard amino acid residues were obtained from ff19.SB^[15]^ FF, while water and counterions parameters were adapted from TIP3^[16]^ FF. Finally, parameters for non-standard Asn residues with N-linked glycan and N-linked two N-acetyl glucosamine (GlcNAc) and three mannose residues of the N-linked glycan at position 110 were obtained from GLYCAM_06j-1^[17]^ FF. Missing FF parameters for compounds **9** and **D1** were generated with GAFF2^[18]^ using the Antechamber^[19]^ software and the atomic charges were computed with the AM1 method with bond charge corrections (AM1-BCC),^[20]^ according to the standard procedure as used in our previous studies^[21]^. All newly assigned parameters are provided in **Tables S3** and **S4**. The protonation state of titratable residues was assigned according to pK_a_ results computed in our previous work on the ICOS-ICOSL system^[2]^. Missing hydrogen atoms were added using the LEAP^[22]^ module of the AmberTools package, which was also used to add an orthorhombic box having approximate dimensions of 68 x 61 x 76 Å^3^. Additionally, 3 negatively charged chloride (Cl^-^) counterions were added to the system at electrostatically favourable positions to neutralize its total charge. The MD simulations were done using NAMD (ver. 2.14)^[23]^, with a cut-off for non-bonding interactions set between 14.5 and 16 Å applied with a smooth switching function. Along the simulations, the temperature was controlled using the Langevin thermostat^[24]^ and the pressure with the Nosé-Hoover Langevin piston^[25]^. The systems underwent energetic optimizations and then were gradually heated to 300 K with a 0.1 K temperature increment, followed by 500 ps of non-biased NPT equilibration. The structures after equilibration were used as starting geometries for 100 ns unrestricted NPT MD simulations with a 2 fs integration step using the SHAKE algorithm^[26]^. In all simulations, periodic boundary conditions (PBC) were applied. The cpptraj^[27]^ program was then used to analyze the results. The time evolution of the r.m.s.d of the ligand’s central ring, i.e. atoms C1, C2, C3, O1, C5, C11, C12 and C13, as labelled in the structure scheme provided in **Table S7** and **S8**, was calculated to measure the deviation of each pose from the initial docking guess. The non-bonding interaction energy (E_int_) as a sum of electrostatic (E_elec_) and Lennard-Jones (E_LJ_) interactions was computed using Linear Interaction Energy (LIE) with a default cut-off set to 12 Å. Stability of hydrogen bond contacts was monitored, using as reference 3.3 Å distance between donor and acceptor and 135º angle define between donor, hydrogen and acceptor atoms.

#### Chemistry

**4-bromo-2-methylbenzene-1,3-diol** (**I1**). To the solution of 2-methylbenzene-1,3-diol (620.7 mg, 5 mmol, 1 eq) in MeOH (50 mL), Oxone^®^ (1690.5 mg, 5.5 mmol, 1.1 eq) and ammonium bromide (516.5 mg, 5.5 mmol, 1.1 eq) were added. The mixture was stirred at r.t. for 0.5 h, and used in the next step without further purification.

**4-bromo-1-((3,4-dichlorobenzyl)oxy)-2-methylbenzene** (**I3**). To the solution of 4-bromo-2-methylphenol (374.1 mg, 2 mmol, 1 eq) in DMF (60 mL) was added 4-(bromomethyl)-1,2-dichlorobenzene (575.8 mg, 2.4 mmol, 1.2 eq), K_2_CO_3_ (1105.6 mg, 8 mmol, 4 eq), and KI (33.2 mg, 0.2 mmol, 0.1 eq). The mixture was stirred at 90 ^o^C for 3 h and monitored by TLC. After being cooled to room temperature, the solvent was removed by evaporation in vacuum. 40 mL H_2_O was added to the residue and extracted by CH_2_Cl_2_ (3 x 40 mL). The combined organic phase was dried via MgSO_4_ and concentrated under vacuum. The crude product was purified by the Combi-Flash system (A = hexane, B = DCM; B%: 0% to 25%, 32 min) to give **I3** as a white solid (475.0 mg, 68.5%). ^1^H NMR (500 MHz, CDCl_3_) δ 7.54 (s, 1H), 7.48 (d, *J* = 8.3 Hz, 1H), 7.31 (s, 1H), 7.28 – 7.24 (m, 2H), 6.71 (d, *J* = 8.7 Hz, 1H), 5.02 (s, 2H), 2.27 (s, 3H).

**6-bromo-3-((3,4-dichlorobenzyl)oxy)-2-methylphenol** (**I4**). The same reaction for compound **I3** was used, and the coarse product was used without further purification.

**1-bromo-4-((3,4-dichlorobenzyl)oxy)-2-(methoxymethoxy)-3-methylbenzene** (**I5**). To the solution of compound **I4** (approximate 5 mmol, coarse, 1 eq) in 50 mL DCM, bromo(methoxy)methane (1249.6 mg, 10 mmol, 2 eq) and N,N-Diisopropylethylamine (DIPEA, 1938.6 mg, 15 mmol, 3 eq) were added. The mixture was stirred at 50 ^o^C for 3 h and monitored by TLC. After being cooled to room temperature, the solvent was removed by evaporation in vacuum, and purified by the Combi-Flash system (A = hexane, B = ethyl acetate; B%: 0% to 15%, 32 min) to give **I5** as an oily white solid (124.6 mg, 3-step yield: 6.1%). ^1^H NMR (500 MHz, CDCl_3_) δ 7.67 (d, *J* = 2.1 Hz, 1H), 7.49 (d, *J* = 8.2 Hz, 1H), 7.42 – 7.31 (m, 2H), 6.84 (d, *J* = 8.9 Hz, 1H), 5.21 (s, 2H), 4.88 (s, 2H), 3.51 (s, 3H), 2.24 (s, 3H). MS: m/z calcd for [M + H]^+^ 404.9582, found 405.17.

**4-((3,4-dichlorobenzyl)oxy)-2-(methoxymethoxy)-3-methyl-1,1'-biphenyl** (**I6**). To the solution of **I5** (459.7 mg, 1.13 mmol, 1 eq) in toluene/H_2_O/EtOH mixture (4/2/1, v/v/v), phenylboronic acid (206.7 mg, 1.70 mmol, 1.5 eq), Ph(PPh_3_)_4_ (130.6 mg, 0.11 mmol, 0.1 eq), and K_2_CO_3_ (624.7 mg, 4.52 mmol, 4 eq) were added successively. The reaction mixture was stired at 90 ^o^C under N_2_ overnight and monitored by TLC. After being cooled to room temperature, the mixture was extracted with CH_2_Cl_2_ (3 x 40 mL), and the combined organic phase was dried via MgSO_4_ and concentrated under vacuum. The crude product was purified by the Combi-Flash system (A = hexane, B = ethyl acetate; B%, 0% to 15%, 32 min) to give I**6** as a white solid (287.9 mg, 63.7%). ^1^H NMR (500 MHz, CDCl_3_) δ 7.55 (d, *J* = 7.6 Hz, 2H), 7.42 (t, *J* = 7.5 Hz, 2H), 7.35 (dd, *J* = 10.2, 7.8 Hz, 2H), 7.19 (d, *J* = 7.7 Hz, 2H), 7.00 (d, *J* = 8.5 Hz, 1H), 6.93 (d, *J* = 8.2 Hz, 1H), 5.28 (s, 2H), 4.39 (s, 2H), 3.55 (s, 3H), 2.27 (s, 3H). MS: m/z calcd for [M + H]^+^ 403.0789, found 403.20.

**4-((3,4-dichlorobenzyl)oxy)-3-methylbenzaldehyde** (**I7**). The same reaction for compound **I3** was used, and **I7** was obtained as white solid (1231.0 mg, 100%). ^1^H NMR (500 MHz, CDCl_3_) δ 9.90 (s, 1H), 7.78 – 7.70 (m, 2H), 7.56 (s, 1H), 7.51 (d, *J* = 8.3 Hz, 1H), 7.30 (d, *J* = 8.5 Hz, 1H), 6.96 (d, *J* = 8.3 Hz, 1H), 5.14 (s, 2H), 2.36 (s, 3H). MS: m/z calcd for [M - H]^-^ 293.0214, found 293.31.

**4-(benzyloxy)-2-hydroxy-3-methylbenzaldehyde** (**I8**). The same reaction for compound **I3** was used, and **I8** was obtained as white solid (1093.0 mg, 90.2%). ^1^H NMR (500 MHz, CDCl_3_) δ 11.49 (s, *J* = 1.0 Hz, 1H), 9.73 (s, 1H), 7.48 – 7.40 (m, 4H), 7.37 (dd, *J* = 8.0, 5.7 Hz, 2H), 6.63 (d, *J* = 8.6 Hz, 1H), 5.21 (s, 2H), 2.20 (s, 3H). MS: m/z calcd for [M + H]^+^ 243.0943, found 243.12.

**4-(benzyloxy)-2-hydroxy-3-methylbenzaldehyde** (**I9**). The same reaction for compound **I5** was used, and **I9** was obtained as white solid (1006.0 mg, 78.1%). ^1^H NMR (500 MHz, CDCl_3_) δ 10.22 (s, 1H), 7.75 (d, *J* = 8.7 Hz, 1H), 7.44 (m, 4H), 7.40 – 7.34 (m, 1H), 6.85 (d, *J* = 8.7 Hz, 1H), 5.19 (s, 2H), 5.11 (s, 2H), 3.63 (s, 3H), 2.27 (s, 3H). MS: m/z calcd for [M + H]^+^ 287.1205, found 287.14.

**(*E*)-3-(4-((3,4-dichlorobenzyl)oxy)-3-methylphenyl)acrylaldehyde** (**I10**). To the solution of **I7** (590.3 mg, 2 mmol, 1 eq) in THF (anhydrous, 50 mL), (1,3-Dioxolan-2-ylmethyl)triphenylphosphonium bromide (1293.9 mg, 3 mmol, 1.5 eq), NaH (240 mg, 10 mmol, 5 eq), and 18-crown-6 (50 mg) were added successively. The mixture was stirred at room temperature overnight and monitored by TLC. 1 M HCl aqueous solution (20 mL) was added and maintained at room temperature for 30 min. The reaction system was then extracted by CH_2_Cl_2_ (3 x 40 mL), and the combined organic phase was dried via MgSO_4_ and concentrated under vacuum. The crude product was purified by the Combi-Flash system (A = hexane, B = ethyl acetate; B%, 0% to 25%, 32 min ) to give **I7** as a yellow solid (290.0 mg, 45.1%). ^1^H NMR (500 MHz, CDCl_3_) δ 9.68 (d, *J* = 7.8 Hz, 1H), 7.56 (s, 1H), 7.50 (d, *J* = 8.2 Hz, 1H), 7.46 – 7.38 (m, 3H), 7.32 – 7.28 (m, 1H), 6.88 (d, *J* = 8.4 Hz, 1H), 6.64 (dd, *J* = 15.8, 7.7 Hz, 1H), 5.11 (s, 2H), 2.33 (s, 3H). MS: m/z calcd for [M + H]^+^ 321.0, found 321.1.

**(*E*)-3-(4-(benzyloxy)-2-(methoxymethoxy)-3-methylphenyl)acrylaldehyde** (**I11**). The same reaction for compound **I10** was used, and **I11** was used in the next step without further purification.

***N*'-((1E,2E)-3-(4-((3,4-dichlorobenzyl)oxy)-3-methylphenyl)allylidene)-4-methylbenzenesulfonohydrazide (I12)**. To the solution of **I10** (279.0 mg, 0.87 mmol, 1 eq) in MeOH (50 mL) was added 4-methylbenzenesulfonohydrazide (356.4 mg, 1.9 mmol, 2 eq). The reaction was stirred at 60 ^o^C for 3 h and monitored by TLC. After cooling to room temperature, the solvent was removed by evaporation in vacuum and the residue was purified by the Combi-Flash system (A = hexane, B = ethyl acetate; B%, 0% to 35 %, 32 min) to give **I12** as a pale yellow oil (301.7 mg, 71.3%). ^1^H NMR (500 MHz, DMSO-*d*_6_) δ 7.74 – 7.65 (m, 5H), 7.48 – 7.40 (m, 4H), 7.35 (d, *J* = 8.6 Hz, 1H), 6.97 (d, *J* = 8.5 Hz, 1H), 6.85 (d, *J* = 16.0 Hz, 1H), 6.70 (dd, *J* = 16.0, 9.2 Hz, 1H), 5.18 (s, 2H), 2.39 (s, 3H), 2.20 (s, 3H). MS: m/z calcd for [M + H]+ 489.1, found 489.2.

***N*'-((1E,2E)-3-(4-(benzyloxy)-2-(methoxymethoxy)-3-methylphenyl)allylidene)-4-methylbenzenesulfonohydrazide** (**I13**). The same reaction for compound **I12** was used, and **I13** was obtained as yellow oil (377.0 mg, 58.2%). ^1^H NMR (500 MHz, CDCl_3_) δ 7.88 – 7.84 (m, 2H), 7.44 – 7.33 (m, 10H), 6.75 (dd, *J* = 9.0, 2.5 Hz, 2H), 5.12 (s, 2H), 4.95 (s, 2H), 3.58 (s, 3H), 2.44 (s, 3H), 2.24 (s, 3H). MS: m/z calcd for [M + H]^+^ 481.1719, found 481.73.

***N*'-((1E,2E)-3-(4-(benzyloxy)-2-(methoxymethoxy)-3-methylphenyl)allylidene)-4-methylbenzenesulfonohydrazide** (**I14**). To the solution of **I13** (377.0 mg, 0.79 mmol, 1 eq) in THF (50 mL) was added CsF (360.0 mg, 2.37 mmol, 3 eq) and benzyltriethylammonium chloride (TEBAC, 45.0 mg, 0.20 mmol, 0.25 eq) were added. The reaction was stirred at 70 ^o^C overnight and monitored by TLC. After being cooled to room temperature, the solvent was removed by evaporation under vacuum and the residue was purified by the Combi-Flash system (A = hexane, B = ethyl acetate; B%, 0% to 40%, 32 min) to give **I14** as a yellow oil (146.0 mg, 57.0%). ^1^H NMR (500 MHz, CDCl_3_) δ 7.66 (s, 1H), 7.48 (d, *J* = 7.4 Hz, 2H), 7.43 (t, *J* = 7.7 Hz, 3H), 7.37 (t, *J* = 7.4 Hz, 1H), 6.78 (d, *J* = 8.6 Hz, 1H), 6.59 (s, 1H), 5.11 (s, 2H), 4.94 (s, 2H), 3.46 (s, 3H), 2.35 (s, 3H). MS: m/z calcd for [M + H]^+^ 325.1474, found 325.72.

**2,4-dihydroxy-3-methylbenzoic acid** (**I15**). To the solution of 2-methylbenzene-1,3-diol (2482.0 mg, 20 mmol, 1 eq) in acetonitrile (ACN, 100 mL), diazabicycloundecene (DBU, 9132.0 mg, 60 mmol, 3 eq) was added. The reaction mixture was stirred at r.t. under CO_2_ for 2 d and monitored by TLC. After the starting reagent was consumed, the mixture was filtered to remove solid impurity and the same reaction for compound **I12** was used and concentrated under vacuum. The crude product was used for the next step without further purification.

**3-(4-((3,4-dichlorobenzyl)oxy)-3-methylphenyl)-1H-pyrazole** (**D1**). The same reaction for compound **I14** was used, and **D1** was obtained as yellow oil (377.0 mg, 58.2%). ^1^H NMR (500 MHz, CDCl_3_) δ 7.58 (d, *J* = 9.4 Hz, 2H), 7.55 – 7.47 (m, 2H), 7.41 (d, *J* = 10.0 Hz, 1H), 7.21 (d, *J* = 10.0 Hz, 1H), 6.75 (d, *J* = 8.6 Hz, 1H), 6.52 (s, 1H), 4.91 (s, 2H), 2.29 (s, 3H). ^13^C NMR (126 MHz, CDCl_3_) δ 156.58, 147.97, 137.49, 133.96, 132.61, 131.72, 130.53, 128.86, 127.34, 126.30, 124.78, 111.39, 102.19, 68.33, 60.53, 16.45, 14.27. HRMS: m/z calcd for [M + H]^+^ 333.0483, found 333.19.

**4-((3,4-dichlorobenzyl)oxy)-3-methyl-1,1'-biphenyl** (**D2**). To the solution of **I3** (173.0 mg, 0.5 mmol, 1 eq) in toluene/H_2_O/EtOH mixture (4/2/1, v/v/v), phenylboronic acid (91.4 mg, 0.75 mmol,1.5 eq), Ph(PPh_3_)_4_ (57.8 mg, 0.05 mmol, 0.1 eq), and K_2_CO_3_ (276.4 mg, 2 mmol, 4 eq) were added successively. The reaction mixture was stired at 90 ^o^C under N_2_ overnight and monitored by TLC. After being cooled to room temperature, the mixture was extracted with CH_2_Cl_2_ (3 x 40 mL), and the combined organic phase was dried via MgSO_4_ and concentrated under vacuum. The crude product was purified by the Combi-Flash system (A = hexane, B = DCM; B%, 0% to 15%, 32 min ) to give **D2** as a white solid (58.2 mg, 33.9%). ^1^H NMR (500 MHz, CDCl_3_) δ 7.61 – 7.56 (m, 3H), 7.50 (d, *J* = 8.2 Hz, 1H), 7.48 – 7.42 (m, 3H), 7.41 (dd, *J* = 8.4, 2.4 Hz, 1H), 7.37 – 7.30 (m, 2H), 6.92 (d, *J* = 8.4 Hz, 1H), 5.09 (s, 2H), 2.39 (s, 3H). ^13^C NMR (126 MHz, CDCl_3_) δ 156.01, 140.84, 137.69, 134.12, 132.74, 131.81, 130.60, 129.79, 128.99, 128.73, 127.37, 126.81, 126.73, 126.31, 125.43, 111.58, 68.59, 16.59. HRMS: m/z calcd for [M - H]^-^ 341.0578, found 341.42.

**4-(4-((3,4-dichlorobenzyl)oxy)-3-methylphenyl)pyridine** (**D3**). The same reaction for compound **D2** was used, and **D3** was obtained as white solid (83.7 mg, 24.3%). ^1^H NMR (500 MHz, Chloroform-*d*) δ 8.64 (d, *J* = 5.2 Hz, 2H), 7.58 (s, 1H), 7.52 – 7.40 (m, 5H), 7.31 (dd, *J* = 8.2, 2.0 Hz, 1H), 6.93 (d, *J* = 8.4 Hz, 1H), 5.10 (s, 2H), 2.38 (s, 3H). ^13^C NMR (126 MHz, CDCl_3_) δ 157.32, 150.19, 147.84, 137.29, 132.81, 131.97, 130.66, 129.51, 128.97, 127.89, 126.28, 125.54, 121.14, 111.65, 68.56, 30.34, 16.58. HRMS: m/z calcd for [M + H]^+^ 344.0531, found 344.24.

**3-(benzyloxy)-2-methyl-6-(1H-pyrazol-3-yl)phenol** (**D4**). To the solution of **I14** (146.0 mg, 0.45 mmol, 1 eq) in 12 mL DCM, trifluoroacetic acid (TFA, 12 mL) was added. The reaction mixture was stired at r.t. for 1 h and monitored by TLC. The solvent was removed by evaporation under vacuum, and the coarse product was purified by the Combi-Flash system (A = hexane, B = ethyl acetate; B%, 0% to 25%, 32 min) to give **D4** as a white solid (128.1 mg, 100%). ^1^H NMR (500 MHz, CDCl_3_) δ 7.61 (d, *J* = 2.6 Hz, 1H), 7.49 (d, *J* = 7.1 Hz, 2H), 7.46 – 7.40 (m, 3H), 7.39 – 7.33 (m, 1H), 6.65 (d, *J* = 2.6 Hz, 1H), 6.59 (d, *J* = 8.6 Hz, 1H), 5.15 (s, 2H), 2.30 (s, 3H). 13C NMR (126 MHz, CDCl_3_) δ 157.67, 154.83, 151.68, 137.47, 129.24, 128.55, 127.79, 127.15, 124.19, 114.01, 110.13, 103.65, 101.74, 70.22, 8.57. HRMS: m/z calcd for [M + H]^+^ 281.1212, found 281.71.

**4-((3,4-dichlorobenzyl)oxy)-3-methyl-[1,1'-biphenyl]-2-ol** (**D5**). The same reaction for compound **D4** was used, and **D5** was obtained as white solid (54.3 mg, 21.3%). ^1^H NMR (500 MHz, CDCl_3_) δ 7.54 (dd, *J* = 8.3, 1.4 Hz, 2H), 7.43 (dd, *J* = 8.3, 6.7 Hz, 2H), 7.39 – 7.32 (m, 2H), 7.19 (d, *J* = 2.1 Hz, 1H), 7.12 (d, *J* = 8.4 Hz, 1H), 6.93 (dd, *J* = 8.1, 2.0 Hz, 1H), 6.71 (d, *J* = 8.3 Hz, 1H), 5.09 (s, 1H), 4.39 (s, 2H), 2.27 (s, 3H). ^13^C NMR (126 MHz, CDCl3) δ 154.93, 154.29, 138.68, 137.19, 132.31, 131.95, 130.24, 130.22, 129.32, 128.39, 128.29, 128.06, 127.48, 126.96, 117.98, 111.39, 73.19, 9.23. HRMS: m/z calcd for [M + H]^+^ 359.0527, found 359.14.

**3-(4-((3,4-dichlorobenzyl)oxy)-3-methylphenyl)-1H-pyrazole** (**D6**). The same reaction for compound **I3** was used, and **D6** was obtained as white solid (481.3 mg, 66.3%). ^1^H NMR (500 MHz, CDCl_3_) δ 10.97 (s, 1H), 10.43 (s, 1H), 7.75 (d, *J* = 2.0 Hz, 1H), 7.67 (d, *J* = 8.3 Hz, 1H), 7.59 (d, *J* = 8.8 Hz, 1H), 7.48 (dd, *J* = 8.3, 2.0 Hz, 1H), 6.48 (d, *J* = 8.8 Hz, 1H), 5.35 (s, 2H), 2.00 (s, 3H). ^13^C NMR (126 MHz, DMSO-*d*_6_) δ 169.96, 162.61, 161.37, 137.49, 131.63, 131.25, 130.46, 128.74, 128.66, 110.90, 108.02, 103.57, 65.08, 30.88, 8.32. HRMS: m/z calcd for [M + H]^+^ 327.0113, found 327.19.

**Table S1.** Structures of candidate compounds obtained from affinity selection mass spectrometry (ASMS) screening and commercial source.

| **Compound ID** | **SMILES** | **Vendor** | **Reference** |
| --- | --- | --- | --- |
| **1** | Clc1ccc(cc1)c2nc(c([nH]2)c3ccc(Cl)cc3)c4ccc(Cl)cc4 | Maybridge | DP01606 |
| **2** | CSc1nc(-c2ccc(Cl)cc2)nc(N)c1-c1ccccc1 | Maybridge | EN00095 |
| **3** | Clc1cccc(Cl)c1/C=C/c1cc[nH]n1 | Maybridge | CD09853 |
| **4** | Clc1cccc(Cl)c1CSc1nc2ccccc2n2cccc12 | Maybridge | CD03388 |
| **5** | Cc1ccc(-c2nc(NC(=O)Nc3c(-c4ccccc4)noc3C)sc2C)cc1Cl | Maybridge | HTS06895 |
| **6** | Cc1c(-c2cnc(NC(=O)c3ccccc3)s2)sc2ccc(Cl)cc12 | Maybridge | MWP00628 |
| **7** | Cc1ccccc1Cn1c(-c2cncs2)nc2ccccc21 | TimTec | HTS13723 |
| **8** | CC(C)(c1ccc(OC(=O)c2ccco2)cc1)c1ccc(OC(=O)c2ccco2)cc1 | ChemDiv | Y513-0428 |
| **9** | Cc1c(OCc2ccc(Cl)c(Cl)c2)ccc(-c2cc[nH]n2)c1O | Maybridge | RH00458 |
| **10** | O=C1NN(c2ccccc2)C(SCc2ccccc2Cl)=Nc2ccccc21 | Maybridge | DP01855 |
| **11** | Oc1c(Cl)cc(Cl)cc1C=NNC(=S)NCC2CCCO2 | Maybridge | RDR00929 |
| **12** | CC(C)(C)c1ccc(C(=O)Nc2ccc3oc(-c4ccccc4)nc3c2)cc1 | ChemDiv | 8010-1349 |
| **13** | COc1ccc(O)c(-c2nc3c(C(C)(C)C)cc(C(C)(C)C)c(O)c3o2)c1 | ChemDiv | 3093-0027 |
| **14** | Cc1cc(Nc2ccc(Br)cc2)nc(-c2ccccc2O)n1 | ChemDiv | 4229-0028 |
| **15** | Cc1nc(-c2ccccn2)nc(N2CCc3ccccc3C2)c1Cl | Maybridge | HTS13930 |
| **16** | O=C(Nc1cccc(-c2nc3ccccc3s2)c1)c1cccc2ccccc12 | ChemDiv | 8495-0305 |
| **17** | Cc1cccc(NC(=O)c2cc(-c3ccccc3)nc3c(C)cc(C)cc23)c1 | Specs | AG-690/13704827 |
| **18** | Cc1ccc(-c2cc(-c3csc(COC(=O)C(C)(C)C)n3)on2)cc1C | Maybridge | SPB06658 |
| **19** | COc1cc(OC)cc(-c2nc(N)c3c(C)c(-c4ccc(OC)c(OC)c4)sc3n2)c1 | Maybridge | CD02075 |
| **20** | CSc1cc(-c2ccc(Cl)cc2)nc(-c2ccc(Cl)cc2)c1 | Maybridge | EN00242 |
| **21** | CC1=CC=C(C=C1)C2=CC(C3=CC=CC=C3)=NC(C4=CC=CC=C4)=C2 | Maybridge | NRB03265 |
| **22** | Cc1cccc(C(=O)Nc2nc(-c3ccccc3)c(-c3ccccc3)s2)c1 | TimTec | HTS05650 |

**Table S2.** A list of hydrogen bonds formed between the -OH group of **9** and involved amino acid of ICOS obtained based on 10,000 structures generated during 100 ns of MD simulations for the selected consensus docked poses together with the final r.m.s.d. Values were computed for the position of the heavy atoms of the central ring of **9**. Only intermolecular hydrogen bonds that appear in over 10% are provided.

| **Binding site and Pose** | **Docking H-bond** | **r.m.s.d. (Å)** | **H- bond** | **Population** | **DAD (Å)** | **D-H-A Angle (º)** |
| --- | --- | --- | --- | --- | --- | --- |
| **β1** | Ser91:OG | 17.1 | Ser85:O | 21.6% | 2.7 | 157.2 |
| **δ1** | Tyr97:OH | 5.3 | - | - | - | - |
| **δ2** | Gln86:OE1 | 3.8 | - | - | - | - |
| **δ3** | Gln39:OE1 | 7.5 | - | - | - | - |
| **δ4** | Gln39:OE1 | 4.9 | Gln39:NE2 | 17.9% | 3.0 | 146.2 |
| **δ5** | Tyr97:OH | 3.0 | - | - | - | - |
| **φ1** | Gln54:NE2 | 4.4 | Gln54:NE2 | 17.8% | 3.0 | 158.0 |
|  |  |  | Gln54:OE1 | 69.6% | 2.7 | 149.9 |
| **φ2** | NAG1:O2N | 2.8 | Gln54:NE2 | 12.2% | 3.0 | 152.0 |

**Table S3.** The population of hydrogen bonds created between 9 and the Gln54 of φ site of ICOS during 100 ns of MD simulations, together with the final r.m.s.d. values computed for the position of the heavy atoms of the central ring of 9. Averaged values were obtained from 10,000 structures generated in each replica.

| **Replica** | **r.m.s.d. (Å)** | **E_int_ (kcal/mol)** | **Hydrogen bond** | **Population** | **DAD (Å)** | **D-H-A Angle (º)** |
| --- | --- | --- | --- | --- | --- | --- |
| **1** | 4.4 | -49.0 ± 9.9 | Gln54:OE1 | 69.6% | 3.0 ± 0.5 | 112.0 ± 42.3 |
| **2** | 4.1 | -56.1 ± 9.2 | Gln54:OE1 | 85.9% | 2.8 ± 0.2 | 116.0 ± 44.1 |
| **3** | 3.7 | -53.1 ± 10.0 | Gln54:OE1 | 86.4% | 2.8 ± 0.3 | 130.1 ± 37.2 |

**Table S4.** Interaction energy (E_int_) computed between **9** compound and glycan and protein residues. All 30,000 structures generated in three replicas of MD simulations were included.

| **9** | | | | | | | |
| --- | --- | --- | --- | --- | --- | --- | --- |
| **Residue** | **E_Elec_ (kcal/mol)** | **E_vdW_ (kcal/mol)** | **E_int_ (kcal/mol)** | **Residue** | **E_Elec_ (kcal/mol)** | **E_vdW_ (kcal/mol)** | **E_int_ (kcal/mol)** |
| Ile22 | -0.22 ± 0.54 | -0.08 ± 0.30 | -0.30 ± 0.77 | Leu80 | 0.13 ± 0.11 | -0.04 ± 0.05 | 0.09 ± 0.11 |
| Gln23 | 0.00 ± 0.15 | -0.02 ± 0.10 | -0.03 ± 0.18 | Lys81 | 0.72 ± 0.72 | -0.03 ± 0.02 | 0.70 ± 0.72 |
| Gly24 | 0.00 ± 0.03 | 0.00 ± 0.01 | 0.00 ± 0.03 | Phe82 | 0.00 ± 0.00 | 0.00 ± 0.00 | 0.00 ± 0.00 |
| Ser25 | 0.00 ± 0.01 | 0.00 ± 0.00 | 0.00 ± 0.01 | Cys83 | 0.00 ± 0.00 | 0.00 ± 0.00 | 0.00 ± 0.00 |
| Ala26 | 0.00 ± 0.00 | 0.00 ± 0.00 | 0.00 ± 0.00 | His84 | 0.00 ± 0.00 | 0.00 ± 0.00 | 0.00 ± 0.00 |
| Asn27 | 0.00 ± 0.00 | 0.00 ± 0.00 | 0.00 ± 0.00 | Ser85 | 0.00 ± 0.00 | 0.00 ± 0.00 | 0.00 ± 0.00 |
| Tyr28 | 0.00 ± 0.00 | 0.00 ± 0.00 | 0.00 ± 0.00 | Gln86 | 0.00 ± 0.00 | 0.00 ± 0.00 | 0.00 ± 0.00 |
| Glu29 | 0.00 ± 0.00 | 0.00 ± 0.00 | 0.00 ± 0.00 | Leu87 | 0.00 ± 0.00 | 0.00 ± 0.00 | 0.00 ± 0.00 |
| Met30 | 0.00 ± 0.01 | 0.00 ± 0.00 | 0.00 ± 0.01 | Ser88 | 0.00 ± 0.00 | 0.00 ± 0.00 | 0.00 ± 0.00 |
| Phe31 | 0.00 ± 0.00 | 0.00 ± 0.00 | 0.00 ± 0.00 | Asn89 | 0.00 ± 0.00 | 0.00 ± 0.00 | 0.00 ± 0.00 |
| Ile32 | 0.00 ± 0.00 | 0.00 ± 0.00 | 0.00 ± 0.00 | Asn90 | 0.00 ± 0.00 | 0.00 ± 0.00 | 0.00 ± 0.00 |
| Phe33 | 0.00 ± 0.00 | 0.00 ± 0.00 | 0.00 ± 0.00 | Ser91 | 0.00 ± 0.00 | 0.00 ± 0.00 | 0.00 ± 0.00 |
| His34 | 0.00 ± 0.00 | 0.00 ± 0.00 | 0.00 ± 0.00 | Val92 | 0.00 ± 0.00 | 0.00 ± 0.00 | 0.00 ± 0.00 |
| Asn35 | 0.00 ± 0.00 | 0.00 ± 0.00 | 0.00 ± 0.00 | Ser93 | 0.00 ± 0.00 | 0.00 ± 0.00 | 0.00 ± 0.00 |
| Gly36 | 0.00 ± 0.00 | 0.00 ± 0.00 | 0.00 ± 0.00 | Phe94 | 0.10 ± 0.04 | 0.00 ± 0.00 | 0.09 ± 0.04 |
| Gly37 | 0.00 ± 0.00 | 0.00 ± 0.00 | 0.00 ± 0.00 | Phe95 | 0.00 ± 0.00 | 0.00 ± 0.00 | 0.00 ± 0.00 |
| Val38 | 0.00 ± 0.00 | 0.00 ± 0.00 | 0.00 ± 0.00 | Leu96 | 0.00 ± 0.00 | 0.00 ± 0.00 | 0.00 ± 0.00 |
| Gln39 | 0.00 ± 0.00 | 0.00 ± 0.00 | 0.00 ± 0.00 | Tyr97 | 0.00 ± 0.00 | 0.00 ± 0.00 | 0.00 ± 0.00 |
| Ile40 | 0.00 ± 0.00 | 0.00 ± 0.00 | 0.00 ± 0.00 | Asn98 | 0.00 ± 0.00 | 0.00 ± 0.00 | 0.00 ± 0.00 |
| Leu41 | 0.00 ± 0.00 | 0.00 ± 0.00 | 0.00 ± 0.00 | Leu99 | 0.00 ± 0.00 | 0.00 ± 0.00 | 0.00 ± 0.00 |
| Cys42 | 0.00 ± 0.00 | 0.00 ± 0.00 | 0.00 ± 0.00 | Asp100 | 0.00 ± 0.03 | 0.00 ± 0.00 | 0.00 ± 0.03 |
| Lys43 | 0.00 ± 0.00 | 0.00 ± 0.00 | 0.00 ± 0.00 | His101 | 0.00 ± 0.00 | 0.00 ± 0.00 | 0.00 ± 0.00 |
| Tyr44 | 0.00 ± 0.00 | 0.00 ± 0.00 | 0.00 ± 0.00 | Ser102 | 0.00 ± 0.01 | 0.00 ± 0.00 | 0.00 ± 0.01 |
| Pro45 | 0.00 ± 0.00 | 0.00 ± 0.00 | 0.00 ± 0.00 | His103 | 0.08 ± 0.14 | -0.02 ± 0.04 | 0.06 ± 0.11 |
| Asp46 | 0.00 ± 0.00 | 0.00 ± 0.00 | 0.00 ± 0.00 | Ala104 | 0.01 ± 0.07 | 0.00 ± 0.01 | 0.01 ± 0.07 |
| Ile47 | 0.00 ± 0.00 | 0.00 ± 0.00 | 0.00 ± 0.00 | Asn105 | -0.02 ± 0.07 | 0.00 ± 0.00 | -0.02 ± 0.07 |
| Val48 | 0.00 ± 0.00 | 0.00 ± 0.00 | 0.00 ± 0.00 | Tyr106 | 0.10 ± 0.18 | -0.07 ± 0.07 | 0.02 ± 0.18 |
| Gln49 | 0.00 ± 0.00 | 0.00 ± 0.00 | 0.00 ± 0.00 | Tyr107 | 0.03 ± 0.12 | -0.02 ± 0.02 | 0.00 ± 0.14 |
| Gln50 | 0.00 ± 0.01 | 0.00 ± 0.00 | 0.00 ± 0.01 | Phe108 | -0.24 ± 0.26 | -0.76 ± 0.60 | -1.00 ± 0.58 |
| Phe51 | 0.05 ± 0.09 | 0.00 ± 0.00 | 0.05 ± 0.09 | Cys109 | 0.25 ± 0.17 | -0.05 ± 0.02 | 0.21 ± 0.17 |
| Lys52 | -6.79 ± 8.49 | -0.68 ± 0.67 | -7.47 ± 8.69 | Asn110 | 1.04 ± 0.81 | -1.05 ± 0.25 | -0.01 ± 0.82 |
| Met53 | 0.08 ± 0.14 | -0.04 ± 0.01 | 0.03 ± 0.14 | Leu111 | -0.08 ± 0.09 | -0.03 ± 0.01 | -0.12 ± 0.09 |
| Gln54 | -7.90 ± 2.58 | -0.17 ± 1.25 | -8.07 ± 1.98 | Ser112 | 0.11 ± 0.59 | -0.11 ± 0.04 | 0.00 ± 0.58 |
| Leu55 | -0.26 ± 0.19 | -0.09 ± 0.03 | -0.35 ± 0.19 | Ile113 | 0.01 ± 0.07 | 0.00 ± 0.00 | 0.01 ± 0.06 |
| Leu56 | 0.04 ± 0.35 | -2.32 ± 0.69 | -2.28 ± 0.82 | Phe114 | -0.14 ± 0.08 | -0.05 ± 0.04 | -0.19 ± 0.10 |
| Lys57 | -0.08 ± 0.35 | -0.21 ± 0.26 | -0.30 ± 0.46 | Asp115 | 0.00 ± 0.00 | 0.00 ± 0.00 | 0.00 ± 0.00 |
| Gly58 | -0.13 ± 0.85 | -0.81 ± 0.51 | -0.94 ± 0.90 | Pro116 | 0.00 ± 0.00 | 0.00 ± 0.00 | 0.00 ± 0.00 |
| Gly59 | -0.44 ± 0.87 | -1.81 ± 0.87 | -2.25 ± 1.39 | Pro117 | 0.00 ± 0.00 | 0.00 ± 0.00 | 0.00 ± 0.00 |
| Gln60 | 0.06 ± 0.40 | -0.41 ± 0.24 | -0.35 ± 0.45 | Pro118 | 0.00 ± 0.00 | 0.00 ± 0.00 | 0.00 ± 0.00 |
| Ile61 | 0.21 ± 0.27 | -1.64 ± 0.58 | -1.43 ± 0.65 | Phe119 | -0.10 ± 0.11 | -0.05 ± 0.05 | -0.15 ± 0.14 |
| Leu62 | 0.15 ± 0.10 | -0.04 ± 0.03 | 0.11 ± 0.10 | Lys120 | -0.01 ± 0.04 | 0.00 ± 0.00 | -0.01 ± 0.04 |
| Cys63 | -0.33 ± 0.18 | -0.02 ± 0.01 | -0.35 ± 0.18 | Val121 | 0.00 ± 0.15 | -0.12 ± 0.04 | -0.12 ± 0.16 |
| Asp64 | -0.78 ± 1.23 | -0.10 ± 0.03 | -0.88 ± 1.23 | The122 | -0.17 ± 0.10 | -0.02 ± 0.01 | -0.19 ± 0.10 |
| Leu65 | -0.14 ± 0.08 | -0.01 ± 0.00 | -0.15 ± 0.08 | Leu123 | -0.01 ± 0.13 | -0.53 ± 0.45 | -0.53 ± 0.51 |
| Thr66 | 0.00 ± 0.14 | -0.02 ± 0.01 | -0.03 ± 0.14 | Thr124 | 0.02 ± 0.07 | -0.01 ± 0.01 | 0.01 ± 0.07 |
| Lys67 | 0.01 ± 0.02 | 0.00 ± 0.00 | 0.01 ± 0.02 | Gly125 | -0.03 ± 0.06 | -0.01 ± 0.01 | -0.03 ± 0.06 |
| Thr68 | 0.02 ± 0.06 | 0.00 ± 0.00 | 0.02 ± 0.06 | Gly126 | 0.01 ± 0.04 | 0.00 ± 0.00 | 0.01 ± 0.04 |
| Lys69 | 0.00 ± 0.01 | 0.00 ± 0.00 | 0.00 ± 0.01 | Tyr127 | 0.00 ± 0.01 | 0.00 ± 0.00 | 0.00 ± 0.01 |
| Gly70 | 0.00 ± 0.01 | 0.00 ± 0.00 | 0.00 ± 0.01 | Leu128 | 0.00 ± 0.00 | 0.00 ± 0.00 | 0.00 ± 0.00 |
| Ser71 | 0.00 ± 0.00 | 0.00 ± 0.00 | 0.00 ± 0.00 | His129 | 0.00 ± 0.02 | 0.00 ± 0.00 | 0.00 ± 0.02 |
| Gly72 | 0.00 ± 0.00 | 0.00 ± 0.00 | 0.00 ± 0.00 | Ile130 | 0.00 ± 0.02 | 0.00 ± 0.00 | 0.00 ± 0.02 |
| Asn73 | 0.00 ± 0.00 | 0.00 ± 0.00 | 0.00 ± 0.00 | NAG1 | -1.78 ± 2.04 | -9.55 ± 1.49 | -11.33 ± 2.68 |
| Thr74 | 0.00 ± 0.00 | 0.00 ± 0.00 | 0.00 ± 0.00 | NAG2 | -6.50 ± 2.66 | -5.12 ± 1.13 | -11.61 ± 3.11 |
| Val75 | 0.00 ± 0.00 | 0.00 ± 0.00 | 0.00 ± 0.00 | BMA3 | -0.58 ± 1.02 | -1.28 ± 1.53 | -1.86 ± 2.17 |
| Ser76 | 0.00 ± 0.00 | 0.00 ± 0.00 | 0.00 ± 0.00 | MAN4 | 0.17 ± 0.52 | -0.15 ± 0.21 | 0.01 ± 0.56 |
| Ile77 | 0.00 ± 0.00 | 0.00 ± 0.00 | 0.00 ± 0.00 | MAN5 | -0.42 ± 1.57 | -1.57 ± 2.20 | -2.00 ± 3.13 |
| Lys78 | 0.09 ± 0.27 | 0.00 ± 0.00 | 0.09 ± 0.26 | NAG201 | 0.00 ± 0.00 | 0.00 ± 0.00 | 0.00 ± 0.00 |
| Ser79 | 0.03 ± 0.06 | 0.00 ± 0.00 | 0.03 ± 0.06 |  |  |  |  |

**Table S5.** Interaction energy (E_int_) computed between **D1** compound and glycan and protein residues. All three replicas are included and only frames with r.m.s.d. below 5.5 Å were considered.

| **D1** | | | | | | | |
| --- | --- | --- | --- | --- | --- | --- | --- |
| **Residue** | **E_Elec_ (kcal/mol)** | **E_vdW_ (kcal/mol)** | **E_int_ (kcal/mol)** | **Residue** | **E_Elec_ (kcal/mol)** | **E_vdW_ (kcal/mol)** | **E_int_ (kcal/mol)** |
| Ile22 | -0.21 ± 0.57 | -0.10 ± 0.35 | -0.31 ± 0.88 | Leu80 | 0.26 ± 0.16 | -0.07 ± 0.08 | 0.20 ± 0.14 |
| Gln23 | 0.02 ± 0.15 | -0.01 ± 0.03 | 0.00 ± 0.14 | Lys81 | 0.34 ± 0.47 | -0.04 ± 0.02 | 0.31 ± 0.48 |
| Gly24 | 0.01 ± 0.03 | 0.00 ± 0.00 | 0.01 ± 0.03 | Phe82 | 0.00 ± 0.01 | 0.00 ± 0.00 | 0.00 ± 0.01 |
| Ser25 | 0.00 ± 0.01 | 0.00 ± 0.00 | 0.00 ± 0.00 | Cys83 | 0.00 ± 0.00 | 0.00 ± 0.00 | 0.00 ± 0.00 |
| Ala26 | 0.00 ± 0.00 | 0.00 ± 0.00 | 0.00 ± 0.00 | His84 | 0.00 ± 0.00 | 0.00 ± 0.00 | 0.00 ± 0.00 |
| Asn27 | 0.00 ± 0.00 | 0.00 ± 0.00 | 0.00 ± 0.00 | Ser85 | 0.00 ± 0.01 | 0.00 ± 0.00 | 0.00 ± 0.01 |
| Tyr28 | 0.00 ± 0.00 | 0.00 ± 0.00 | 0.00 ± 0.00 | Gln86 | 0.00 ± 0.00 | 0.00 ± 0.00 | 0.00 ± 0.00 |
| Glu29 | 0.00 ± 0.00 | 0.00 ± 0.00 | 0.00 ± 0.00 | Leu87 | 0.00 ± 0.00 | 0.00 ± 0.00 | 0.00 ± 0.00 |
| Met30 | 0.00 ± 0.01 | 0.00 ± 0.00 | 0.00 ± 0.01 | Ser88 | 0.00 ± 0.00 | 0.00 ± 0.00 | 0.00 ± 0.00 |
| Phe31 | 0.00 ± 0.01 | 0.00 ± 0.00 | 0.00 ± 0.01 | Asn89 | 0.00 ± 0.00 | 0.00 ± 0.00 | 0.00 ± 0.00 |
| Ile32 | 0.00 ± 0.00 | 0.00 ± 0.00 | 0.00 ± 0.00 | Asn90 | 0.00 ± 0.00 | 0.00 ± 0.00 | 0.00 ± 0.00 |
| Phe33 | 0.00 ± 0.00 | 0.00 ± 0.00 | 0.00 ± 0.00 | Ser91 | 0.00 ± 0.00 | 0.00 ± 0.00 | 0.00 ± 0.00 |
| His34 | 0.00 ± 0.00 | 0.00 ± 0.00 | 0.00 ± 0.00 | Val92 | 0.00 ± 0.00 | 0.00 ± 0.00 | 0.00 ± 0.00 |
| Asn35 | 0.00 ± 0.00 | 0.00 ± 0.00 | 0.00 ± 0.00 | Ser93 | 0.00 ± 0.00 | 0.00 ± 0.00 | 0.00 ± 0.00 |
| Gly36 | 0.00 ± 0.00 | 0.00 ± 0.00 | 0.00 ± 0.00 | Phe94 | 0.06 ± 0.04 | 0.00 ± 0.00 | 0.06 ± 0.04 |
| Gly37 | 0.00 ± 0.00 | 0.00 ± 0.00 | 0.00 ± 0.00 | Phe95 | 0.00 ± 0.00 | 0.00 ± 0.00 | 0.00 ± 0.00 |
| Val38 | 0.00 ± 0.00 | 0.00 ± 0.00 | 0.00 ± 0.00 | Leu96 | 0.00 ± 0.00 | 0.00 ± 0.00 | 0.00 ± 0.00 |
| Gln39 | 0.00 ± 0.00 | 0.00 ± 0.00 | 0.00 ± 0.00 | Tyr97 | 0.00 ± 0.00 | 0.00 ± 0.00 | 0.00 ± 0.00 |
| Ile40 | 0.00 ± 0.00 | 0.00 ± 0.00 | 0.00 ± 0.00 | Asn98 | 0.00 ± 0.00 | 0.00 ± 0.00 | 0.00 ± 0.00 |
| Leu41 | 0.00 ± 0.00 | 0.00 ± 0.00 | 0.00 ± 0.00 | Leu99 | 0.00 ± 0.00 | 0.00 ± 0.00 | 0.00 ± 0.00 |
| Cys42 | 0.00 ± 0.00 | 0.00 ± 0.00 | 0.00 ± 0.00 | Asp100 | 0.00 ± 0.01 | 0.00 ± 0.00 | 0.00 ± 0.01 |
| Lys43 | 0.00 ± 0.00 | 0.00 ± 0.00 | 0.00 ± 0.00 | His101 | 0.00 ± 0.00 | 0.00 ± 0.00 | 0.00 ± 0.00 |
| Tyr44 | 0.00 ± 0.00 | 0.00 ± 0.00 | 0.00 ± 0.00 | Ser102 | 0.00 ± 0.00 | 0.00 ± 0.00 | 0.00 ± 0.00 |
| Pro45 | 0.00 ± 0.00 | 0.00 ± 0.00 | 0.00 ± 0.00 | His103 | 0.01 ± 0.03 | 0.00 ± 0.00 | 0.01 ± 0.03 |
| Asp46 | 0.00 ± 0.00 | 0.00 ± 0.00 | 0.00 ± 0.00 | Ala104 | 0.00 ± 0.03 | 0.00 ± 0.00 | 0.00 ± 0.03 |
| Ile47 | 0.00 ± 0.00 | 0.00 ± 0.00 | 0.00 ± 0.00 | Asn105 | 0.01 ± 0.12 | -0.01 ± 0.03 | 0.00 ± 0.11 |
| Val48 | 0.00 ± 0.00 | 0.00 ± 0.00 | 0.00 ± 0.00 | Tyr106 | 0.06 ± 0.17 | -0.08 ± 0.15 | -0.02 ± 0.19 |
| Gln49 | 0.00 ± 0.00 | 0.00 ± 0.00 | 0.00 ± 0.00 | Tyr107 | 0.02 ± 0.18 | -0.03 ± 0.03 | -0.01 ± 0.20 |
| Gln50 | 0.00 ± 0.01 | 0.00 ± 0.00 | 0.00 ± 0.02 | Phe108 | -0.01 ± 0.22 | -0.51 ± 0.57 | -0.52 ± 0.48 |
| Phe51 | 0.12 ± 0.10 | -0.01 ± 0.01 | 0.12 ± 0.10 | Cys109 | 0.11 ± 0.09 | -0.04 ± 0.02 | 0.07 ± 0.10 |
| Lys52 | 4.20 ± 1.44 | -0.85 ± 0.56 | 3.36 ± 1.15 | Asn110 | -0.03 ± 0.50 | -0.85 ± 0.28 | -0.88 ± 0.52 |
| Met53 | -0.08 ± 0.14 | -0.05 ± 0.02 | -0.12 ± 0.15 | Leu111 | 0.04 ± 0.06 | -0.03 ± 0.01 | 0.02 ± 0.06 |
| Gln54 | -0.29 ± 0.93 | -1.56 ± 0.61 | -1.84 ± 1.23 | Ser112 | 0.18 ± 0.10 | -0.11 ± 0.05 | 0.07 ± 0.10 |
| Leu55 | -0.16 ± 0.13 | -0.09 ± 0.03 | -0.25 ± 0.14 | Ile113 | -0.04 ± 0.04 | 0.00 ± 0.00 | -0.04 ± 0.04 |
| Leu56 | -0.17 ± 0.27 | -2.11 ± 0.83 | -2.28 ± 0.87 | Phe114 | -0.03 ± 0.09 | -0.06 ± 0.05 | -0.10 ± 0.13 |
| Lys57 | -0.11 ± 0.31 | -0.17 ± 0.28 | -0.27 ± 0.53 | Asp115 | 0.00 ± 0.00 | 0.00 ± 0.00 | 0.00 ± 0.00 |
| Gly58 | -0.02 ± 1.03 | -0.85 ± 0.56 | -0.87 ± 1.19 | Pro116 | 0.00 ± 0.00 | 0.00 ± 0.00 | 0.00 ± 0.00 |
| Gly59 | 0.10 ± 0.77 | -1.80 ± 0.90 | -1.71 ± 1.06 | Pro117 | 0.00 ± 0.00 | 0.00 ± 0.00 | 0.00 ± 0.00 |
| Gln60 | 0.06 ± 0.33 | -0.37 ± 0.25 | -0.31 ± 0.42 | Pro118 | 0.00 ± 0.00 | 0.00 ± 0.00 | 0.00 ± 0.00 |
| Ile61 | -0.03 ± 0.15 | -1.74 ± 0.75 | -1.77 ± 0.81 | Phe119 | -0.06 ± 0.06 | -0.08 ± 0.04 | -0.13 ± 0.08 |
| Leu62 | 0.10 ± 0.09 | -0.03 ± 0.02 | 0.07 ± 0.09 | Lys120 | -0.01 ± 0.03 | 0.00 ± 0.00 | -0.01 ± 0.03 |
| Cys63 | -0.17 ± 0.13 | -0.03 ± 0.02 | -0.20 ± 0.12 | Val121 | 0.04 ± 0.12 | -0.11 ± 0.04 | -0.07 ± 0.14 |
| Asp64 | -1.99 ± 0.90 | -0.09 ± 0.05 | -2.08 ± 0.94 | The122 | -0.07 ± 0.07 | -0.01 ± 0.01 | -0.08 ± 0.07 |
| Leu65 | -0.11 ± 0.10 | -0.01 ± 0.01 | -0.12 ± 0.11 | Leu123 | -0.01 ± 0.17 | -0.46 ± 0.39 | -0.48 ± 0.51 |
| Thr66 | -0.23 ± 0.11 | -0.04 ± 0.03 | -0.26 ± 0.12 | Thr124 | 0.00 ± 0.08 | -0.01 ± 0.01 | -0.01 ± 0.08 |
| Lys67 | 0.00 ± 0.03 | 0.00 ± 0.00 | 0.00 ± 0.03 | Gly125 | -0.03 ± 0.06 | -0.01 ± 0.01 | -0.03 ± 0.07 |
| Thr68 | 0.03 ± 0.10 | 0.00 ± 0.00 | 0.03 ± 0.10 | Gly126 | 0.02 ± 0.04 | 0.00 ± 0.00 | 0.02 ± 0.04 |
| Lys69 | 0.00 ± 0.02 | 0.00 ± 0.00 | 0.00 ± 0.02 | Tyr127 | -0.01 ± 0.02 | 0.00 ± 0.00 | -0.01 ± 0.02 |
| Gly70 | 0.00 ± 0.01 | 0.00 ± 0.00 | 0.00 ± 0.01 | Leu128 | 0.00 ± 0.01 | 0.00 ± 0.00 | 0.00 ± 0.01 |
| Ser71 | 0.00 ± 0.00 | 0.00 ± 0.00 | 0.00 ± 0.00 | His129 | 0.00 ± 0.00 | 0.00 ± 0.00 | 0.00 ± 0.00 |
| Gly72 | 0.00 ± 0.00 | 0.00 ± 0.00 | 0.00 ± 0.00 | Ile130 | 0.00 ± 0.00 | 0.00 ± 0.00 | 0.00 ± 0.00 |
| Asn73 | 0.00 ± 0.00 | 0.00 ± 0.00 | 0.00 ± 0.00 | NAG1 | -1.53 ± 1.53 | -8.91 ± 1.77 | -10.44 ± 2.69 |
| Thr74 | 0.00 ± 0.00 | 0.00 ± 0.00 | 0.00 ± 0.00 | NAG2 | -4.73 ± 2.11 | -5.76 ± 1.28 | -10.49 ± 2.75 |
| Val75 | 0.00 ± 0.00 | 0.00 ± 0.00 | 0.00 ± 0.00 | BMA3 | -0.66 ± 0.92 | -0.30 ± 0.27 | -0.96 ± 1.10 |
| Ser76 | 0.00 ± 0.00 | 0.00 ± 0.00 | 0.00 ± 0.00 | MAN4 | -0.02 ± 0.78 | -0.15 ± 0.27 | -0.17 ± 0.77 |
| Ile77 | 0.00 ± 0.00 | 0.00 ± 0.00 | 0.00 ± 0.00 | MAN5 | -0.67 ± 1.65 | -0.29 ± 0.43 | -0.96 ± 1.79 |
| Lys78 | 0.29 ± 0.52 | -0.01 ± 0.01 | 0.29 ± 0.51 | NAG201 | 0.00 ± 0.00 | 0.00 ± 0.00 | 0.00 ± 0.00 |
| Ser79 | 0.15 ± 0.14 | 0.00 ± 0.00 | 0.15 ± 0.14 |  |  |  |  |

**Table S6**. Cartesian coordinates of the six (α, β, γ, δ, ε, φ) putative binding sites obtained by clustering of blind docking results.

| **Site** | **Coordinates** | | |
| --- | --- | --- | --- |
|  | **X** | **Y** | **Z** |
| **α** | 38.523 | -43.976 | -13.836 |
| **β** | 39.870 | -45.139 | -3.863 |
| **γ** | 34.156 | -45.684 | 2.549 |
| **δ** | 47.235 | -56.860 | -7.006 |
| **ε** | 43.332 | -61.085 | -4.560 |
| **φ** | 30.002 | -58.699 | -11.718 |

**Table S7.** Missing atom types, point charges, and AMBER force field parameters assigned to **9** using GAFF2 and structures optimized AM1 level with bond charge corrections (AM1-BCC).

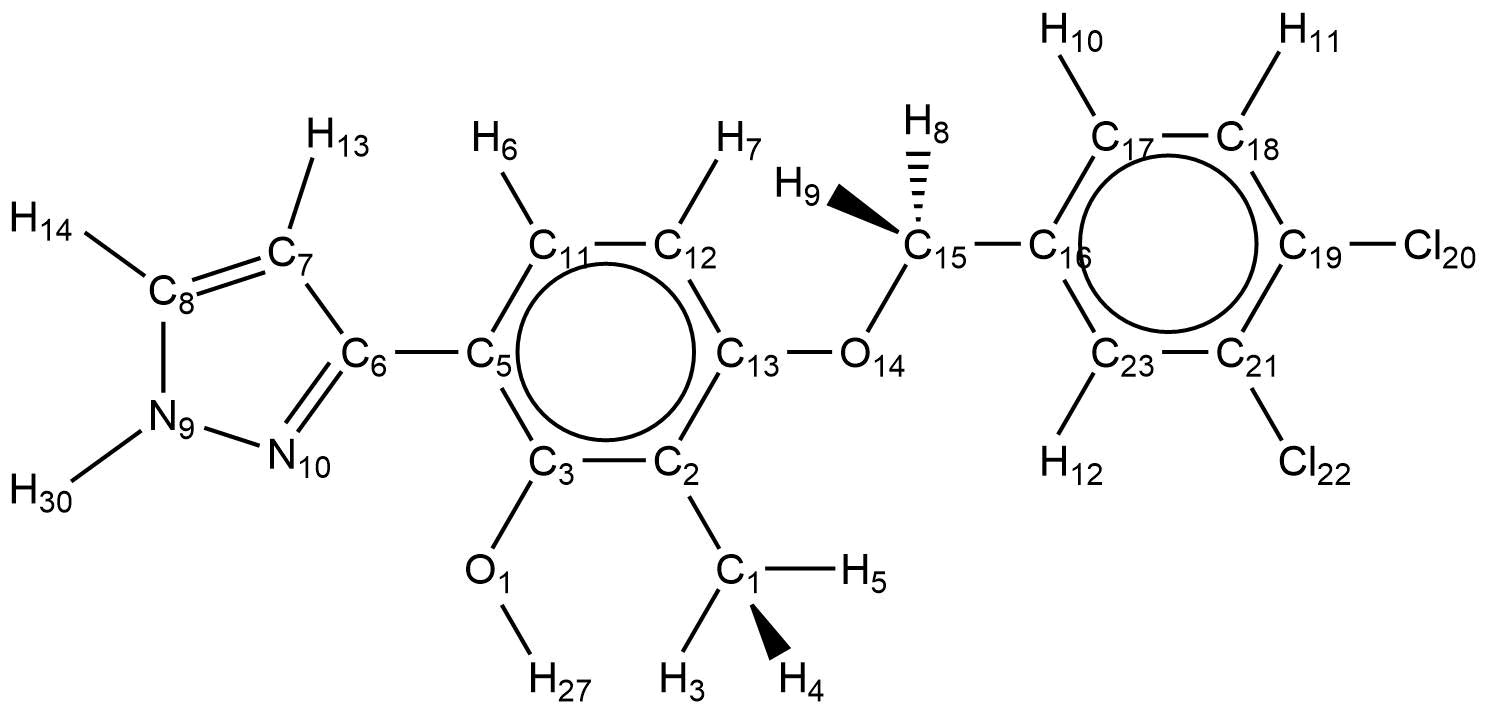

| **9** | | | | | |
| --- | --- | --- | --- | --- | --- |
| **Atom Name** | **Atom Type** | **Charge** | **Atom Name** | **Atom Type** | **Charge** |
| Cl22 | cl | -0.0674 | H7 | ha | 0.138 |
| C21 | ca | 0.0164 | C2 | ca | -0.1893 |
| C23 | ca | -0.092 | C1 | c3 | -0.0478 |
| H12 | ha | 0.168 | H3 | hc | 0.048367 |
| C19 | ca | 0.0184 | H4 | hc | 0.048367 |
| Cl20 | cl | -0.0674 | H5 | hc | 0.048367 |
| C18 | ca | -0.116 | C3 | ca | 0.1701 |
| H11 | ha | 0.153 | O1 | oh | -0.4941 |
| C17 | ca | -0.12 | H27 | ho | 0.423 |
| H10 | ha | 0.141 | C5 | ca | -0.1605 |
| C16 | ca | -0.0823 | C6 | cc | 0.3658 |
| C15 | c3 | 0.1747 | C7 | cc | -0.2583 |
| H8 | h1 | 0.0557 | H13 | ha | 0.179 |
| H9 | h1 | 0.0557 | C8 | cd | -0.1623 |
| O14 | os | -0.3319 | H14 | h4 | 0.174 |
| C13 | ca | 0.1661 | N9 | na | -0.0186 |
| C12 | ca | -0.24 | H30 | hn | 0.3187 |
| C11 | ca | -0.046 | N10 | nd | -0.5208 |
| H6 | ha | 0.152 |  |  |  |
| **Parameters** | | | | | |
| **BOND** | | | **ANGLE** | | |
| ca-cl | 305.6 | 1.75 | ca-ca-cl | 57.9 | 119.39 |
| ca-ca | 461.1 | 1.398 | ca-ca-ha | 48.2 | 119.88 |
| ca-ha | 345.8 | 1.086 | ca-ca-ca | 66.6 | 120.02 |
| c3-ca | 321 | 1.516 | c3-ca-ca | 63.5 | 120.77 |
| c3-h1 | 330.6 | 1.097 | ca-c3-h1 | 47 | 109.56 |
| c3-os | 308.6 | 1.432 | ca-c3-os | 68.3 | 108.95 |
| ca-os | 376.6 | 1.37 | c3-os-ca | 62.5 | 117.96 |
| c3-hc | 330.6 | 1.097 | h1-c3-h1 | 39.2 | 108.46 |
| ca-oh | 384 | 1.364 | h1-c3-os | 50.8 | 109.78 |
| ho-oh | 371.4 | 0.973 | ca-ca-os | 69.6 | 119.2 |
| ca-cc | 385.1 | 1.456 | ca-ca-cc | 65 | 120.79 |
| cc-cc | 419.8 | 1.428 | ca-c3-hc | 46.8 | 110.47 |
| cc-nd | 525.4 | 1.317 | ca-ca-oh | 69.5 | 119.9 |
| cc-ha | 349.1 | 1.084 | hc-c3-hc | 39.4 | 107.58 |
| cc-cd | 500.9 | 1.373 | ca-oh-ho | 49 | 108.58 |
| cd-h4 | 352 | 1.082 | ca-cc-cc | 67.2 | 111.04 |
| cd-na | 425.8 | 1.38 | ca-cc-nd | 67.6 | 123.24 |
| hn-na | 408.4 | 1.01 | cc-cc-ha | 47.1 | 121.07 |
| na-nd | 528.4 | 1.354 | cc-cc-cd | 68.2 | 114.19 |
|  | | | cc-nd-na | 74.6 | 103.82 |
|  | | | cc-cc-nd | 71.6 | 112.56 |
|  | | | cc-cd-h4 | 47.3 | 128.48 |
|  | | | cc-cd-na | 73.4 | 106.99 |
|  | | | cd-cc-ha | 48.5 | 121.76 |
|  | | | cd-na-hn | 46.8 | 125.5 |
|  | | | cd-na-nd | 70 | 112.36 |
|  | | | h4-cd-na | 49.8 | 120.53 |
|  | | | hn-na-nd | 49.9 | 119.55 |
| **IMPROPER** | | | **DIHEDRAL** | | |
| ca-ca-ca-cl 1.1 180.0 2.0 ca-ca-ca-ha 1.1 180.0 2.0  c3-ca-ca-ca 1.1 180.0 2.0 ca-ca-ca-os 1.1 180.0 2.0  ca-ca-ca-oh 1.1 180.0 2.0  ca-ca-ca-cc 1.1 180.0 2.0  ca-cc-cc-nd 1.1 180.0 2.0  cc-cd-cc-ha 1.1 180.0 2.0  cc-h4-cd-na 1.1 180.0 2.0  cd-hn-na-nd 1.1 180.0 2.0 | | | cl-ca-ca-ha 4 14.500 180.000 2.000 ca-ca-ca-cl 4 14.500 180.000 2.000 cl-ca-ca-cl 4 14.500 180.000 2.000 ca-ca-ca-ca 4 14.500 180.000 2.000 c3-ca-ca-ca 4 14.500 180.000 2.000 ca-ca-ca-ha 4 14.500 180.000 2.000 h1-c3-ca-ca 6 0.000 0.000 2.000 os-c3-ca-ca 6 0.000 0.000 2.000 c3-ca-ca-ha 4 14.500 180.000 2.000 ha-ca-ca-ha 4 14.500 180.000 2.000 ca-c3-os-ca 3 1.150 0.000 3.000 ca-ca-os-c3 2 1.800 180.000 2.000 cc-cd-na-hn 4 6.800 180.000 2.000 cc-cd-na-nd 4 6.800 180.000 2.000 ha-cc-cd-h4 4 16.000 180.000 2.000 ha-cc-cd-na 4 16.000 180.000 2.000 cd-na-nd-cc 2 9.600 180.000 2.000 h4-cd-na-hn 4 6.800 180.000 2.000 h4-cd-na-nd 4 6.800 180.000 2.000 hn-na-nd-cc 2 9.600 180.000 2.000 ha-cc-cc-nd 4 16.000 180.000 2.000 | | |
| **DIHEDRAL** | | |  |  |  |
| ca-ca-cc-nd 4 2.800 180.000 2.000  cc-ca-ca-ha 4 14.500 180.000 2.000 ca-ca-oh-ho 2 1.800 180.000 2.000 c3-ca-ca-oh 4 14.500 180.000 2.000 cc-ca-ca-oh 4 14.500 180.000 2.000 ca-cc-cc-ha 4 16.000 180.000 2.000 ca-cc-cc-cd 4 16.000 180.000 2.000 ca-cc-nd-na 2 9.500 180.000 2.000 cc-cc-cd-h4 4 16.000 180.000 2.000 cc-cc-cd-na 4 16.000 180.000 2.000 cd-cc-cc-nd 4 16.000 180.000 2.000 | | |  |  |  |

**Table S8.** Missing atom types, point charges, and AMBER force field parameters assigned to **D1**using GAFF2 and structures optimized AM1 level with bond charge corrections (AM1-BCC).

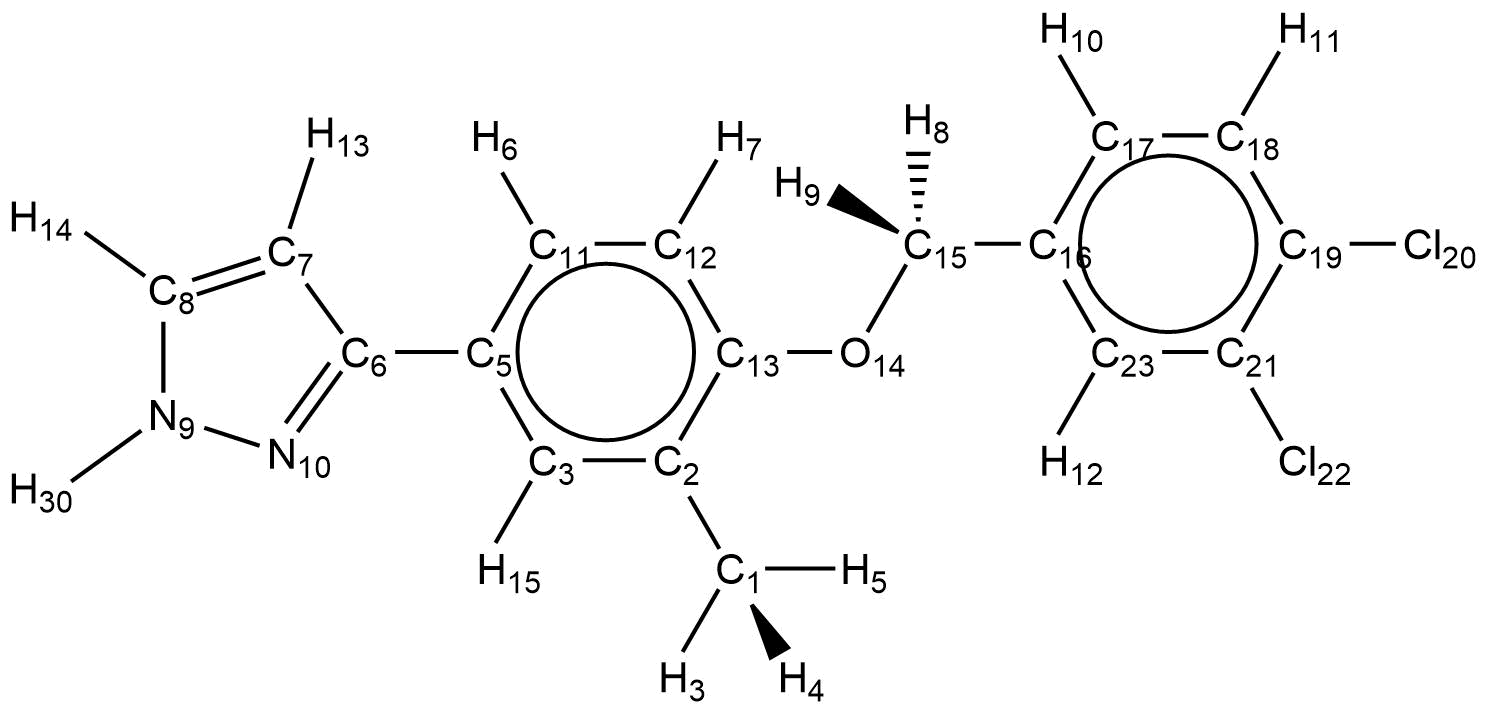

| **D1** | | | | | |
| --- | --- | --- | --- | --- | --- |
| **Atom Name** | **Atom Type** | **Charge** | **Atom Name** | **Atom Type** | **Charge** |
| Cl22 | cl | -0.0694 | H6 | ha | 0.149 |
| C21 | ca | 0.0164 | H7 | ha | 0.143 |
| C23 | ca | -0.115 | C2 | ca | -0.0993 |
| H12 | ha | 0.15 | C1 | c3 | -0.0418 |
| C19 | ca | 0.0214 | H4 | hc | 0.0457 |
| Cl20 | cl | -0.0674 | H5 | hc | 0.0457 |
| C18 | ca | -0.118 | H3 | hc | 0.0457 |
| H11 | ha | 0.154 | C3 | ca | -0.093 |
| C17 | ca | -0.097 | H15 | ha | 0.137 |
| H10 | ha | 0.157 | C5 | ca | -0.1355 |
| C16 | ca | -0.1143 | C6 | cc | 0.3598 |
| C15 | c3 | 0.1717 | C7 | cc | -0.2843 |
| H9 | h1 | 0.0697 | H13 | ha | 0.166 |
| H8 | h1 | 0.0697 | C8 | cd | -0.1513 |
| O14 | os | -0.3269 | H14 | h4 | 0.176 |
| C13 | ca | 0.1151 | N9 | na | -0.0176 |
| C12 | ca | -0.195 | H30 | hn | 0.3207 |
| C11 | ca | -0.077 | N10 | nd | -0.5118 |
| **Parameters** | | | | | |
| **BOND** | | | **ANGLE** | | |
| ca-cl | 187.98 | 1.757 | ca-ca-cl | 64.06 | 119.38 |
| ca-ca | 354.25 | 1.399 | ca-ca-ha | 44.9 | 119.88 |
| ca-ha | 360.69 | 1.086 | ca-ca-ca | 63.67 | 120.02 |
| c3-ca | 243.91 | 1.515 | c3-ca-ca | 60.74 | 120.83 |
| c3-h1 | 344.22 | 1.097 | ca-c3-h1 | 43.81 | 109.6 |
| c3-os | 282.27 | 1.427 | ca-c3-os | 77.6 | 108.88 |
| ca-os | 343.34 | 1.369 | c3-os-ca | 88.26 | 118.39 |
| c3-hc | 345.25 | 1.096 | h1-c3-h1 | 35.64 | 108.55 |
| ca-cc | 296.5 | 1.453 | h1-c3-os | 56.2 | 110.34 |
| cc-cc | 322.02 | 1.427 | ca-ca-os | 79 | 119.11 |
| cc-nd | 419.72 | 1.316 | ca-ca-cc | 62.19 | 120.91 |
| cc-ha | 364.12 | 1.084 | ca-c3-hc | 43.61 | 110.63 |
| cc-cd | 386.24 | 1.373 | hc-c3-hc | 35.8 | 107.73 |
| cd-h4 | 367.29 | 1.082 | ca-cc-cc | 60.65 | 124.73 |
| cd-na | 332.45 | 1.383 | ca-cc-nd | 67.02 | 123.41 |
| hn-na | 481.77 | 1.01 | cc-cc-ha | 44.2 | 119.71 |
| na-nd | 290.41 | 1.357 | cc-cc-cd | 63.77 | 119.19 |
|  |  |  | cc-nd-na | 93.94 | 103.84 |
|  |  |  | cc-cc-nd | 67.03 | 126.05 |
|  |  |  | cc-cd-h4 | 44.2 | 127.64 |
|  |  |  | cc-cd-na | 71.99 | 108.99 |
|  |  |  | cd-cc-ha | 45.3 | 121.4 |
|  |  |  | cd-na-hn | 55.43 | 125.24 |
|  |  |  | cd-na-nd | 88.18 | 112.21 |
|  |  |  | h4-cd-na | 48.08 | 120.17 |
|  |  |  | hn-na-nd | 61.31 | 119.24 |
| **IMPROPER** | | | **DIHEDRAL** | | |
| ca-ca-ca-cl 1.1 180.0 2.0 ca-ca-ca-ha 1.1 180.0 2.0  c3-ca-ca-ca 1.1 180.0 2.0 ca-ca-ca-os 1.1 180.0 2.0  ca-ca-ca-cc 1.1 180.0 2.0  ca-cc-cc-nd 1.1 180.0 2.0  cc-cd-cc-ha 1.1 180.0 2.0  cc-h4-cd-na 1.1 180.0 2.0  cd-hn-na-nd 1.1 180.0 2.0 | | | ca-ca-cc-nd 4 4.000 180.000 -2.000  ca-ca-cc-nd 1 0.750 180.000 -3.000  ca-ca-cc-nd 1 0.820 180.000 1.  cc-ca-ca-ha 4 14.500 180.000 2.000 ca-cc-cc-ha 4 16.000 180.000 2.000 ca-cc-cc-cd 4 16.000 180.000 2.000 ca-cc-nd-na 2 9.500 180.000 2.000 cc-cc-cd-h4 4 16.000 180.000 2.000 cc-cc-cd-na 4 16.000 180.000 2.000  cc-cd-na-hn 4 6.800 180.000 2.000 cc-cd-na-nd 4 6.800 180.000 2.000 ha-cc-cd-h4 4 16.000 180.000 2.000 ha-cc-cd-na 4 16.000 180.000 2.000 cd-na-nd-cc 2 9.600 180.000 2.000 h4-cd-na-hn 4 6.800 180.000 2.000 h4-cd-na-nd 4 6.800 180.000 2.000 hn-na-nd-cc 2 9.600 180.000 2.000 ha-cc-cc-nd 4 16.000 180.000 2.000 cd-cc-cc-nd 4 16.000 180.000 2.000  cc-cc-nd-na 2 9.500 180.000 2.000  hc-c3-ca-ca 1 0.000 0.000 1.000 ca-ca-ca-cc 4 14.500 180.000 2.000 ca-ca-cc-cc 1 3.770 180.000 2.000  ha-ca-ca-os 4 14.500 180.000 2.000 | | |
| **DIHEDRAL** | | |  |  |  |
| cl-ca-ca-ha 4 14.500 180.000 2.000 ca-ca-ca-cl 4 14.500 180.000 2.000 cl-ca-ca-cl 4 14.500 180.000 2.000 ca-ca-ca-ca 4 14.500 180.000 2.000 c3-ca-ca-ca 4 14.500 180.000 2.000 ca-ca-ca-ha 4 14.500 180.000 2.000 h1-c3-ca-ca 6 0.000 0.000 2.000 os-c3-ca-ca 6 0.000 0.000 2.000 c3-ca-ca-ha 4 14.500 180.000 2.000  ha-ca-ca-ha 4 14.500 180.000 2.000 ca-c3-os-ca 3 1.150 0.000 3.000 ca-ca-os-c3 1 1.660 180.000 2.000 h1-c3-os-ca 3 1.150 0.000 3.000 ca-ca-ca-os 4 14.500 180.000 2.000  c3-ca-ca-os 4 14.500 180.000 2.000 | | |  |  |  |

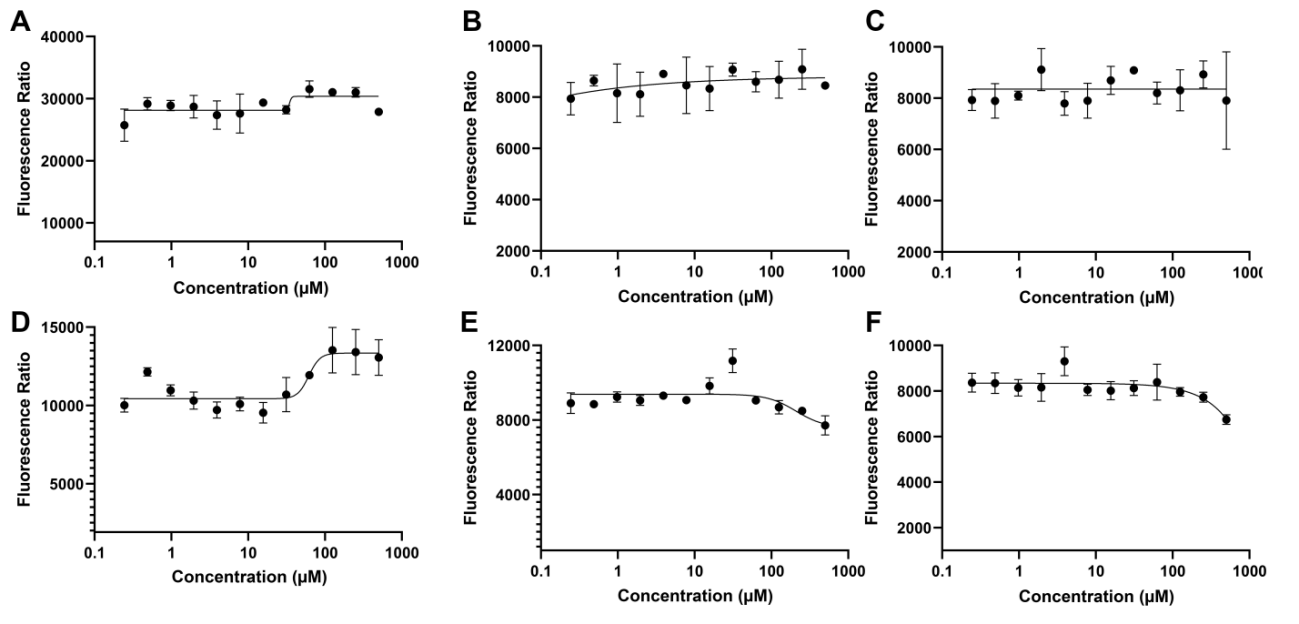

**Figure S1**. Inhibition curve of compound D1 - D6 with ICOS/ICOS-L (10 nM / 10 nM) using FRET assay. Buffer for RRET assay: PBS with 5% DMSO.

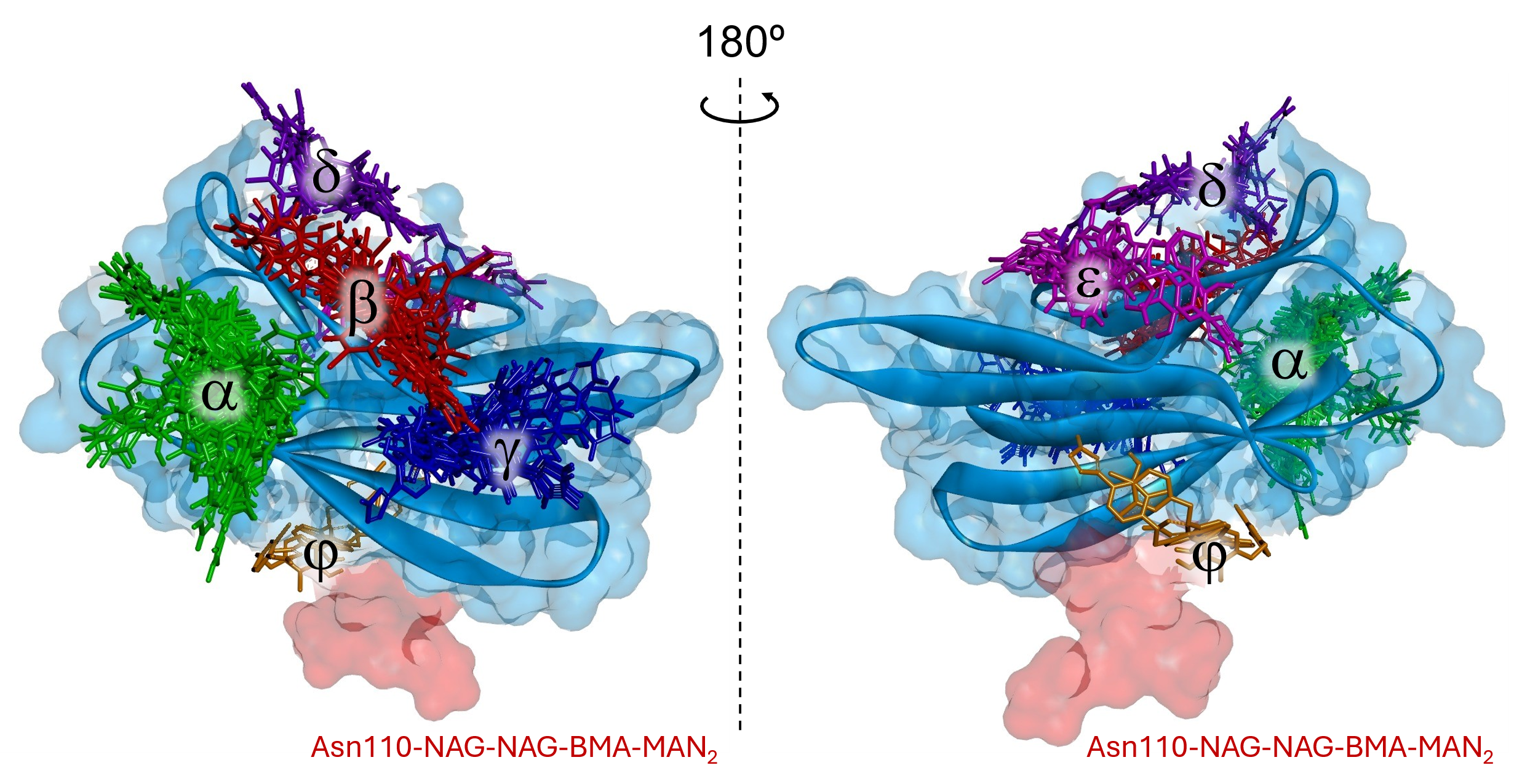

**Figure S2**. ICOS structure with highlighted six (α, β, γ, δ, ε, φ) possible binding regions of RH00458 compound.

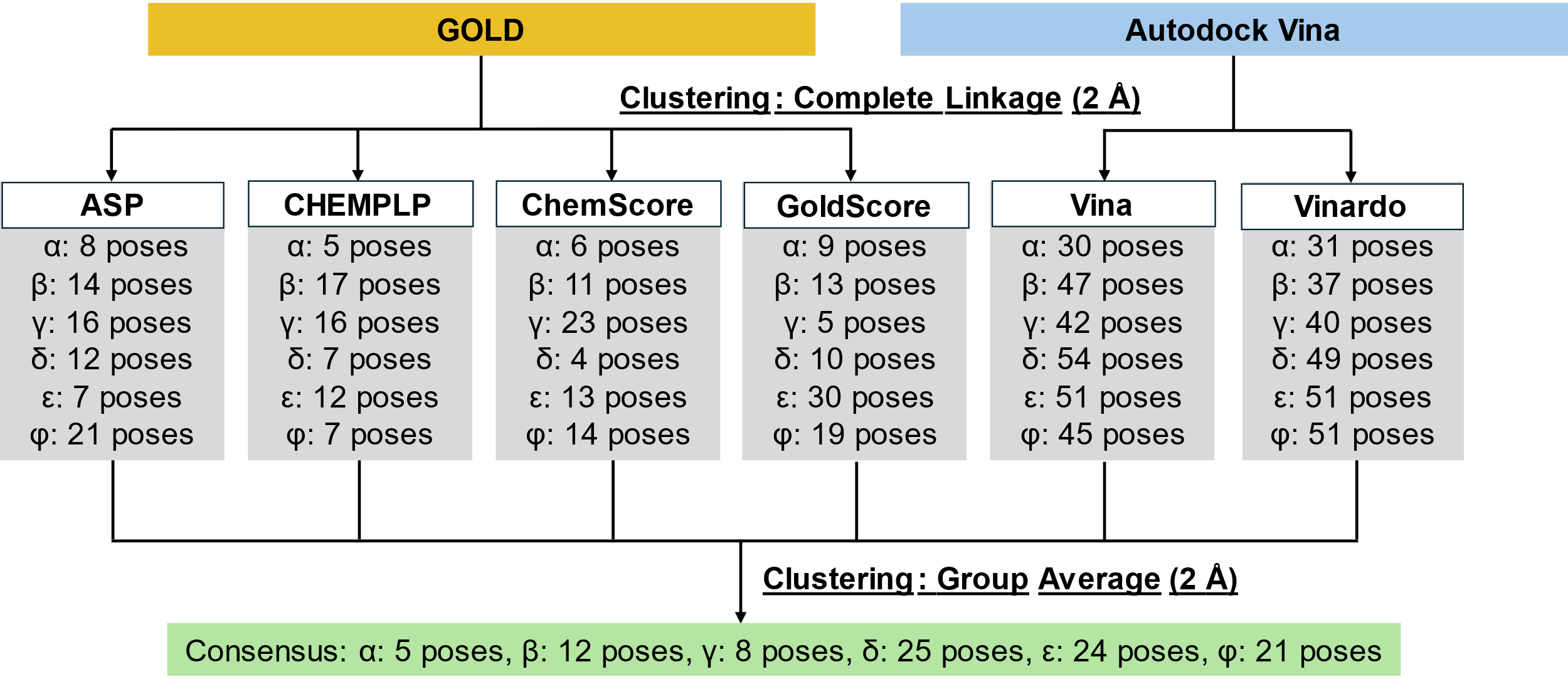

**Figure S3.** Summary of consensus docking approach.

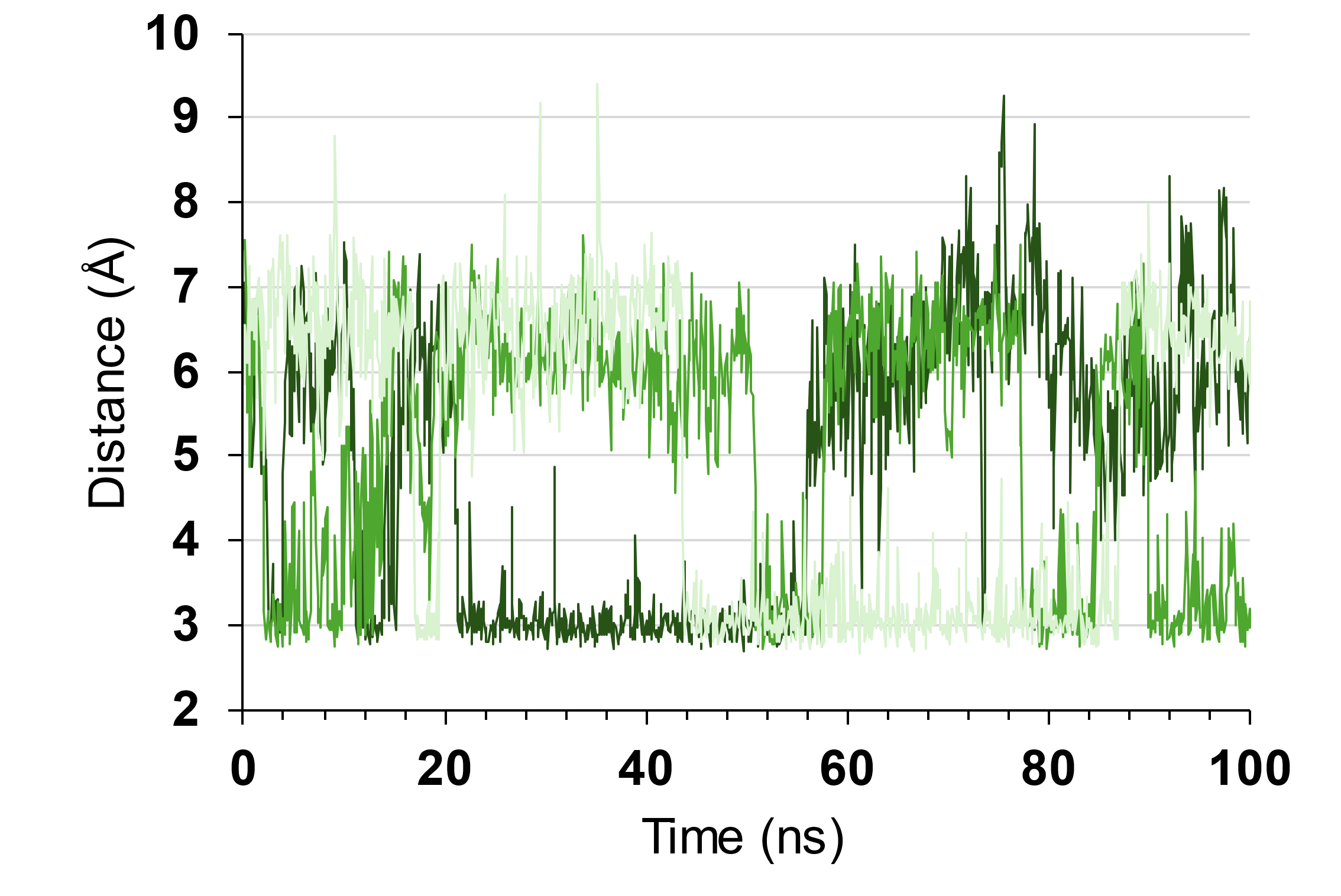

**Figure S4.** Evolution of O1 and Lys52:Nζ distances from 3 replicas of 100 ns MD simulations for **9**.

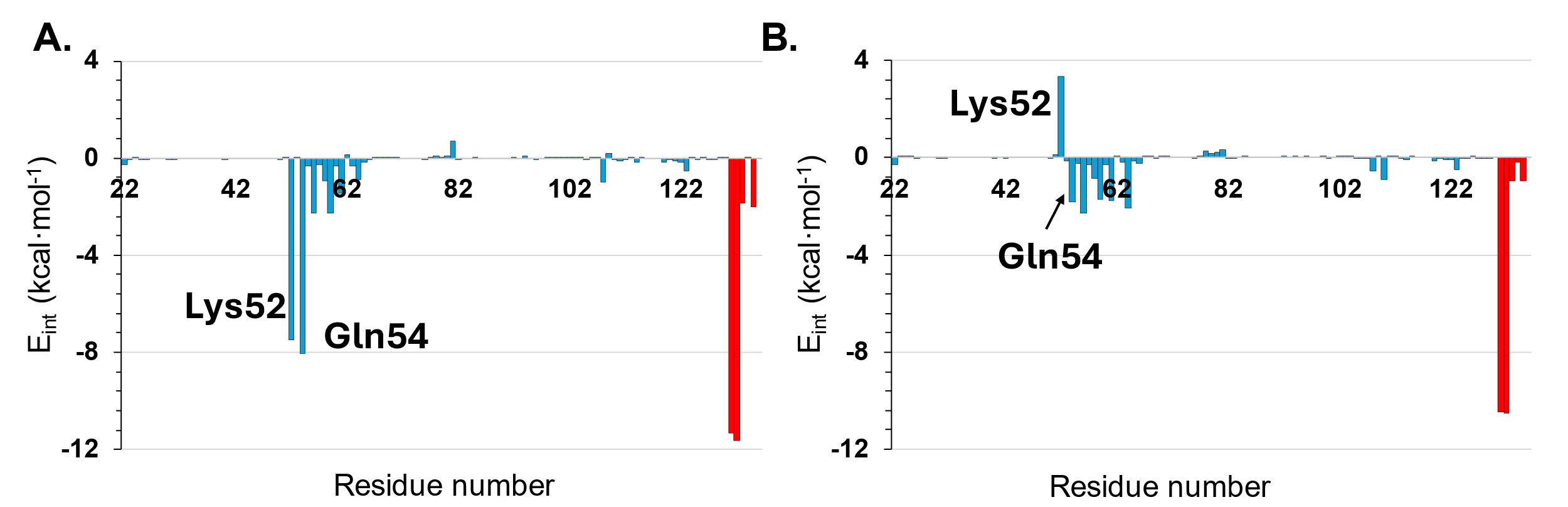

**Figure S5.** Overall interaction energy (E_int_) computed per residue for **9** (**A**) and **D1** (**B**) compounds and glycan (in red) and protein (in blue) residues. In the case of the 9, all 30,000 structures generated in three replicas of MD simulations were included. In the case of the D1 only frames with r.m.s.d. below 5.5 Å were considered.

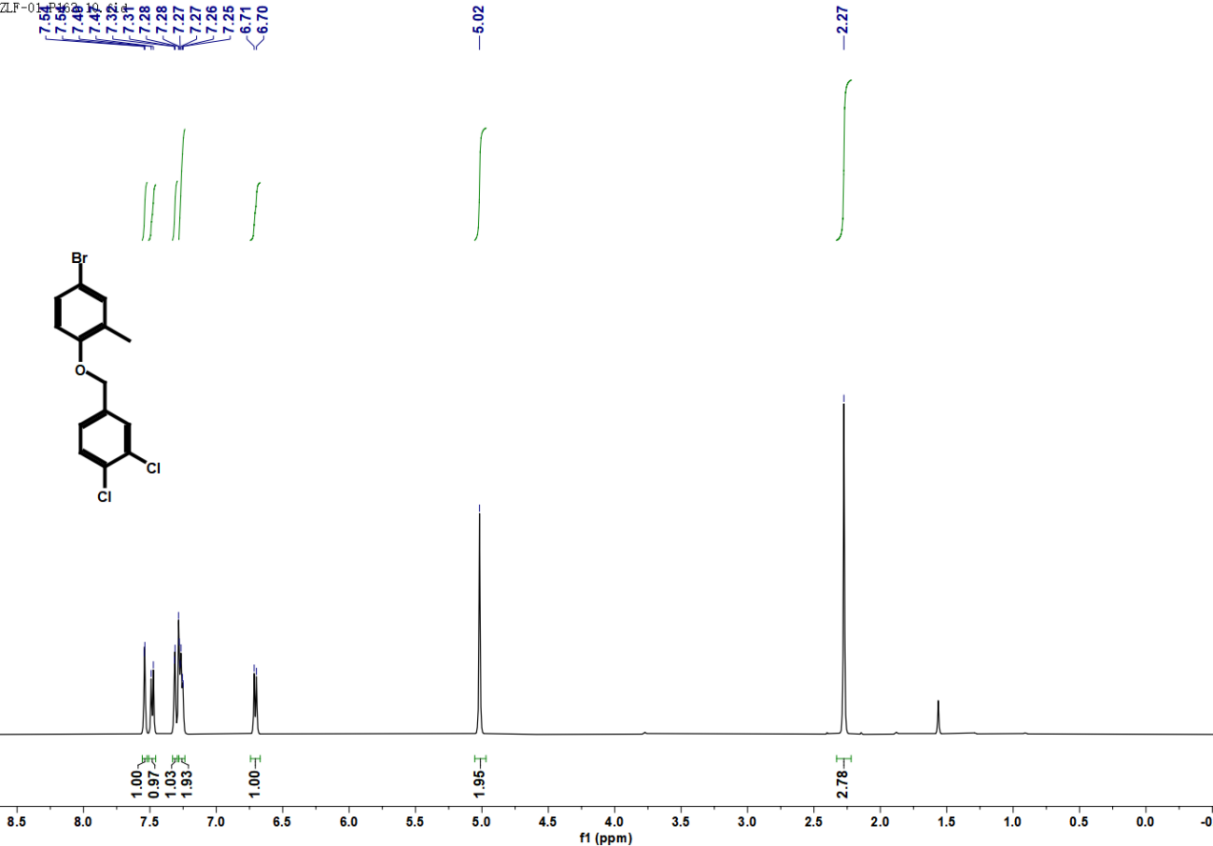

**Figure S6.** ^1^H NMR spectrum of compound **I3**.

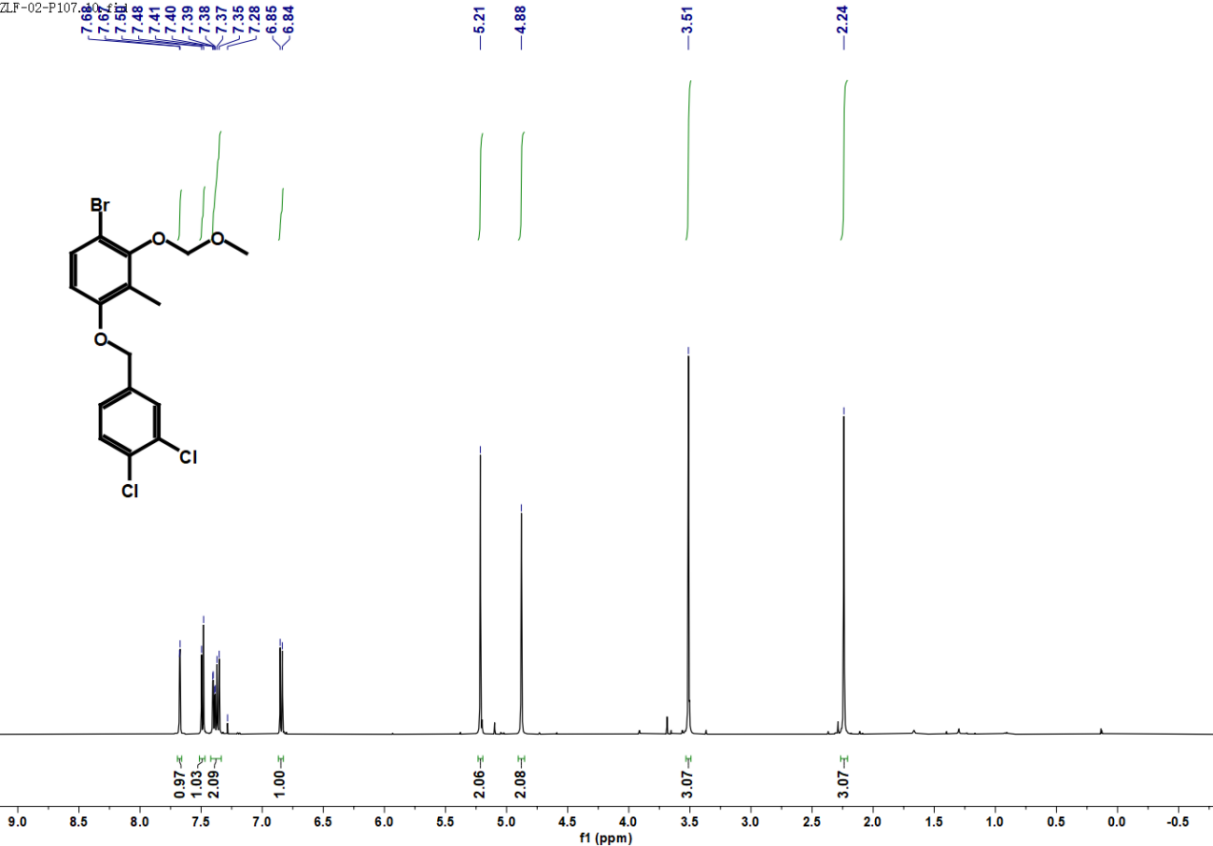

**Figure S7.** ^1^H NMR spectrum of compound **I5**.

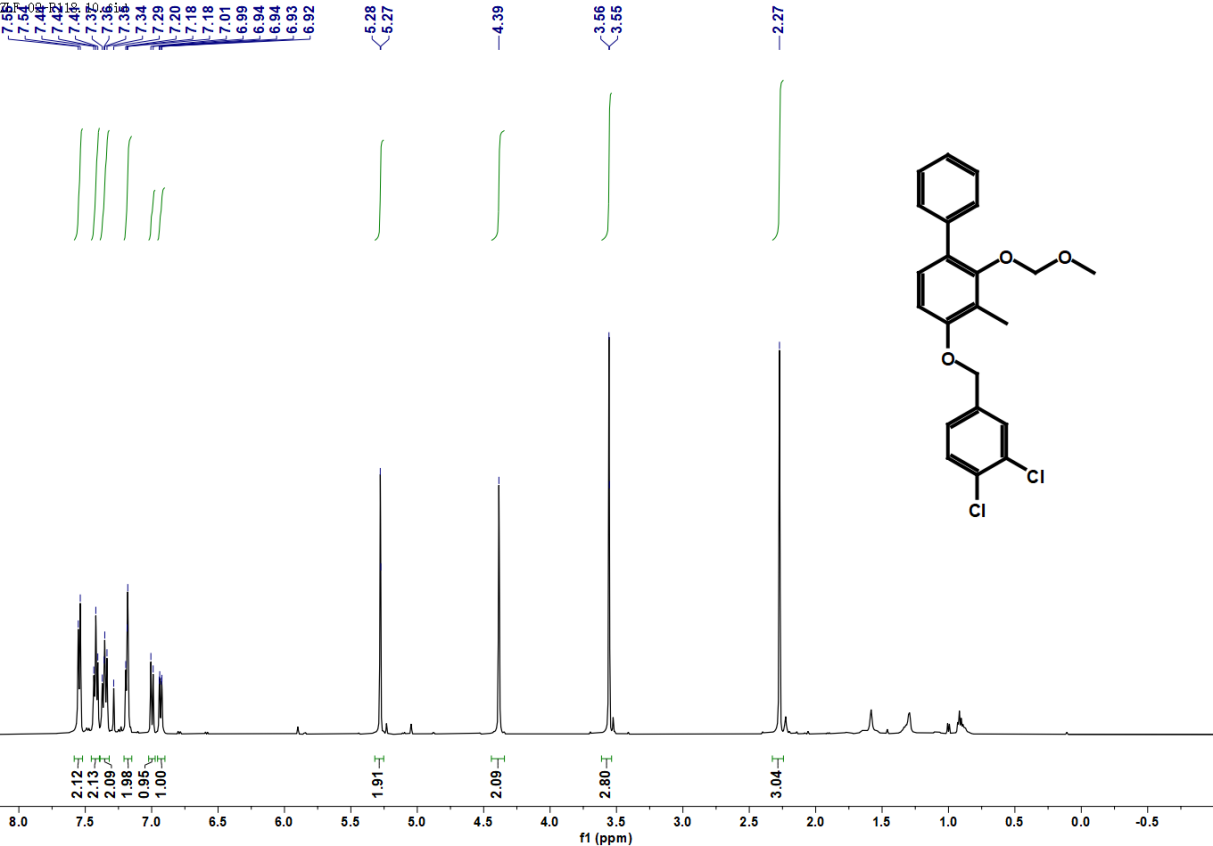

**Figure S8.** ^1^H NMR spectrum of compound **I6**.

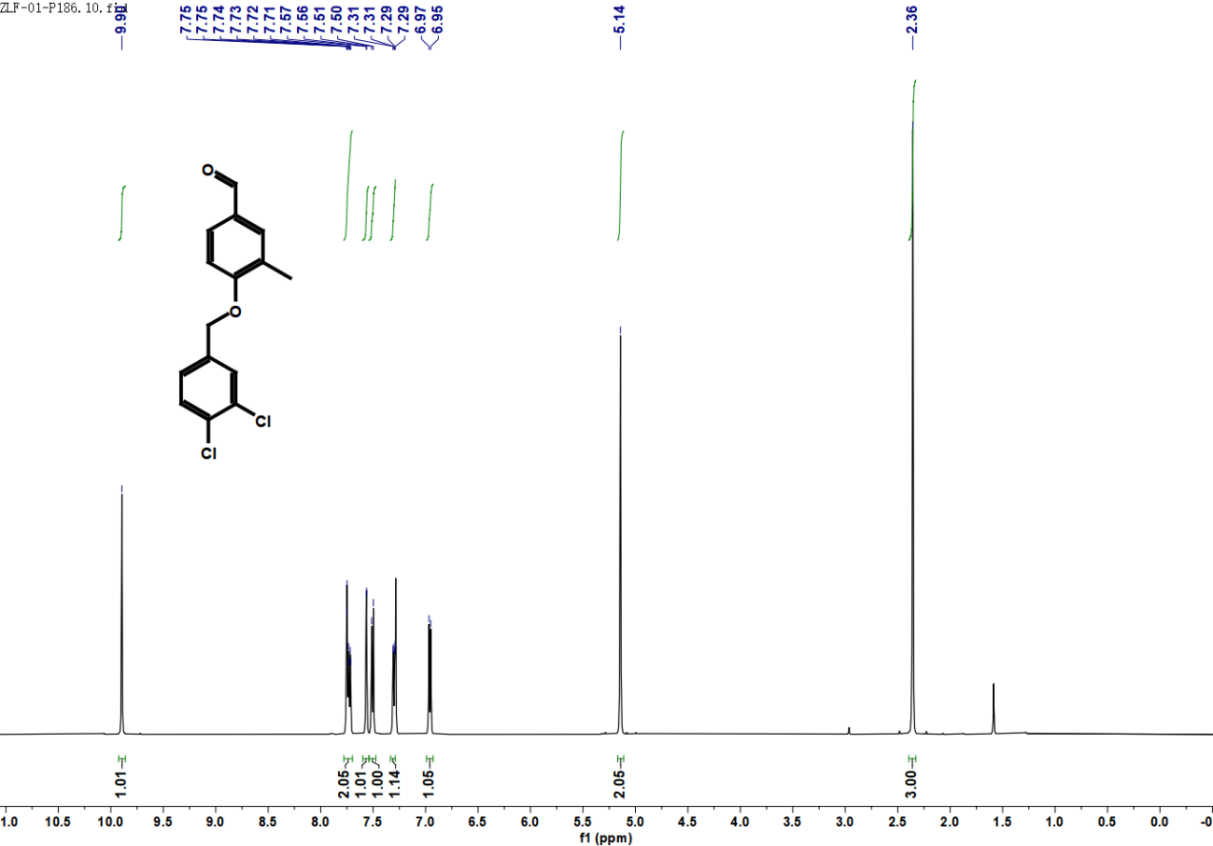

**Figure S9.** ^1^H NMR spectrum of compound **I7**.

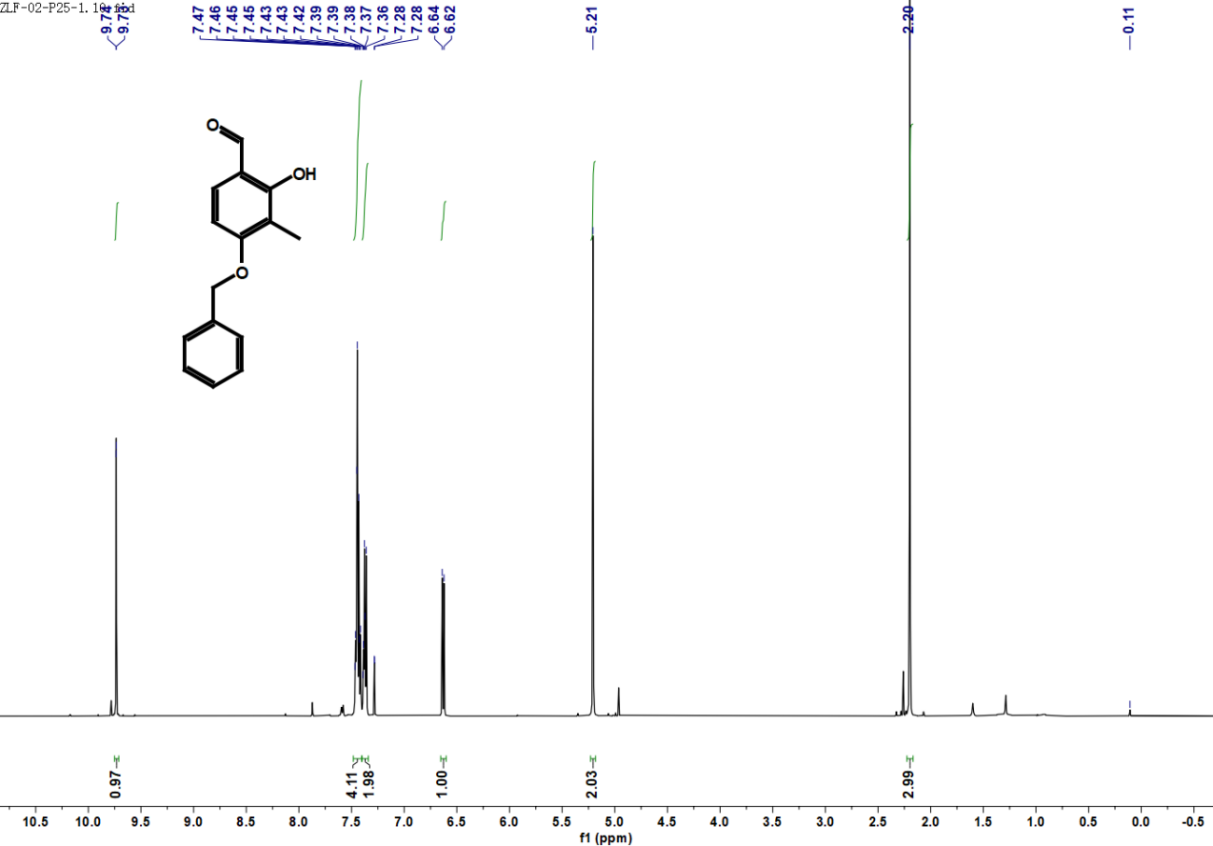

**Figure S10.** ^1^H NMR spectrum of compound **I8**.

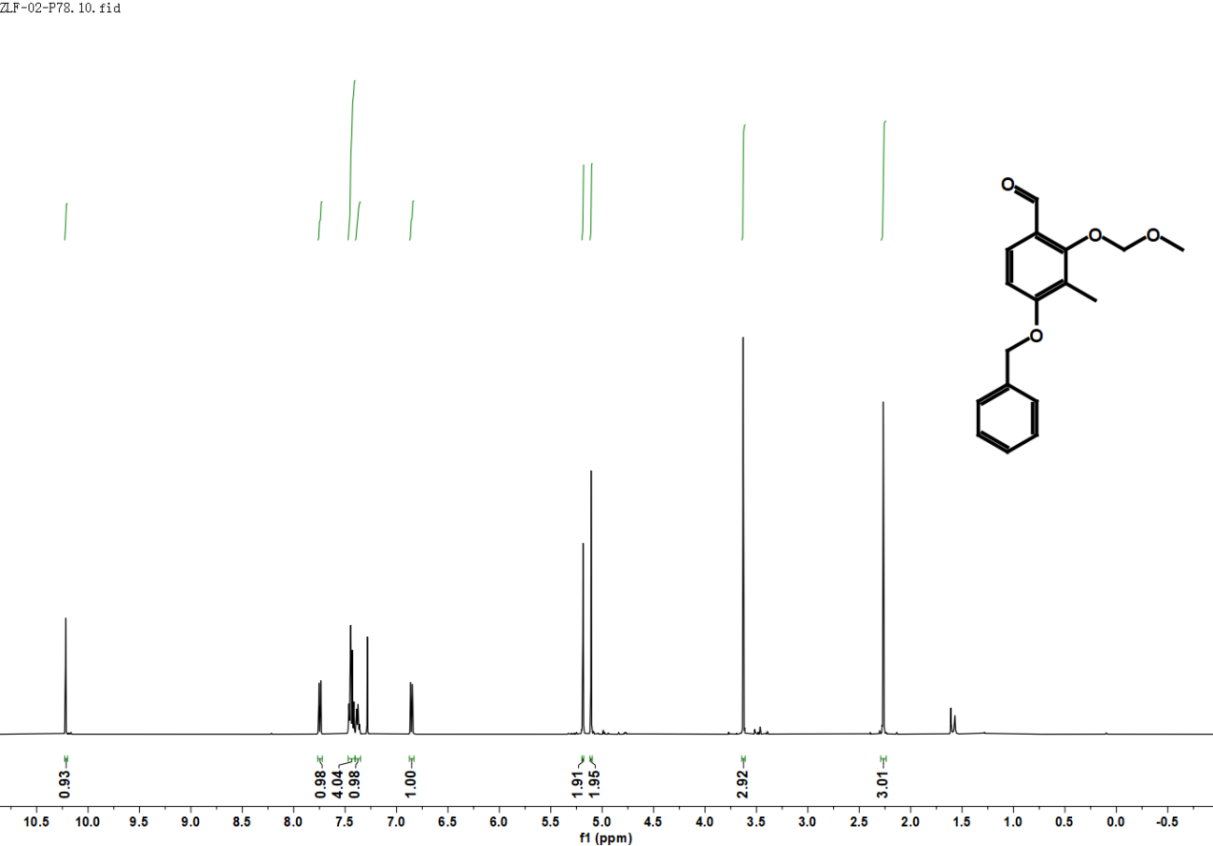

**Figure S11.** ^1^H NMR spectrum of compound **I9**.

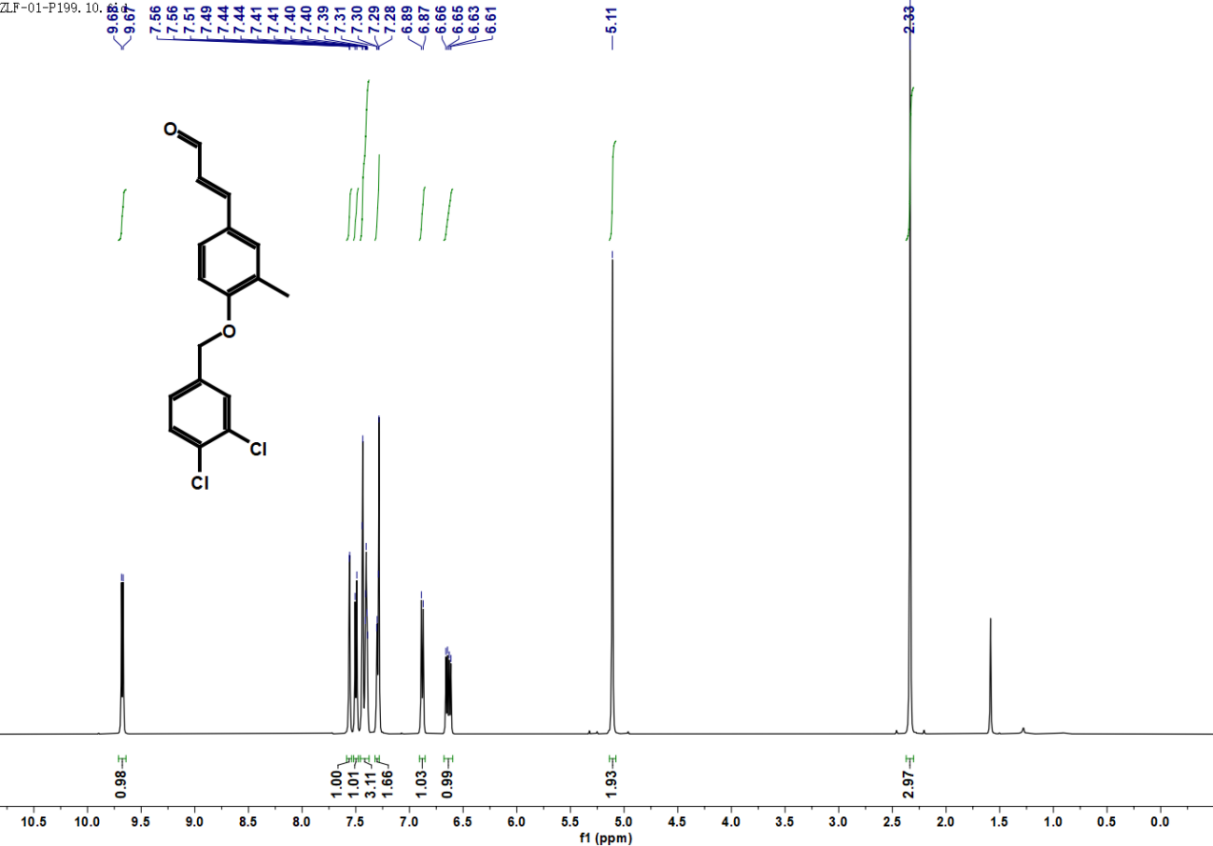

**Figure S12.** ^1^H NMR spectrum of compound **I10**.

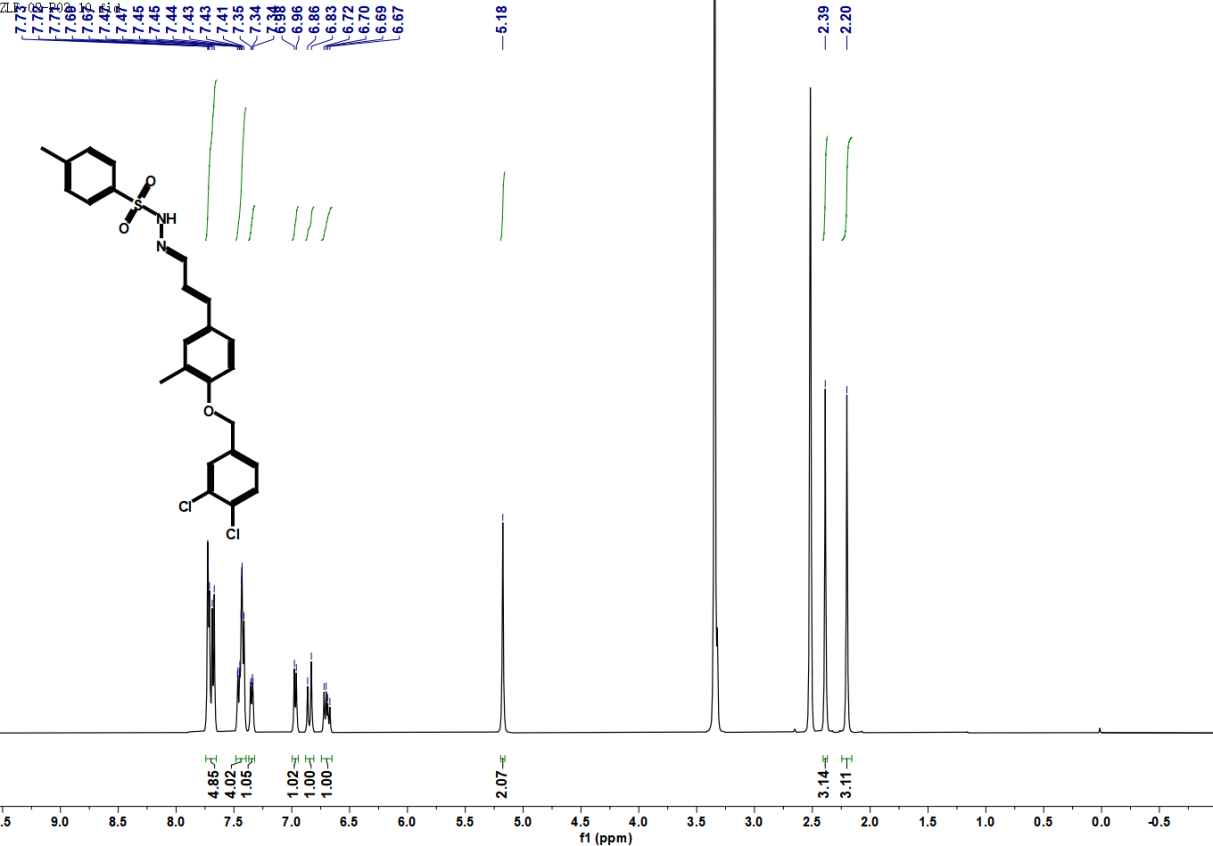

**Figure S13.** ^1^H NMR spectrum of compound **I12**.

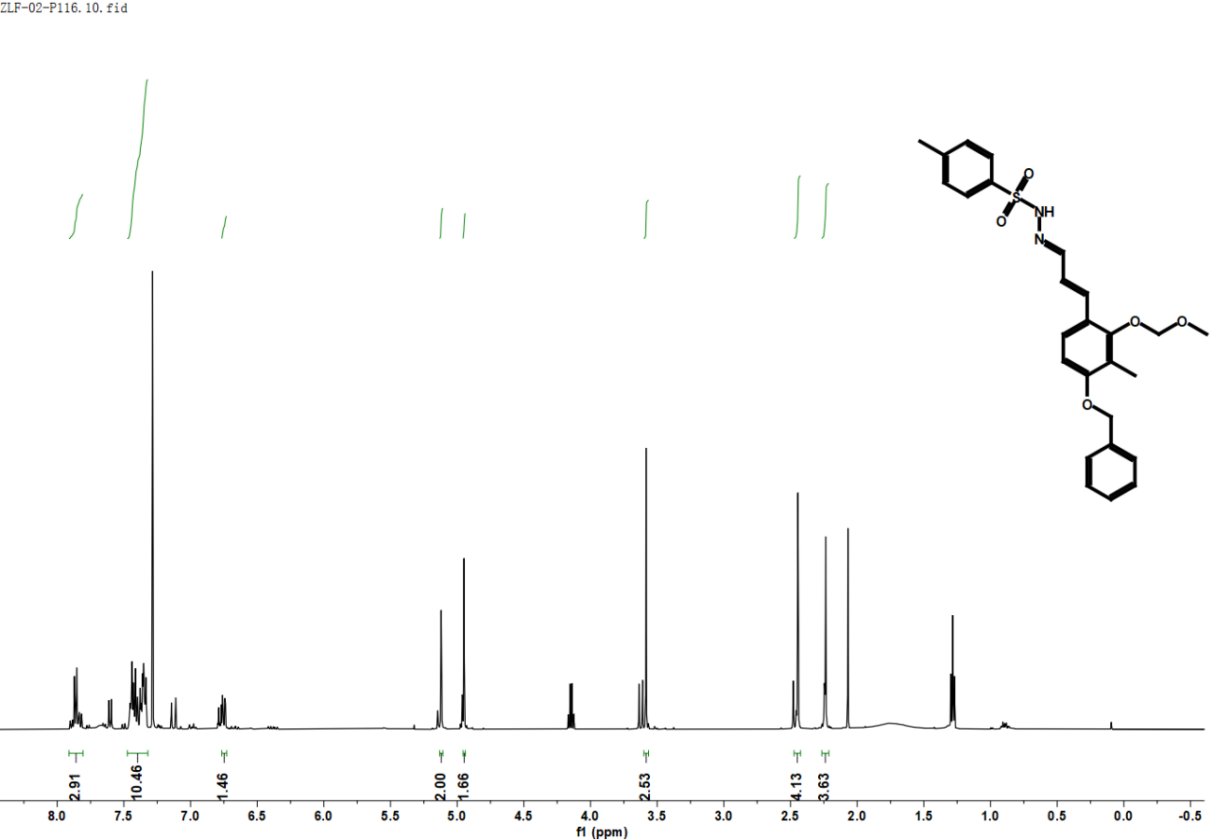

**Figure S14.** ^1^H NMR spectrum of compound **I13**.

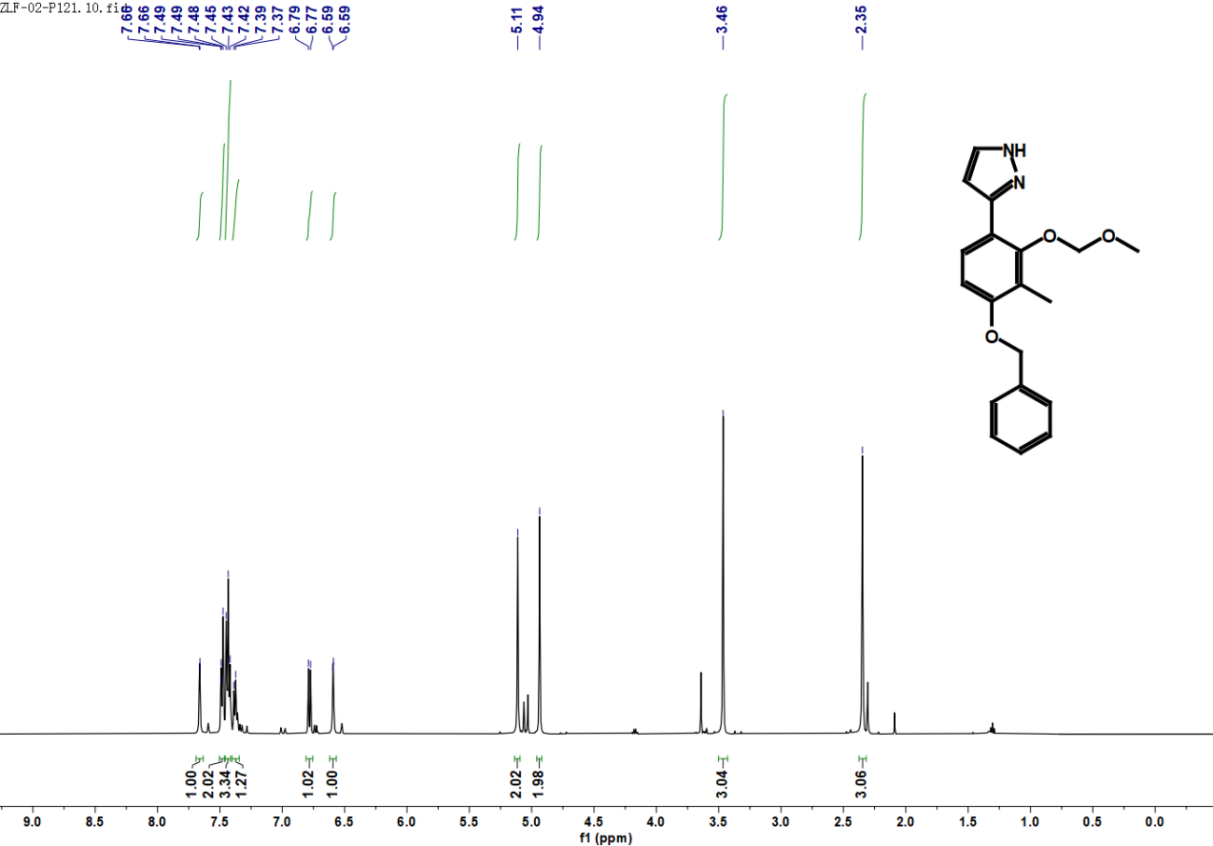

**Figure S15.** ^1^H NMR spectrum of compound **I14**.

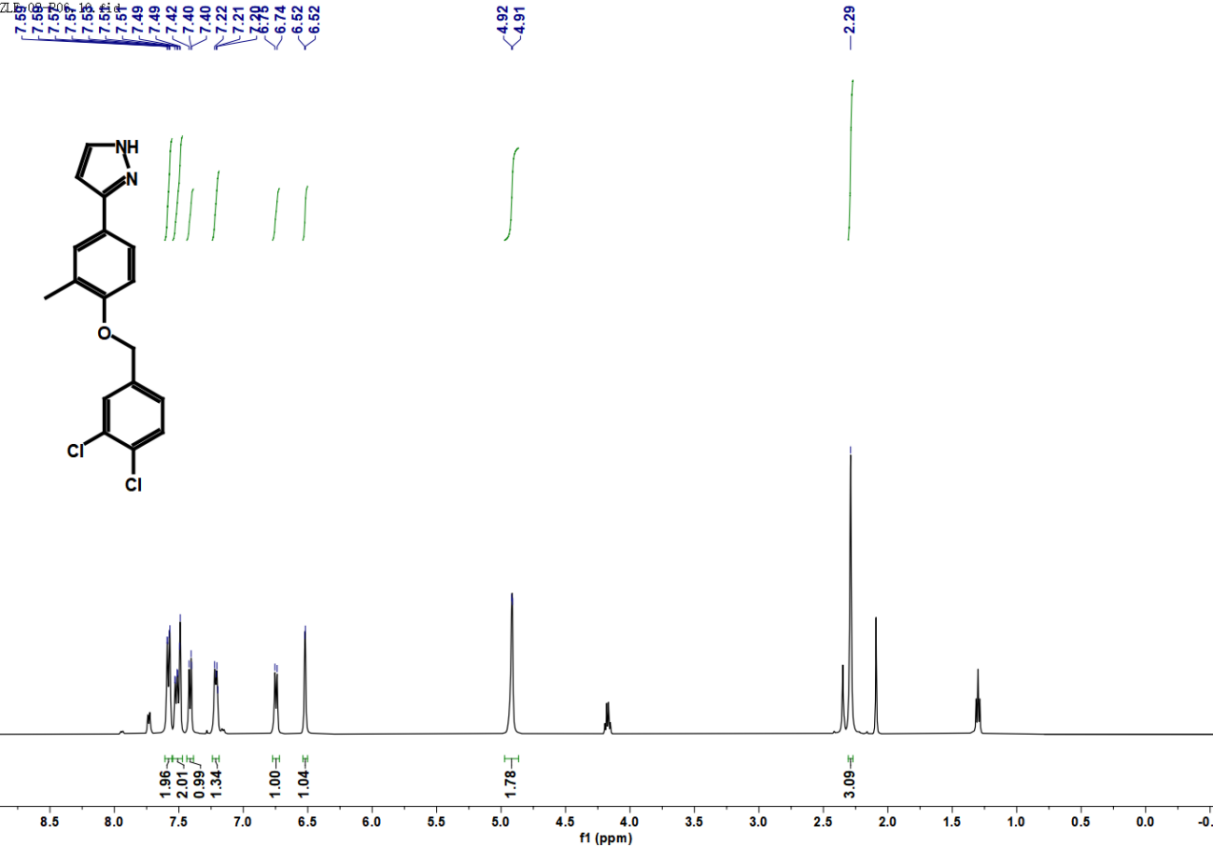

**Figure S16.** ^1^H NMR spectrum of compound **D1**.

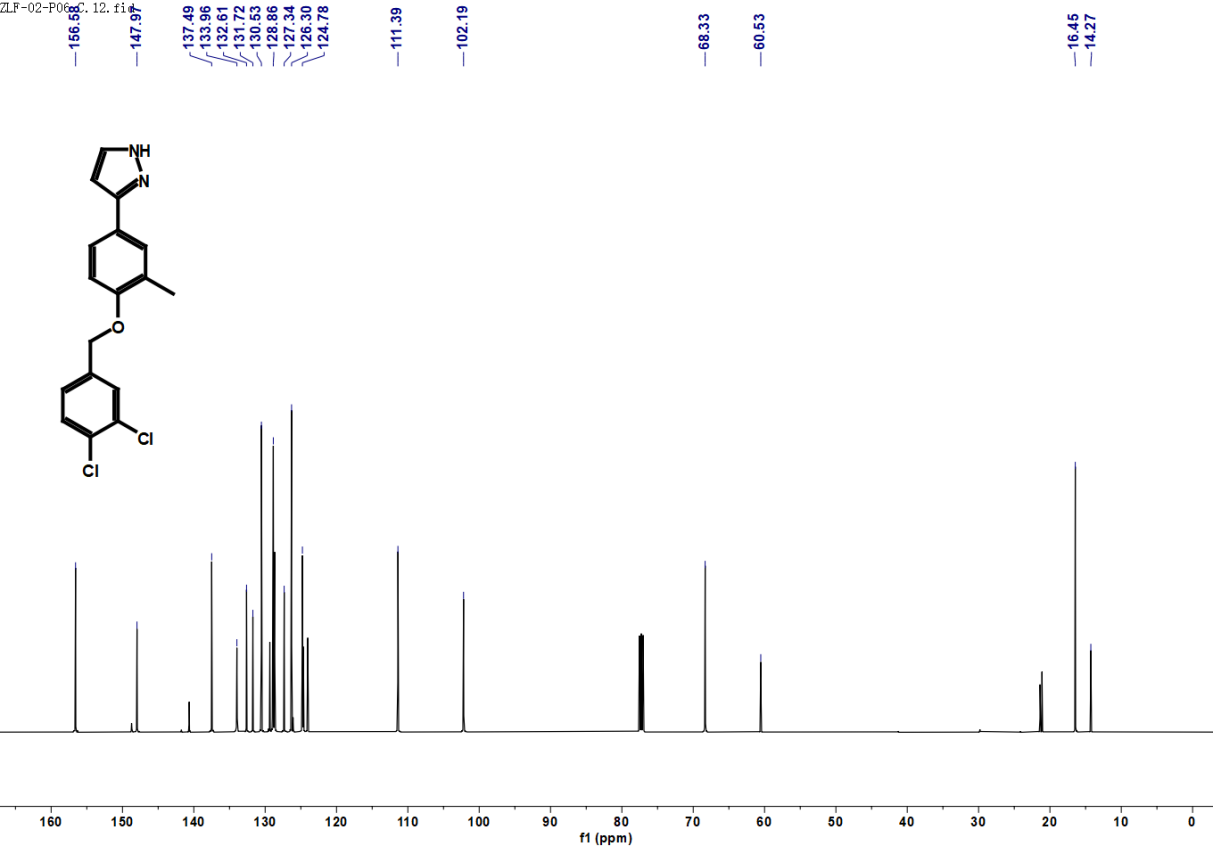

**Figure S17.** ^13^C NMR spectrum of compound **D1**.

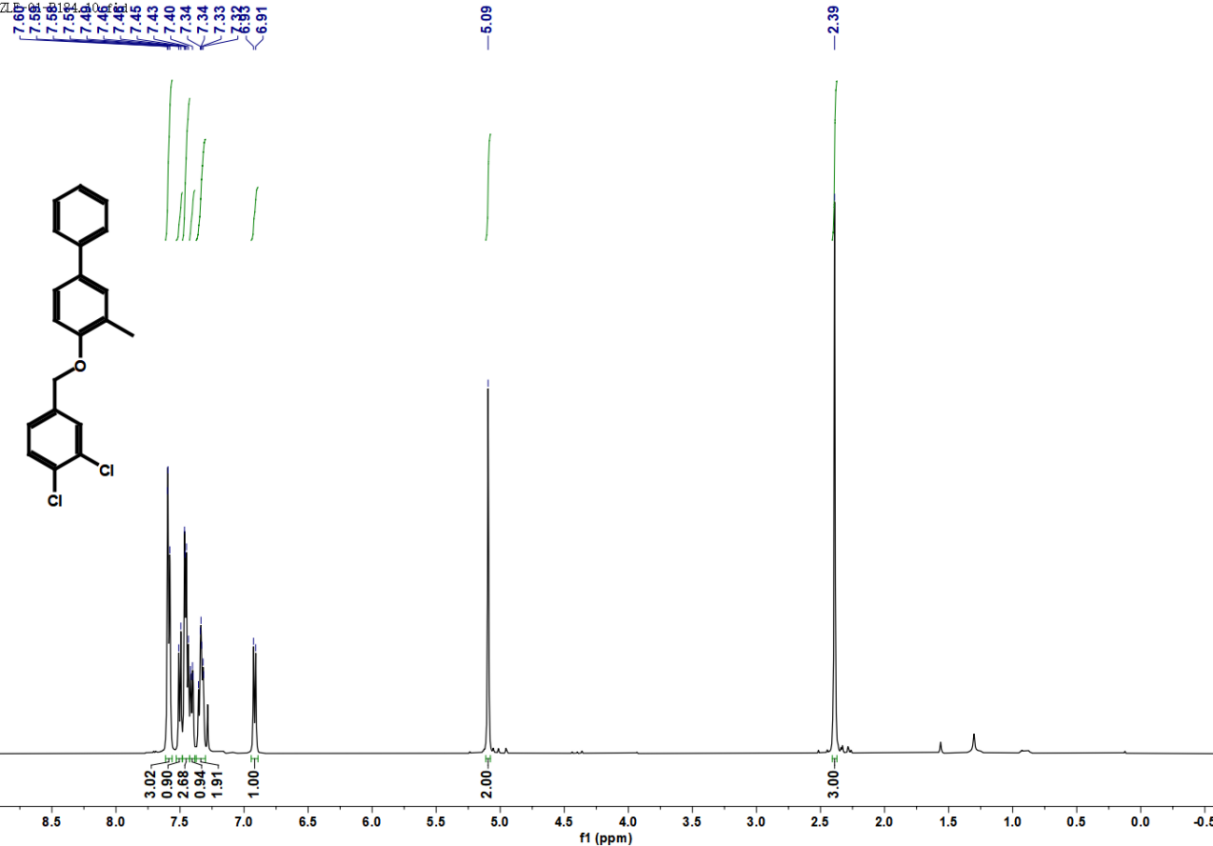

**Figure S18.** ^1^H NMR spectrum of compound **D2**.

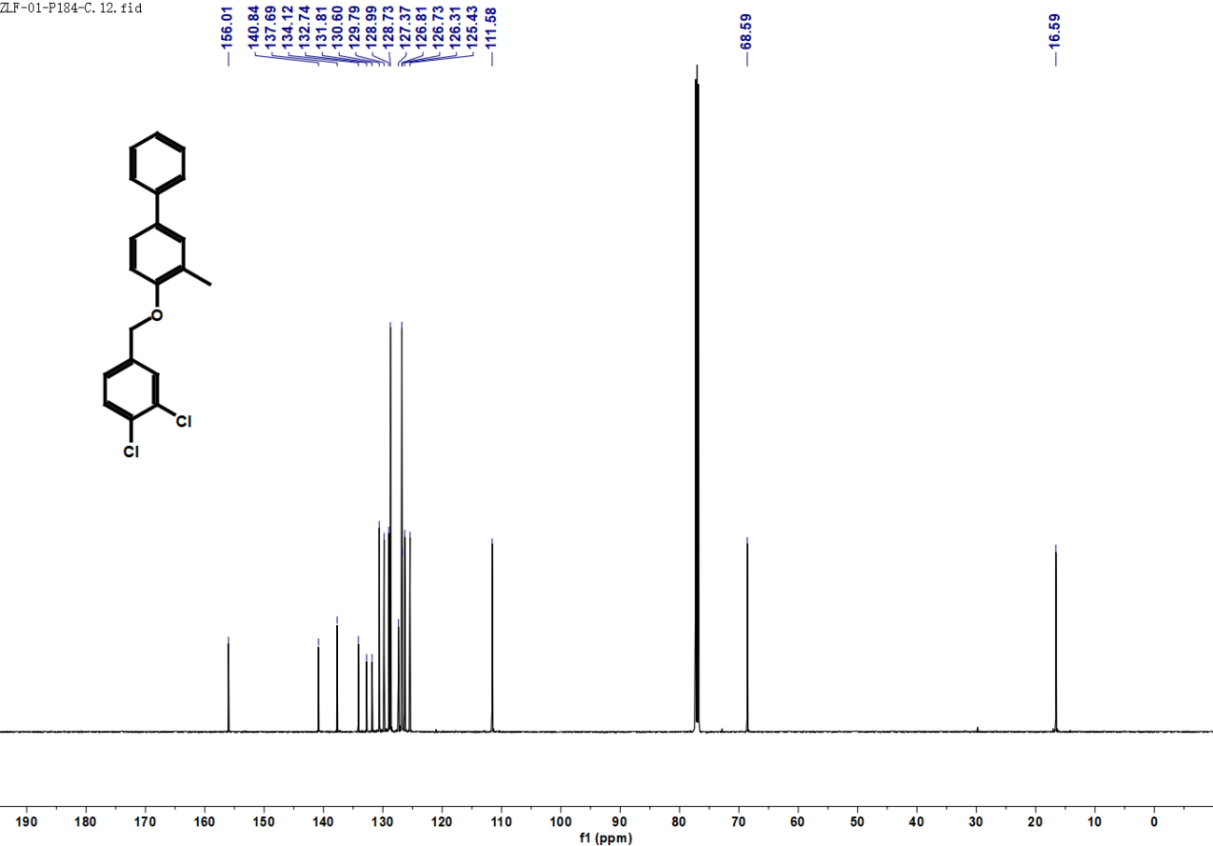

**Figure S19.** ^13^C NMR spectrum of compound **D2**.

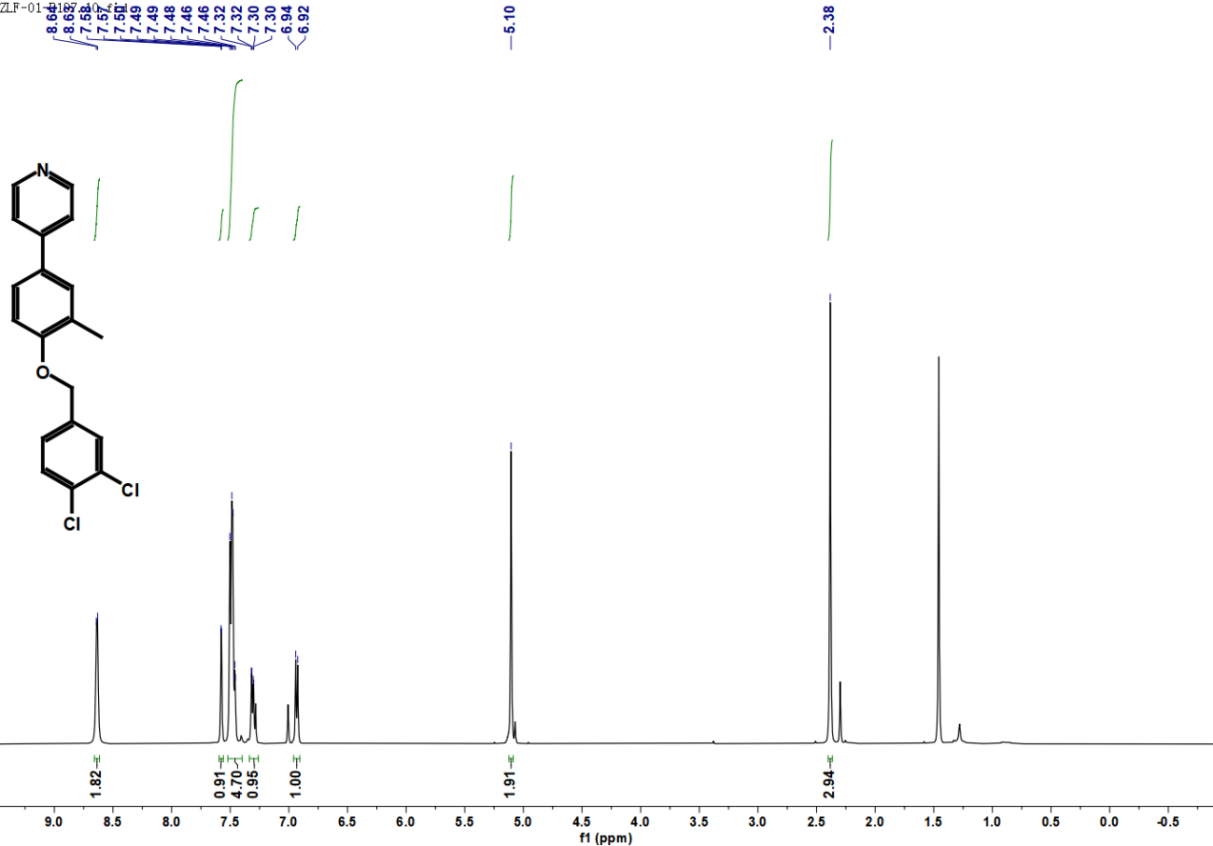

**Figure S20.** ^1^H NMR spectrum of compound **D3**.

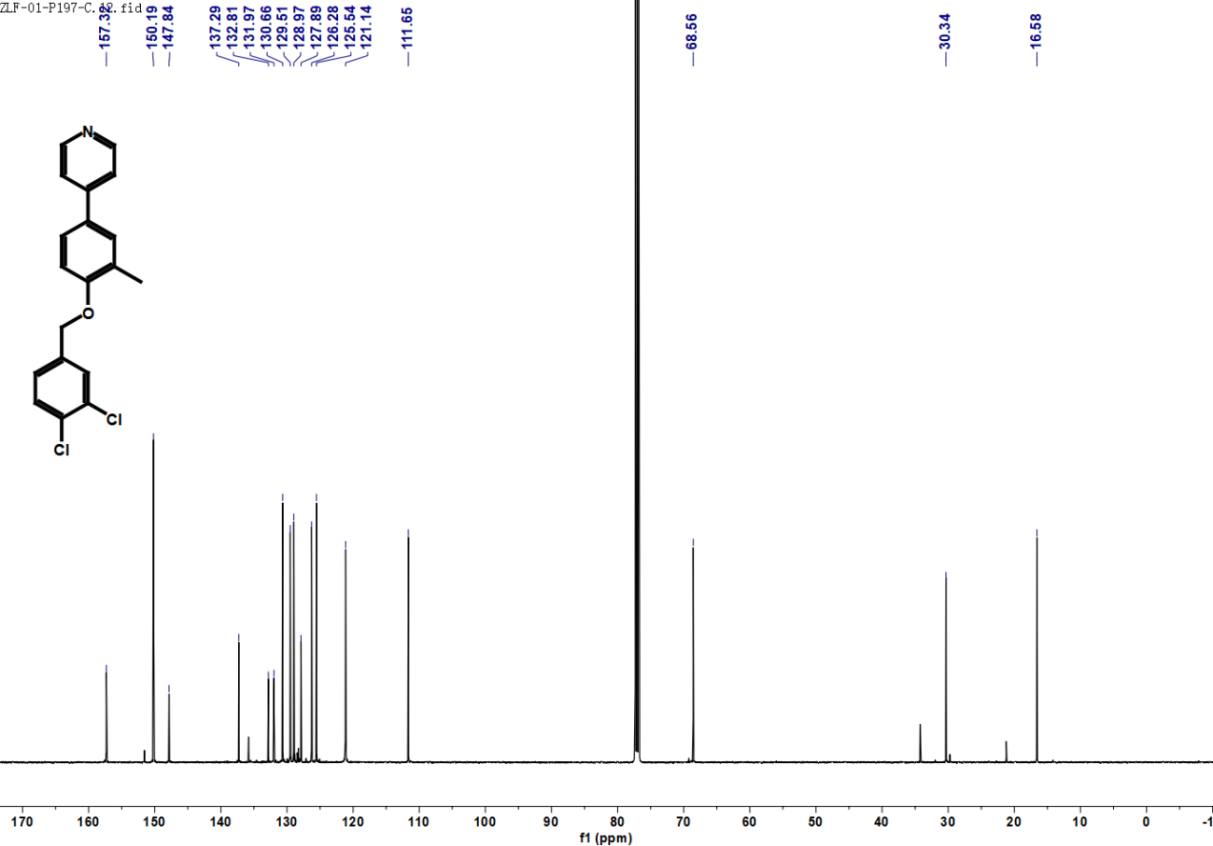

**Figure S21.** ^13^C NMR spectrum of compound **D3**.

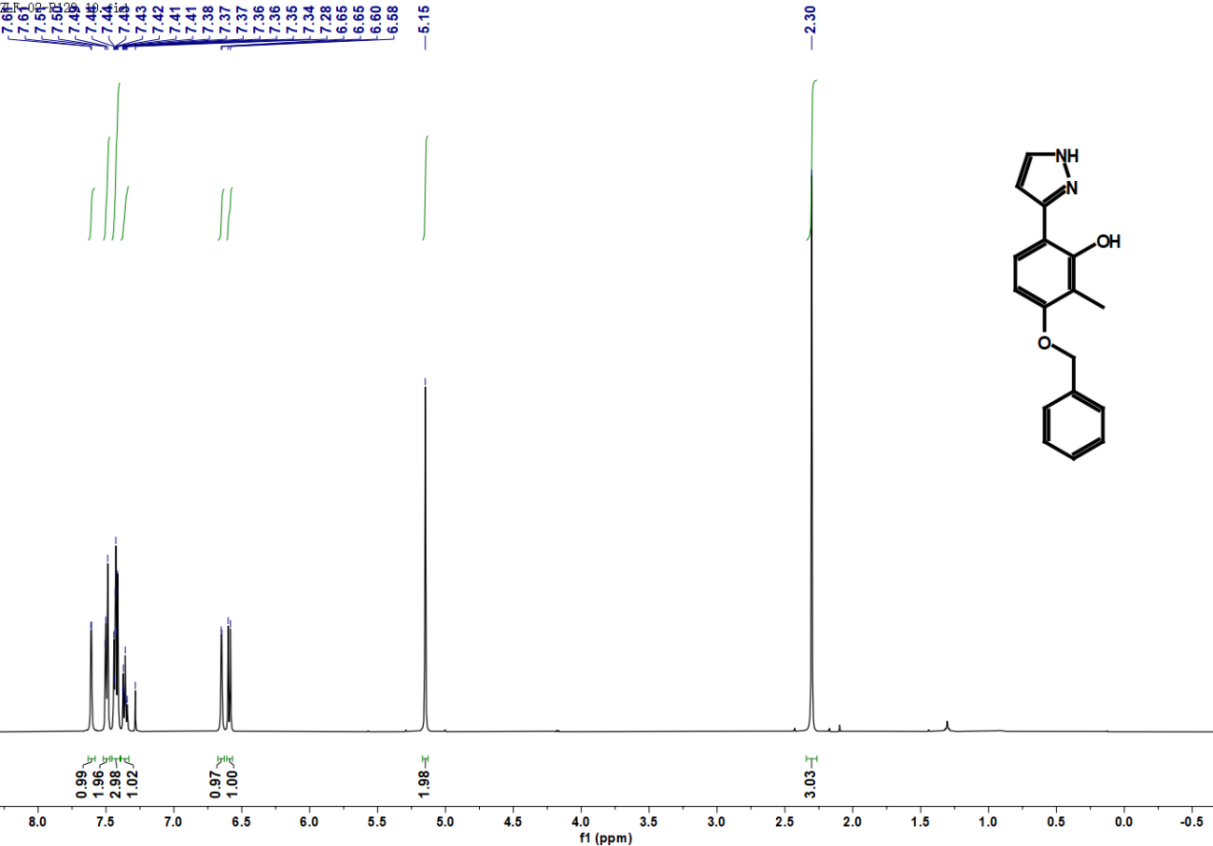

**Figure S22.** ^1^H NMR spectrum of compound **D4**.

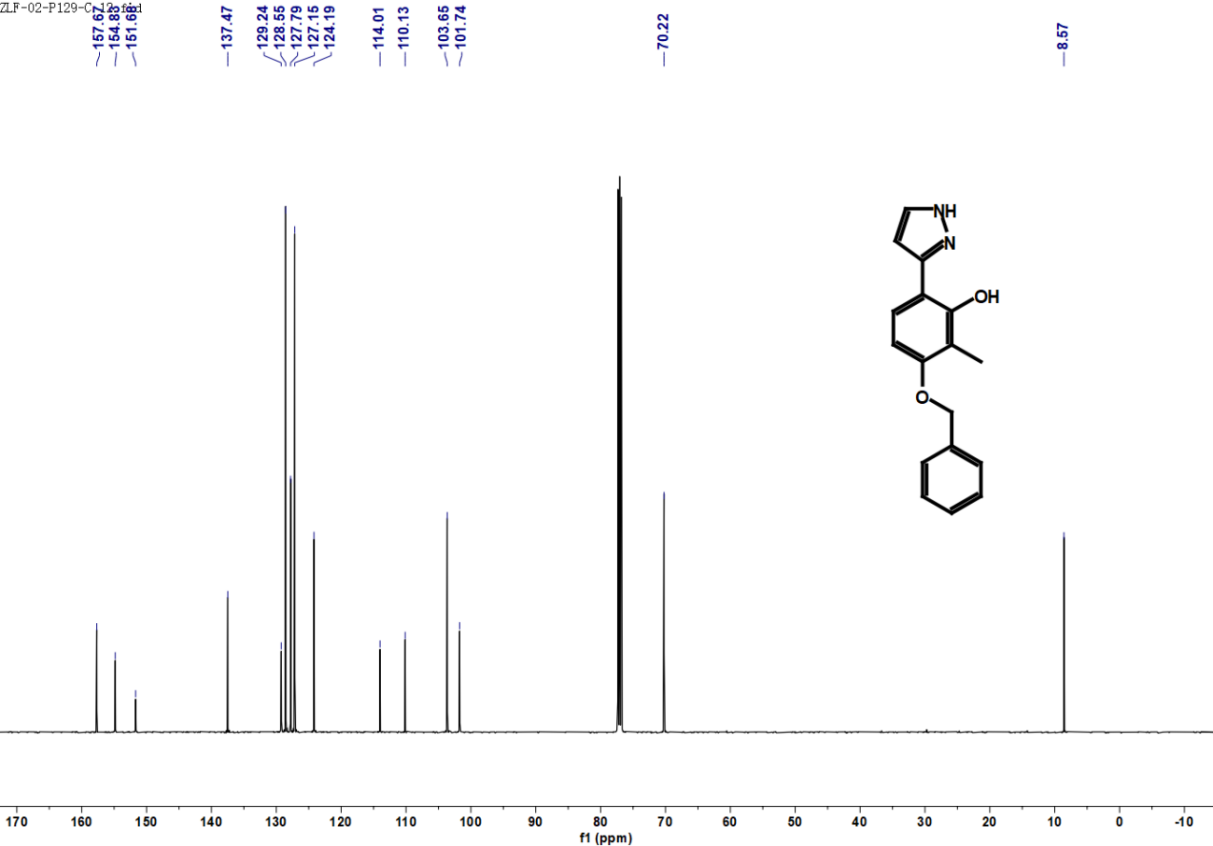

**Figure S23.** ^13^C NMR spectrum of compound **D4**.

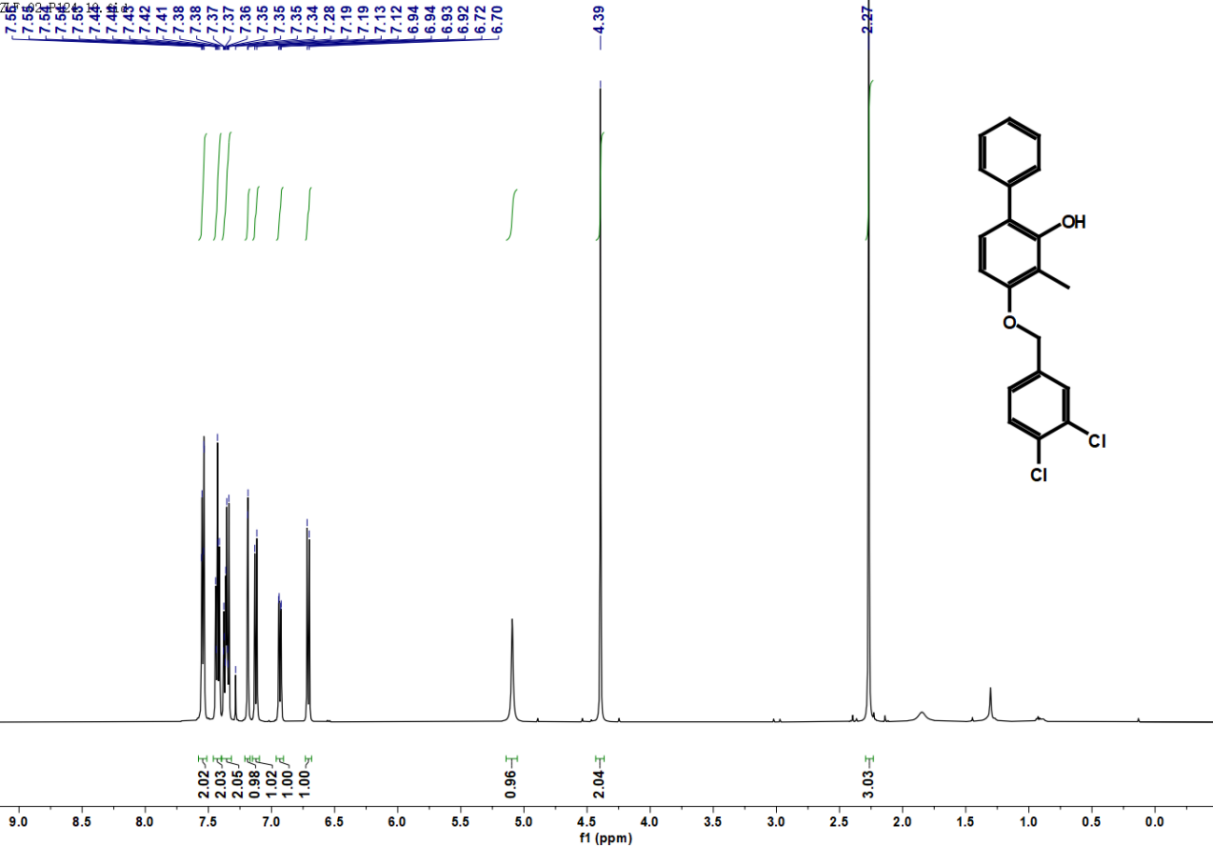

**Figure S24.** ^1^H NMR spectrum of compound **D5**.

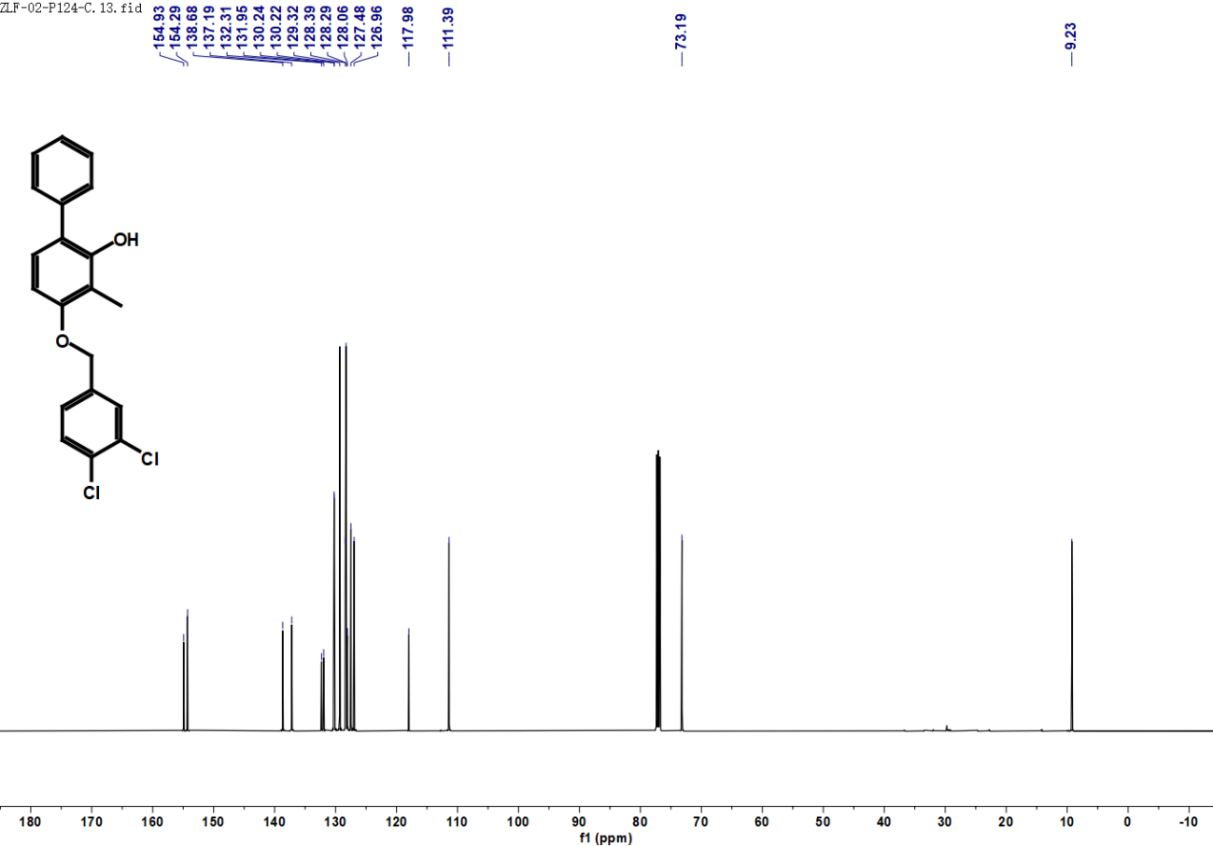

**Figure S25.** ^13^C NMR spectrum of compound **D5**.

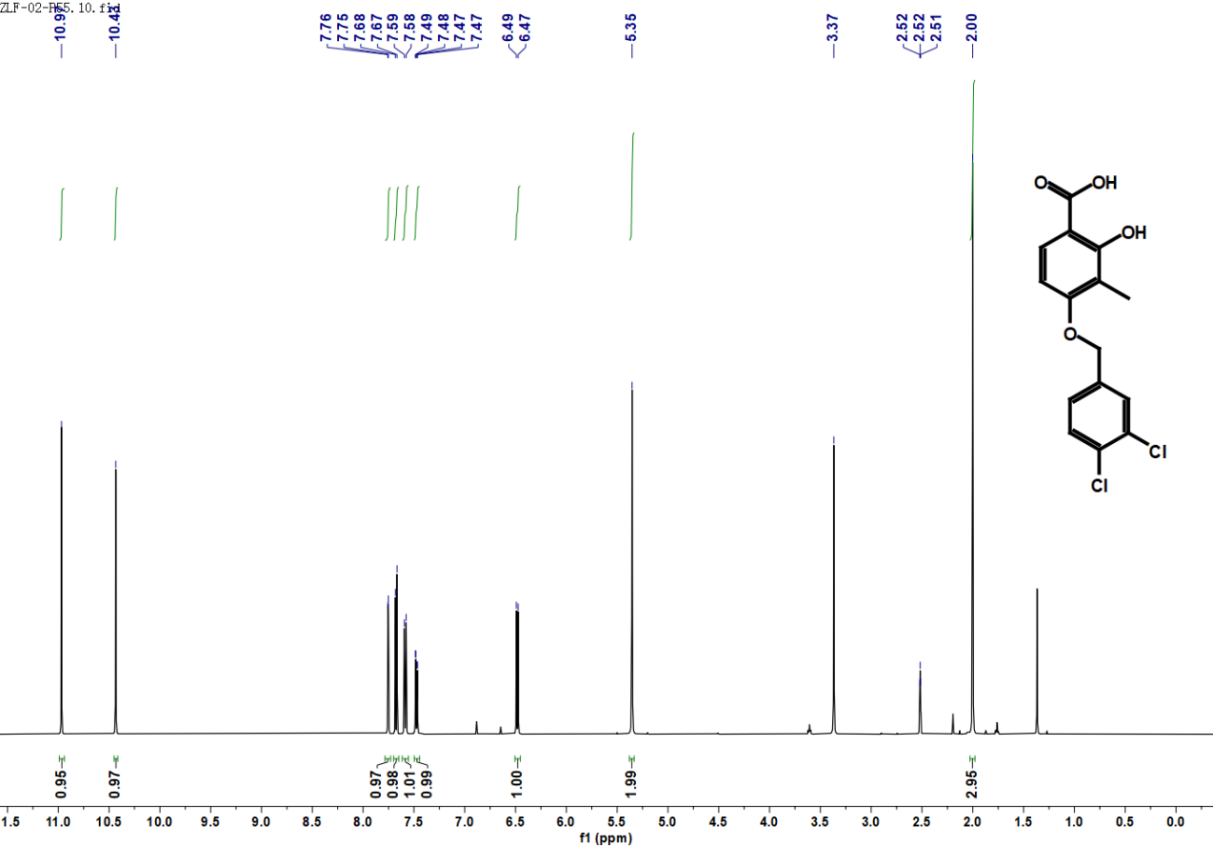

**Figure S26.** ^1^H NMR spectrum of compound **D6**.

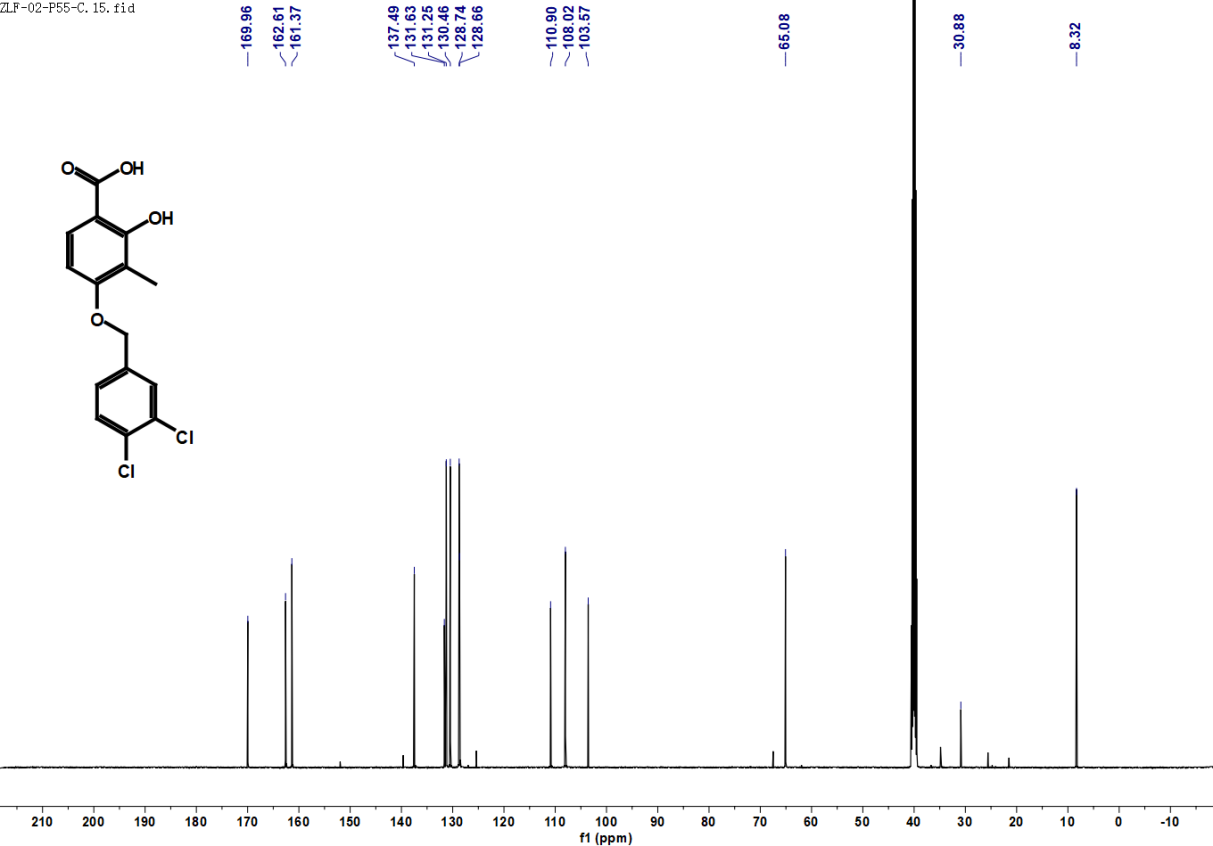

**Figure S27.** ^13^C NMR spectrum of compound **D6**.
